# Divergent Representation and Processing of Task Cues in Sensory and Prefrontal Cortices of Preterm-Born Mice

**DOI:** 10.1101/2024.11.26.625455

**Authors:** Emily McCoy, Vyshnavi Pendala, Mona Fariborzi, Lara Y. Demir, Olivia P. Buell, Samuel C. Fedde, Jacqueline B. Stinger, Luciano Elbaum, Troy D. Holsworth, Phillip Amenyo-Awude, Xin Tong, Adema Ribic

**Affiliations:** Program in Fundamental Neuroscience; Interdisciplinary Fellowship in Quantitative Neurobiology of Behavior; Interdisciplinary Fellowship in Reintegrating the Phenotype; Department of Psychology, College and Graduate School of Arts and Sciences, University of Virginia, Charlottesville, VA 22904

**Keywords:** preterm, prefrontal, visual, task, representation, processing

## Abstract

Preterm birth is a leading risk factor for atypicalities in cognitive and sensory processing, but it is unclear how prematurity impacts circuits that support these functions. To address this, we trained adult male and female mice born a day early (preterm mice) on a visual discrimination task and found that they fail to achieve high levels of performance due to increased responding to the non-rewarded cue (false alarms). While the representation of task cues measured with *in vivo* electrophysiology is intact in the primary visual cortex (V1) of trained preterm mice, the representation of the non-rewarded cue is significantly weaker in regular spiking, putative pyramidal neurons in the prefrontal cortex (PFC), a brain area that mediates response inhibition. Responses to both task cues are blunted in electrophysiologically and optogenetically identified fast-spiking Parvalbumin interneurons in preterm mice, indicating impaired processing of task cues in their PFC. Indeed, single trial neuronal responses evoked by the non-rewarded cue predict the behavioral outcome more accurately in term than in preterm mice. Similar cue representation and processing is present in the PFC of adolescent term-born mice, suggesting that preterm birth impedes prefrontal maturation. Surprisingly, environmental enrichment, a well-established paradigm that promotes sensory maturation, fails to improve the performance of preterm mice. Altogether, our study describes the long-term impact of preterm birth on prefrontal and visual circuits and suggests a limited capacity of early interventions for reducing the risk of cognitive deficits after preterm birth.

## Introduction

Children born prematurely (<37 weeks of gestation) are at 4-5x higher risk for developing intellectual disability (ID) and attention-deficit/hyperactivity disorder (ADHD) ^1–3^, at 10x higher risk for developing autism spectrum disorder (ASD) ^4,5^, and display persistent deficits in executive function and academic performance ^6–8^. As preterm labor occurs in >10% of all births in US and worldwide, preterm birth is the leading risk factor for cognitive and neurodevelopmental conditions ^6,9–11^. Yet, the neural circuits whose function may be disrupted by preterm birth are still unknown.

Impairments in the structure and function of the frontal cortex are frequently associated with executive dysfunction and neurodevelopmental conditions with high incidence in the preterm population ^9,12^. Unsurprisingly, neuroimaging and neurophysiological studies in preterm infants and children revealed reduced volume of and connectivity between frontal, thalamic and sensory areas ^11,13–16^. On a cellular level, studies of postmortem prefrontal cortex samples obtained from preterm fetuses demonstrated reduced density of inhibitory interneurons and altered expression of genes related to inhibitory neurotransmission ^17,18^. Some of the changes in brain structure can be recapitulated in animal models of prematurity, which also display disrupted layer-specific distribution and proportion of cortical interneurons ^18–21^, and alterations in the expression of multiple neurotransmission markers ^22,23^. Behaviorally, animal models of prematurity often display persistent hyperactivity ^23–26^, and some cognitive deficits ^18,27^, but the association between preterm birth-related changes in the frontal cortex and high incidence of cognitive dysfunction in the preterm population remains unclear.

To address this, we performed a detailed neurophysiological study of prefrontal function in adult mice born prematurely that were trained on a prefrontal cortex (PFC)-dependent visual discrimination task. While adult mice born 1 day early do not display any changes in baseline visual processing and locomotor activity, they show a significantly impaired learning trajectory. Preterm mice respond more often than term mice to the non-rewarded cue, have weaker representation of the non-rewarded cue in the PFC, and show disinhibition of neuronal firing during task performance. Cue representation in regular-spiking (RS) and evoked activity of fast-spiking (FS) neurons in the adult preterm PFC resemble those of adolescent term born mice, suggesting that preterm birth impairs prefrontal maturation. Surprisingly, postnatal environmental enrichment-a classical paradigm for improving the outcomes of impaired neurodevelopment ^21,28–32–fails^ to improve the task performance in preterm mice, indicating limited efficacy of early interventions in improving negative neurodevelopmental outcomes after preterm birth ^33^. Together, our study describes circuit- and cell-type specific impairments in prefrontal function and top-down sensory processing after preterm birth, paving a path towards uncovering the circuit mechanisms of cognitive dysfunction in the preterm born population.

## Methods

### Mice

Mice were maintained on C57BL/6 background (The Jackson Laboratory, Bar Harbor, ME) on reverse 12:12 light:dark cycle (lights off 11 AM-11 PM), with food and water *ad libitum,* except during visual discrimination training (see below). Animals of both sexes were used between postnatal day 21 and 7 months of age. Preterm mice were generated through timed breedings, where the day after the pairing was considered as gestational day (GD) 0. Once the pregnancy was confirmed (>1.5g increase in weight at GD 10), pregnant dams were habituated to handlers by daily handling sessions. Mifepristone (MFP, Milipore Sigma, Burlington, MA or HelloBio, Bristol, UK) was dissolved in DMSO (Milipore Sigma) and 150 µg was injected subcutaneously on GD 17. Preterm mice were delivered on GD 18. The cage with preterm mice was occasionally supplemented with external heat and oxygen to prevent hypothermia and hypoxia, commonly observed in preterm mice. Control term mice were obtained from timed pregnant dams injected with DMSO on GD 17, and with MFP on GD 18. All dams received continuous enrichment in form of plastic igloos and nesting material, as well as sunflower seeds (Bio-Serv, Flemington, NJ) beginning 3-4 days before parturition. Animals were treated in accordance with the University of Virginia Institutional Animal Care and Use Committee guidelines.

### Environmental enrichment

Nursing dams with their litters were transferred to a standard rat cage when the pups reached postnatal day (P) 5. Dams were co-housed with a nursing dam and its litter that was age matched and on BALB/c strain (purchased from Jackson Laboratory) to facilitate social enrichment. Cages contained multiple enrichment objects: InnoDome and InnoWheels from BioServ, gnawing sticks, multiple nesting pads, shredded paper and hay for building nests, as well as a variety of chew toys (pumice chews, willow twigs, willow balls, veggie chews, and hay tunnels, hideaways and sticks). Large enrichment objects were rearranged in the cage every 2 days, while chew toys were replaced with new ones every 2-3 days. Once the litters reached P25, they were weaned and group-housed in standard cages at 2-3 mice/cage to allow for placement of an InnoDome and InnoWheel, as well as 2-3 small chew toys. When the mice reached P60, they were prepared for visual discrimination task as described below.

### Viruses

Following viruses were purchased from Addgene: AAV-S5E2-ChR2-mCherry (135634-AAV1; 2x10^13^ vg/ml), AAV-EF1a-double floxed-hChR2(H134R)-mCherry-WPRE-HGHpA (20297-AAV9; 3.2x10^13^ vg/ml) and AAV-hSyn-HI-eGFP-Cre-WPRE-SV40 (105540-AAVrg; 1.7x10^13^ vg/ml). All viruses were diluted 1:10 in sterile PBS before the injection.

### Surgeries

Animals were anesthetized with isoflurane in oxygen (2-2.5% induction, 1–1.5% maintenance), warmed with a heating pad at 38°C and given subcutaneous injections of Buprenorphine SR (1 mg/kg) or Rimadyl (5 mg/kg) and 0.25% Bupivacaine (beneath their scalp). Eyes were covered with Puralube (Decra, Northwich, UK). Scalp and fascia from Bregma to behind lambda were removed, and the skull was cleaned, dried and covered with a thin layer of Scotchbond adhesive (3M, Maplewood, MN). Skin edges were sealed with VetBond (3M).

For virus injections, the head was immobilized in a custom-made stereotactic apparatus and small craniotomies were made on the left hemisphere with a dental drill at the following coordinates: primary visual cortex (V1): 0.5 mm anterior to lambda, 2-3 mm lateral to midline; anterior cingulate cortex (ACC): 2-2.5 mm anterior to bregma, 0.2-0.5 mm lateral to midline. NanoFil syringe (35 G beveled needle; WPI, Sarasota, FL) was then used to inject the virus at 1 nl/s using a Syringe Pump (KD Scientific, Holliston, MA). Volume of injected virus was 300 nl, injected in 2 150 nl increments at 2 different depths: 500 and 250 µm beneath the brain surface for V1 (targeting layers 2/3 and 5), and 700 and 800 µm for the PFC. The syringe was left in place for 5-10 minutes after the injection to allow the virus solution to diffuse and then very slowly raised to the surface to prevent the backflow. Muscimol (TMR-X conjugate; HelloBio, Princeton, NJ) was injected in the same manner, volume and at the same PFC coordinates of trained term mice at 1.6 mM concentration ^34^.

Head plates were attached after the Scotchbond adhesive application for animals that received no virus injection, or immediately after the virus injection for those that did. The head plate (stainless steel, SendCutSend, Reno, NV) was attached with dental cement (RelyX Ultimate, 3M). After the cement was cured, the well of the head plate was filled with silicone elastomer (Reynold Advanced Materials, Brighton, MA) to protect the skull.

If the mice received Rimadyl for analgesia, they were given Rimadyl dissolved in hydrogel (1% food-grade agar in distilled water) during the recovery ^35^. Animals were group-housed after the implantation and monitored daily for signs of shock or infection. On the day of electrophysiological recording, the animals were anesthetized as above and craniotomies (∼0.5 mm in diameter) were made above V1, PFC and cerebellum with 18G needles. If mice were injected with viruses, the existing craniotomies were reopened. The brain surface was covered in 2-3% low melting point agarose (Promega, Madison, WI) in sterile saline and then capped with silicone elastomer. Animals were allowed to recover for 1.5-2 h before the recording.

### Electrophysiology and optogenetics

For the recording sessions, mice were placed in the head-plate holder above an aluminum mesh treadmill and allowed to habituate for 5-10 minutes. The silicone plug was removed, the reference (insulated silver wire electrode; A-M Systems, Carlsborg, WA) was placed in cerebellum and the well was covered with warm sterile saline. A multisite electrode spanning all cortical layers (A1x16-5mm-50-177-A16; Neuronexus Technologies, Ann Arbor, MI) was coated with DiI (Invitrogen) to allow post hoc insertion site verification and then inserted in the V1 through the craniotomy. For PFC, a 4x4 shank probe (A4x4-3mm-50-125-177-A16) was used for the recordings, coated with DiI, DiO or DiD based on the type of fluorophore used in the experiment (mCherry or EGFP; listed in Viruses). The electrodes were slowly (5-10 µm/s) lowered to the appropriate depth: 800 µm for the PFC and until the uppermost recording site had entered the brain for the V1 and allowed to settle for 15-30 minutes. For optogenetics, a 473 nm laser diode with 0.4 µm fiber tip (0.22 NA, Doric Lenses, Quebec, Canada) was lowered to the craniotomy. 10 ms square pulses were delivered at 3-5 mW/mm^2^ (200 pulses at 0.5 Hz) using Spike2 (CED, Cambridge, UK) after the last recording session for optogenetic identification of neuronal subtypes (optotagging). The well was filled with 3% agarose prior to the recordings to stabilize the electrode and the whole region was kept moist with surgical gelfoam soaked in sterile saline (Pfizer, MA). The signals from the recording probes were fed into a 16-channel amplifier (Model 3500; A-M Systems, Sequim, WA) and amplified 200x, before being sampled at 25 kHz using Spike2 and Power 1401-3 data acquisition unit (CED). After the recording ended, the electrodes were slowly retracted, and the well was cleaned and protected with silicone elastomer.

### Visual stimuli

For visual stimuli, blank screen was generated with MATLAB (MathWorks, Natick, MA) using the PsychToolBox extension (Brainard, 1997) and presented on a gamma corrected 27” LCD. The screen was centered 20-25 cm from the mouse’s right eye, covering ∼80° of visual space. For calculating baseline orientation selectivity, sinusoidal gratings at 6 orientations 30° apart were presented at 100% contrast at 0.15 cpd, with 1 s long stimuli and 1 s interstimulus interval (blank screen). For visual discrimination training, visual stimuli (120° and 60° gratings) were presented at 0.15 cpd and 100% contrast in a random sequence for 3 seconds, followed by a 15 second long interstimulus interval (blank screen). Presentation of 120° triggered the delivery of 10 µl of water through a syringe pump (New Era Pump Systems, Farmingdale, NY), while the presentation of 60° had no consequence.

### Behavior

4-7 days after the headpost implantation, mice were gradually water restricted (from 3 ml of water/day to 1 ml of water/day) for 5-7 days. Food was available *ad libitum*. Mice were weighed daily and their weight was maintained at 85% of initial weight to prevent dehydration. During water restriction, mice were gradually habituated to handling by the experimenters, the treadmill, and the water delivery spout during daily habituation sessions ^36^. After water restriction, the visual discrimination training began, where mice were headfixed above the treadmill and a stainless-steel gavage needle (18G) was positioned near the mouse’s mouth for water delivery. The pump was controlled by the Power1401-3 data acquisition interface (CED) for lick detection ^37^. Mice were trained in 2 daily sessions consisting of 50 120° and 60° presentations each, for a total of 200 trials per day. Licks of the spout during the 10 s after the onset of stimulus presentation were quantified according to the signal detection theory as Hits (the fraction of trials in which the mouse licked to the 120° orientation), Misses (the fraction of trials in which the mouse failed to lick to the 120° orientation), False Alarms (the fraction of trials in which the mouse licked to the 60° orientation) and Correct Rejections (the fraction of trials in which the mouse failed to lick to the 60° orientation), where the performance was measured as discriminability (d’)=z(H) - z(FA). Mice were trained until they achieved a d’ of 2 or above for 3 consecutive sessions or up to 4000 trials if they failed to reach the criterion.

For open field, mice were placed in a 24x24 inch plexiglass arena after a brief habituation to the room and the experimenters. Their activity was recorded for 10 minutes and analyzed using an open-source toolbox ^38^.

For measuring locomotor behavior during task performance, a small piezoelectric disc was placed in contact with the treadmill and the signal was digitized using Power 1401-3 and Spike2. Waveform averages in response to both cues were constructed using Spike2 and used to compare the amount of treadmill movement during the training.

Animals were monitored for signs of distress for the duration of the experiment. Animals that developed corneal damage, cataracts, and symptoms that had the potential to impact their behavior or physiology (such as immobility and other signs of pain, sudden weight loss or impeded weight gain) were excluded from the study. All mice except those in the MFP Term and Environmental Enrichment groups underwent electrophysiology either at the beginning or at the end of the training, or both. If the animals were recorded at the onset of training, that was counted as the first training session.

### Histology and imaging

After the last training session or electrophysiological recording, mice were anesthetized with a mixture of ketamine and xylazine and transcranially perfused with warm 0.1 M phosphate buffer, followed by warm 4% paraformaldehyde (Electron Microscopy Sciences, Hatfield, PA). Brains were postfixed 1 hr at room temperature, followed by overnight fixation at 4°C. Brains were sectioned into 40-80 µm sections using a vibratome and stored in 1x phosphate buffered saline (PBS) and 0.01% sodium azide. For immunohistochemistry, sections were rinsed in PBS, non-specific binding was blocked with 3% normal horse serum (heat inactivated, ThermoFisher, Waltham, MA) and 0.3% Triton-X 100 (Sigma-Aldrich) in PBS (sterile filtered). Antibodies were incubated overnight at 4°C. Parvalbumin (anti-goat) was used at 1/200 (Swant, Belinzona, Switzerland) and detected with donkey anti-goat Alexa 647 (ThermoFisher). After staining, sections were rinsed in distilled water, mounted on glass slides, briefly dried, and coverslipped with Aquamount (Polysciences, Warrington, PA). Images were acquired using Leica Stellaris 8 at 2048x2048 resolution using 10x HC PLAN FLUOTAR air (NA=0.3 for tiling) or 40x HC PL APO CS2 (NA=1.3) oil immersion objective.

### Electrophysiological and behavioral data analysis

Locomotion measured with piezoelectric sensors was analyzed in Spike2 (CED) using waveform averaging and cues as triggers. Spikes were isolated from the recordings using template matching in Spike2 (CED). Briefly, the recordings were band-pass filtered (0.7-7 kHz) and smoothed (1 ms). Threshold for detection was set at 3 SDs of the mean baseline (blank screen) and a window of 0.9-1 ms. Isolated units were clustered using waveform properties (amplitude, spike half-width and slope of repolarization), and the clusters were checked manually for quality in addition to confirming there were no spikes during the refractory period (2 ms) using interval histograms. Units with refractory period violations or units with large variations in amplitude (>10%) were excluded from analysis. For orientation selectivity, custom-written scripts in Spike2 were used to plot the baseline-subtracted responses of isolated units to the presented orientations (10 ms bin) and construct a tuning curve, which was then fitted with a Gaussian to determine the preferred (O_pref_) and orthogonal (O_orth_) orientations, and calculate the orientation selectivity index (OSI) as (R_pref_- R_orth_)/(R_pref_+R_orth_), where R is the firing rate at preferred and orthogonal orientations. We considered the neurons with at least 3x elevated firing rate to the preferred orientation (averaged over 0.8 s and compared to averaged 0.2 s of baseline) as visually responsive. Tuning width was calculated as half-width at half-maximum of the O_pref_.

For optotagging, spikes were isolated from optogenetic recording sessions and PSTHs (0.1-1 ms bin) were constructed in response to the laser pulses. Units with reliable responses (>50%) of trials and short latencies (within 10 ms of the laser pulse) were considered optotagged and their waveforms were used to identify their spiking during the task performance. Unit waveforms were further compared between optotagging and task recordings for quality assurance. Units whose waveforms differed by more than 10% in amplitude and width between the two sessions were discarded from analysis.

Selectivity indices (SIs) were calculated from 10 ms peristimulus time histogram (PSTHs) as (R_cue_-R_baseline_)/(R_cue_+R_baseline_), where R_cue_ is the firing rate during the initial 1-1.5s presentation of 120° and 60° stimuli. Behavioral responses during the task were analyzed using custom scripts in Spike2 by converting the detected licks into events and constructing PSTHs in response to the non-rewarded cue during the 2 possible outcomes (Correct Rejection and False Alarm).

Stationary and moving stages were not analyzed separately as animals were typically walking in short bouts during the recordings. Recordings where large motion artifacts were present when the animal was simply balancing on the treadmill or grooming were excluded from analysis, along with recordings from locations outside of the PFC or V1 (as determined through post-hoc mapping of insertion sites).

### Quantification and statistical analysis

All analyses were performed with the researchers blind to the condition. Statistical analyses were performed in GraphPad Prism 9.0 (GraphPad Inc., La Jolla, USA), R and JASP, as indicated in text and figure legends. N refers to the number of animals or single units, as indicated in figure legends or in text. For orientation selectivity analysis, OSIs, proportions of selective cells (OSI>0.3) and tuning width (OSI>0.3) means (normal data distribution, Shapiro-Wilk test) or medians (non-normal distribution) were calculated per mouse before comparison. For comparison of preferred orientations, selective neurons (OSI>0.3) were only segregated by group (Term/Preterm). For all other analyses, units significantly modulated by cues were determined through comparison of pre-cue and post-cue firing using Wilcoxon signed rank test. The proportions of significantly modulated units (positively and negatively) were determined and reported per animal, and PSTHs and SIs represent significantly modulated units. PSTHs were compared using permutation test of area under the curve (AUC) calculated from 0.5 s precue and 1 s (V1) or 3 s (PFC) postcue activity (10 ms bins). Cue representation was determined as (%R-%NR)/(%R+%NR), where %R and %NR are the proportion of significantly positively modulated units by the rewarded and non-rewarded cue, respectively. All unit data was tested for normality using Shapiro-Wilk test and data violating the assumption of normality was tested using non-parametric tests (Mann-Whitney t-test, Kolmogorov-Smirnoff for comparison of distributions or non-parametric ANOVA).

Normally distributed data was tested using t-tests and ANOVA with post-hoc tests for comparing groups as indicated in text and figure legends. Levene’s test was used to test homoscedasticity with appropriate corrections used for heteroscedasticity. For linear mixed model (LMM) in Figure 2e, an LMM with Animal Group as a fixed effect and Subject and Training Day as random effects was fitted to the data using the following formula and maximum likelihood method: d’’ ∼ Group + (1 + Group || Subject) + (1 + Group || Training Day). To test the sex effects, LMM with Sex and Animal Group as fixed effects and Subject and Training Day as random effects was fitted using the following formula and maximum likelihood method: d’’ ∼ Sex + Animal Group+ (1 + Sex + Animal Group | Subject) + (1 + Sex + Animal Group | Training Day). For Figure 2i, lick counts were averaged per 20 trials and an LMM with Animal Group and Trial Block as fixed effects and Subject as the random effect was fitted to the data using the following formula and maximum likelihood method: Lick count’ ∼ Group + ’Trial block’ + (1 + Group + ’Trial Block’ | Animal). For Figure 6b, an LMM with Age as a fixed effect and Subject and Training Day was fitted to the data using the following formula and maximum likelihood method: ’d’’ ∼ ’Agè + (1 + ’Agè | Training Day) + (1 + ’Agè | Animal). For Figure 7b, an LMM with Enrichment and Birth as fixed effects and Subject and Training Day as the random effects was fitted to the data using the following formula and maximum likelihood method: ’d’’ ∼ Enrichment + Birth + (1 + Enrichment + Birth | Subject) + (1 + Enrichment + Birth | Training Day). For linear regression, SI values from all neurons per animal were averaged and used as predictors of d’ as described in text and figure legends. For logistic regression, trial by trial spike time stamps were used to calculate the evoked rate per trial (number of spikes in 3 s window postcue-number of spikes in 3 s window precue). As behavioral outcomes were biased towards Correct Rejection, random oversampling was used to balance the datasets. Data are reported as mean ± SEM (standard error of mean) or median±MAD (median absolute deviation), as indicated in text or figure legends, where N represents the number of animals or the number of single units. Target power for all sample sizes was 0.8 and alpha was set to 0.05.

**Figure 1.**
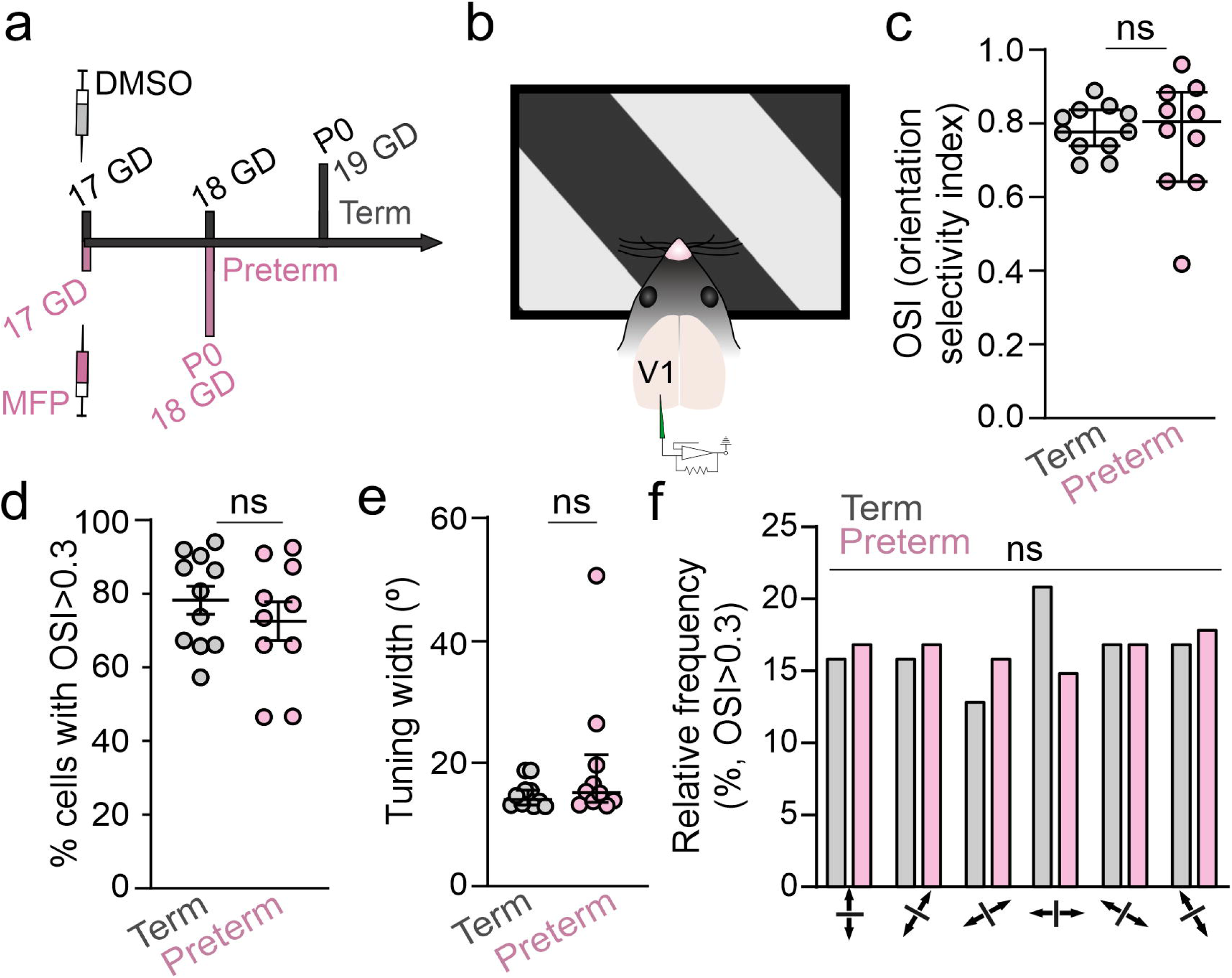
Adult preterm mice have intact orientation selectivity. **a)** Preterm mice were generated through subcutaneous injection of mifepristone (MFP) to timed pregnant dams at gestational day (GD) 17. Preterm mice were delivered at GD18, a day early. Control term mice were delivered at GD19 by dams injected with vehicle (DMSO) at GD17. **b)** Primary visual cortex (V1) of mice was recorded while mice were presented with 6 different orientations 30° apart. Preterm mice had no significant differences in **c)** orientation selectivity index (OSI median±MAD : Term=0.78±0.05, Preterm=0.81±0.08, Mann Whitney t-test p=0.86, U=52), **d)** fraction of neurons with OSI<0.3 (mean±SEM: Term=77.59±3.81, Preterm=72.04±5.22, t-test p=0.4, t=0.86, df=16.84), **e)** tuning width of selective neurons (median±MAD: Term=14.02±1.08, Preterm=15.19±1.73, Mann Whitney t-test p=0.29, U=70.5), or **f)** the distribution of preferred frequencies (p=0.92, X^2^ test). N=414 neurons from 11 term mice and 306 neurons from 10 preterm mice.

**Figure 2.**
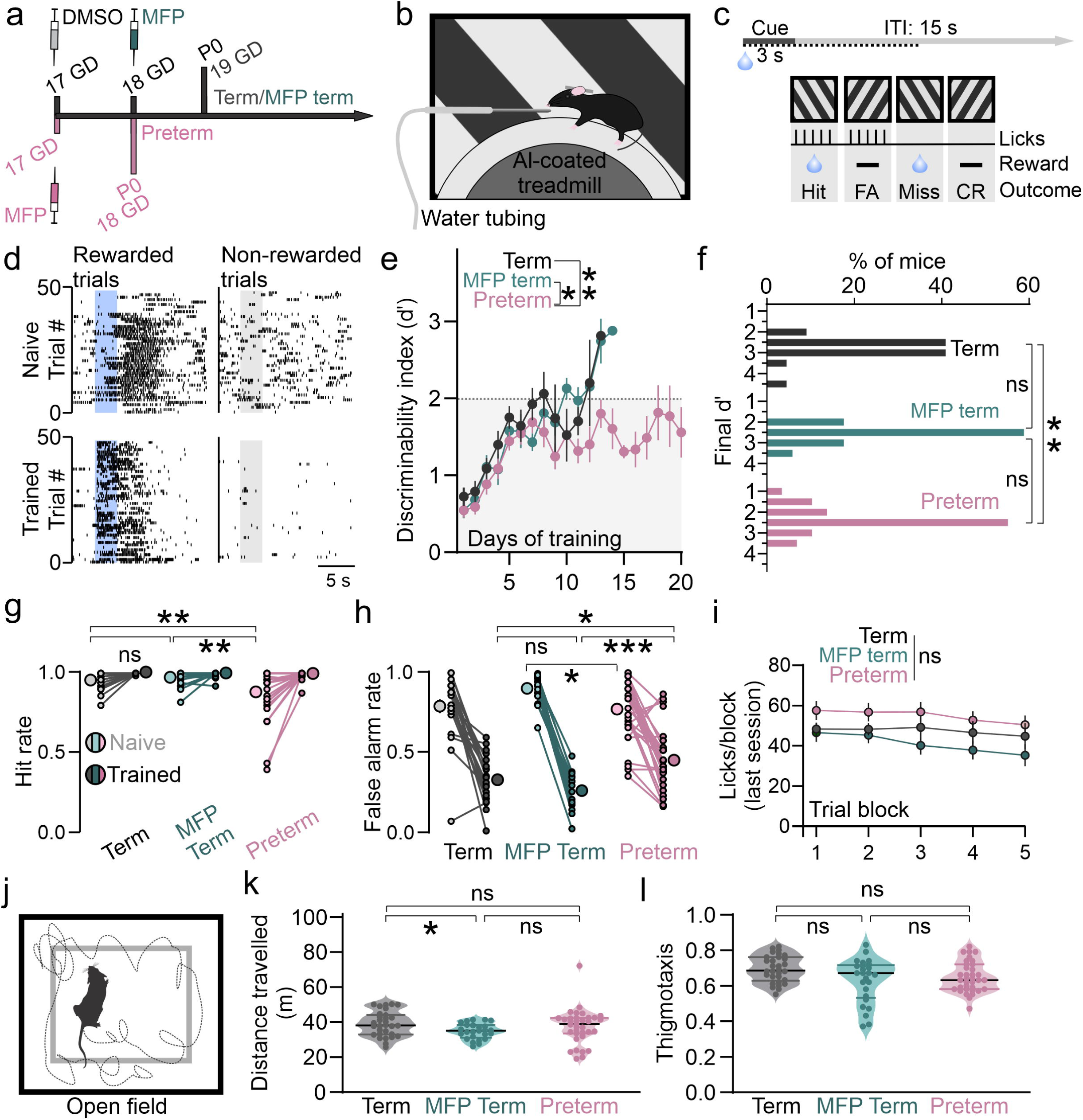
Preterm mice show impaired learning and behavioral inhibition in a visual discrimination task. **a)** Schematics of gestation length for experimental groups. Preterm mice were generated through subcutaneous injection of MFP to timed pregnant dams at GD17, and control term mice were delivered at GD19 by dams injected with vehicle (DMSO) at GD17 or with MFP at GD18. **b)** Head-fixed term and preterm mice were trained to discriminate two orientations while locomoting on a treadmill positioned in front of a screen displaying the task cues. The reward (water) was delivered through a spout positioned near the mouth. **c)** Task structure: a single session consisted of 50 randomized, 3 s long presentations of rewarded and non-rewarded cue each, with an intertrial interval of 15 s. The detection window was 10 s. Four possible outcomes are Hit, False Alarm (FA), Miss and Correct Rejection (CR), as indicated. **d)** Representative lick raster of naïve (top) and trained (bottom) mice to the rewarded (left) and non-rewarded (right) cues, indicated with a blue and grey rectangle, respectively. Note the reduced reaction times after the presentation of the rewarded cue and withholding of licks during and after the non-rewarded cue in trained mice. **e)** Behavioral performance was measured as d’, which showed linear improvements in both groups of term mice, but not in preterm mice (LMM, Term vs Preterm p=0.007, MFP Term vs Preterm p=0.02, Term vs MFP Term p=0.78; N=22 Term, 29 Preterm and 17 MFP Term mice). **f)** A significant fraction of preterm mice (17.24%) failed to reach the criterion, in contrast to term mice (Kolmogorov-Smirnov test of d’ values from the last training session Term vs Preterm: D=0.43, p=0.009, Term vs MFP Term: D=0.34, p=0.14, MFP Term vs Preterm: D=0.2893, p=0.23). **g)** Hit rates of naïve preterm mice are significantly lower at the onset of training (ANOVA birth x training p=0.016, F_(2, 65)_=4.37; Bonferroni post-hoc Naïve Term vs. Preterm p=0.005, Term vs MFP Term p>0.9, MFP Term vs Preterm p=0.0013; all p>0.9 among Trained groups). **h)** False Alarm rates were significantly higher in trained preterm mice, indicating impaired response inhibition (ANOVA birth x training p<0.0001, F_(2, 65)_=10.66; Bonferroni post-hoc Naïve Term vs. Preterm p>0.9, Term vs MFP Term p=0.13, MFP Term vs Preterm p=0.04; Trained Term vs Preterm p=0.036, Term vs MFP Term p=0.53, MFP Term vs Preterm p=0.0007). **i)** Comparison of lick frequencies (binned in 20 trial blocks) during the last session revealed a significant effect of time in session (LMM; Trial Block p<0.0001), with lick frequencies dropping towards the end of the session at the same level across groups (Animal Group p=0.15). **j)** Schematics of the open field test. Mice were released into an open arena and their locomotor activity was recorded and quantified. **k)** Term and preterm mice crossed a similar distance in the open field, confirming similar levels of locomotor activity (Welch’s ANOVA (F_(2,56.98)_=3.9, p=0.03; Dunnett’s T3 multiple comparisons test Term vs. MFP Term p=0.03, Term vs. Preterm p=0.83, MFP Term vs. Preterm p=0.41; N=30 Term, 25 MFP Term and 35 Preterm mice). **l)** Term and preterm mice had comparable preference for the walls of the open field arena, indicating comparable exploratory drive (Welch’s ANOVA F_(2,51.62)_=4.08, p=0.03; Dunnett’s T3 multiple comparisons test Term vs. MFP Term p=0.07, Term vs. Preterm p=0.06, MFP Term vs. Preterm p=0.87).

**Figure 3.**
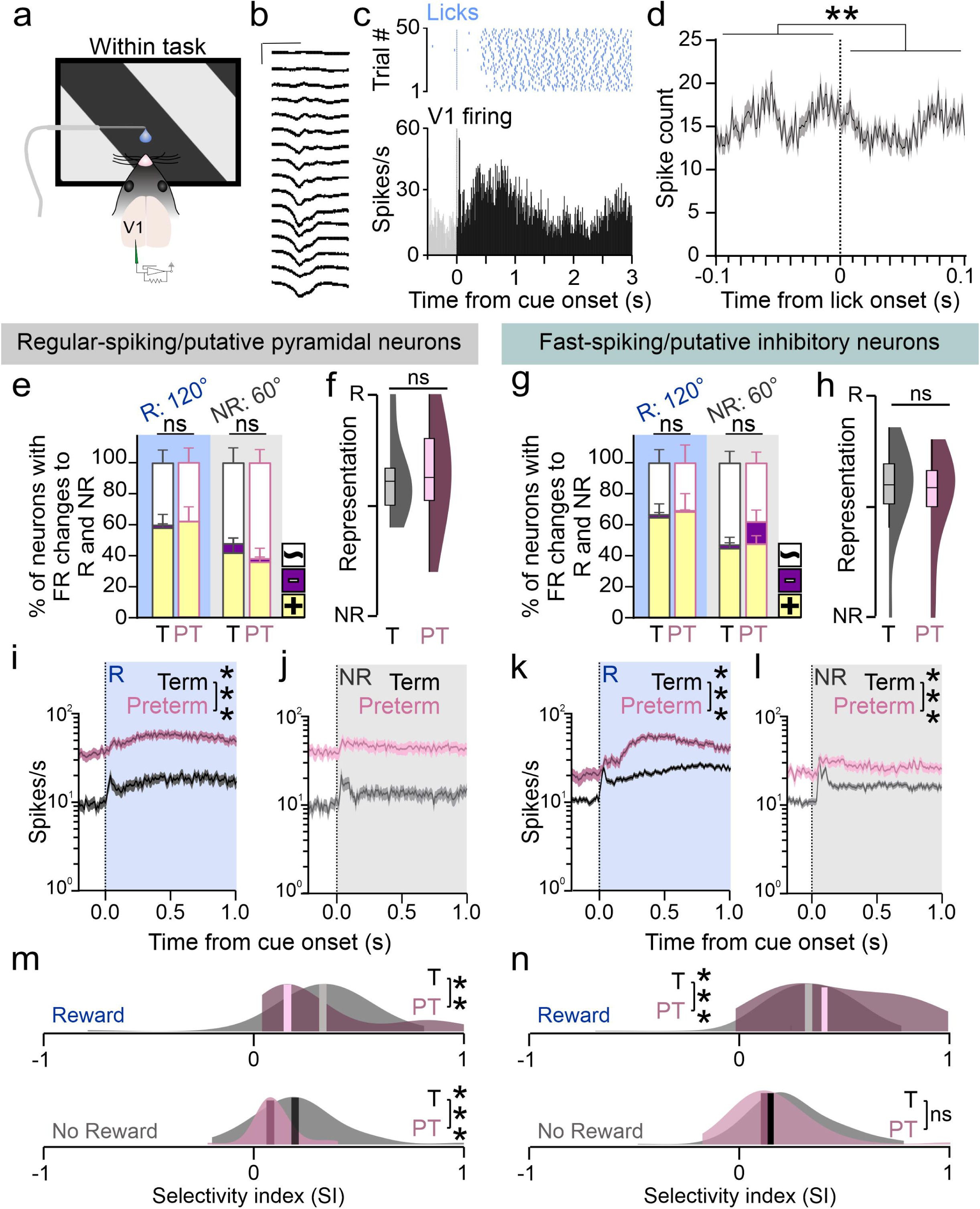
Processing of task cues is impaired in the V1 of trained preterm mice. **a)** V1 activity of trained term and preterm mice was recorded while the mice were engaged in the task. **b)** Representative example of averaged field potential response (visually evoked potential/VEP) to task cues used for confirming the recording location. Scale bars: 100 ms and 50 µV. **c)** Top: lick raster, and Bottom: peristimulus time histogram (PSTH) of a representative V1 neuron showing the transient and sustained components of the response during the cue presentation. **d)** Correlogram of V1 unit activity and licks revealed a peak in firing of V1 neurons prior to lick onset (Wilcoxon matched-pairs signed rank test, p=0.018, W=-1354). **e-h)** Representation of task cues is intact in regular-spiking (RS, e and f) and fast-spiking (FS, g and h) neurons in the V1 of preterm mice (RM ANOVA with birth and the direction of modulation as factors; RS Reward: birth x modulation p=0.93, F_(2,38)_=0.07; RS No Reward: birth x modulation p=0.7, F=0.36; f: 2-tailed t-test, p=0.9, t=0.0046, df=19; FS Reward: birth x modulation p=0.95, F=0.09; FS No Reward: birth x modulation p=0.24, F=1.46; h: p=0.66, t=0.45; bar graphs represent mean±SEM of % of positively modulated/yellow, negatively modulated/magenta and unmodulated/white neurons, box plots indicate median and quartiles). **i-l)** PSTHs of V1 neurons significantly modulated by task cues revealed profoundly elevated firing in both RS (i and j) and FS (k and l) V1 neurons in preterm mice (permutation tests of area under the PSTH curve, p=0.000 for all). Data are represented as mean±SEM**. m-n)** Distributions of selectivity indices (SI) of neurons significantly modulated by task cues revealed a significant shift to lower values for both cues in RS neurons in preterm mice (m) and an increase in higher values for the rewarded cue in FS neurons (n). K-S test for all: RS Reward D=0.35, p=0.0007; RS No Reward D=0.46, p=0.00005, FS Reward D=0.29, p=0.000006; No Reward D=0.11, p=0.36; Ns are indicated in main text).

**Figure 4.**
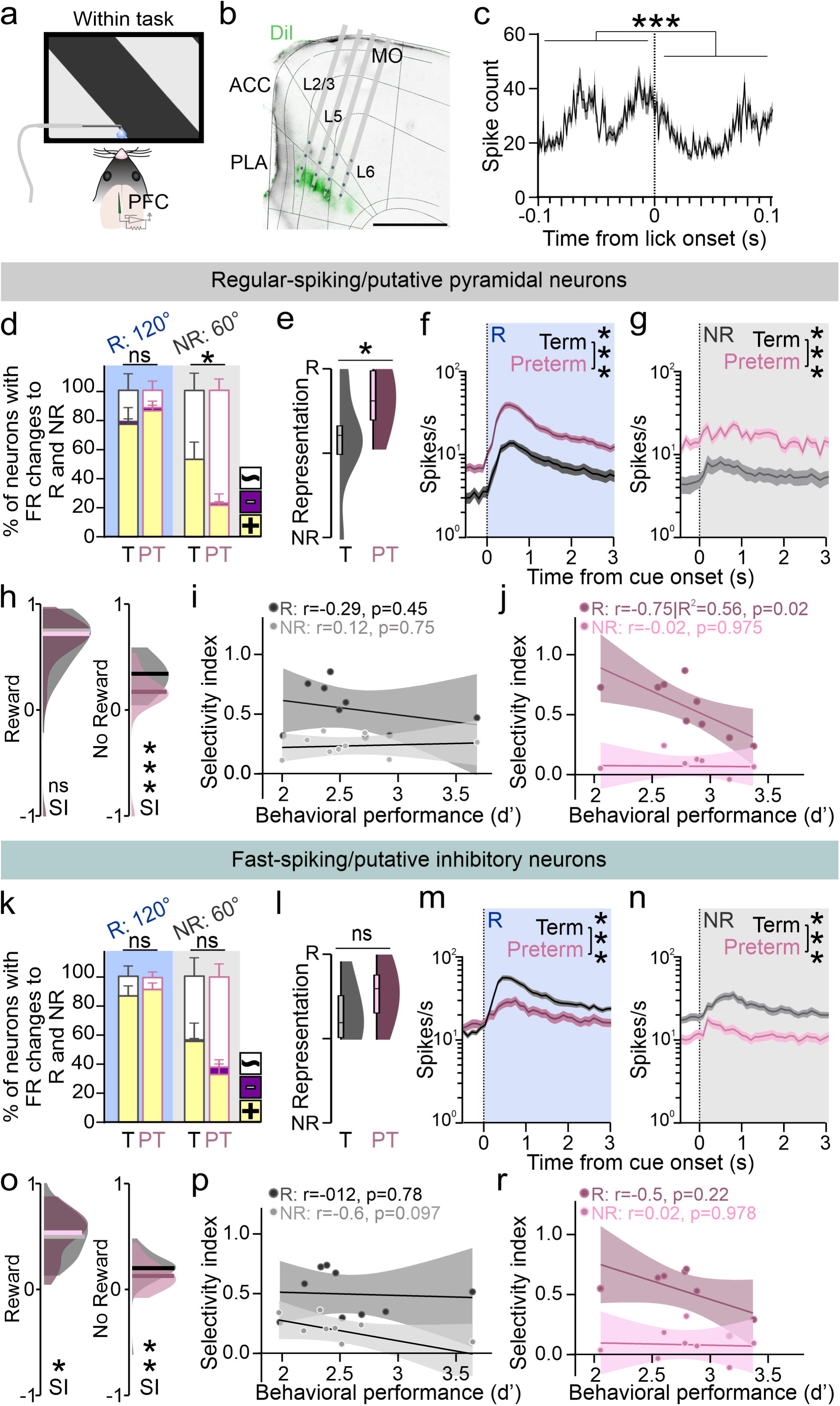
Representation and processing of task cues is impaired in the PFC of preterm mice. **a)** PFC activity of trained term and preterm mice was recorded while the mice were engaged in the task. **b)** Representative electrode tracks in the coronal section of the PFC. DiI (green) marks the tracks, PFC areas and layers are outlined (Allen Brain Atlas). Scale bar: 500 µm. **c)** Correlogram of PFC unit activity and licks revealed peaks in firing of PFC neurons prior to lick onset (Wilcoxon matched-pairs signed rank test, p<0.0001, W=-2770). **d)** Representation of the non-rewarded cue is significantly weakened in preterm mice (ANOVA Reward: birth x modulation, p=0.6, F_(2,34)_=0.52; No Reward: p=0.014, F=4.88. Data represents mean±SEM of % of positively modulated/yellow, negatively modulated/magenta and unmodulated/white neurons of 9 mice per group. **e)** Overall cue representation is shifted in favor of the rewarded cue in preterm mice (Mann-Whitney test p=0.045, U=20.5; medians and interquartile range are indicated in box plots). **f-g)** PSTHs of RS neurons significantly modulated by task cues revealed significant hyperactivity in preterm PFC. Permutation test of PSTH AUCs, p=0.000, Ns are indicated in main text. Data represents mean±SEM. **h)** Distributions of SIs indicated that RS neurons in preterm mice are significantly less responsive to the non-rewarded cue, with several neurons completely suppressed (values close to -1). Reward: K-S test, D=0.13, p=0.36; No Reward: D=0.4, p=0.000014; medians are indicated. **i)** Linear regression of SI values for term and **(j)** preterm mice revealed that the SI for the rewarded cue are correlated with and predictive of performance (d’) in preterm but not term mice (Pearson’s correlation and linear regression; r values, 95% confidence intervals and regression are shown in graph). **k-l)** Representation of cues is not impaired in FS neurons of preterm PFC (Reward: birth x modulation, p=0.71, F(2,34)=0.34; No Reward: p=0.2, F=1.67; Mann-Whitney test p=0.17, U=28). m-n) PSTHs of FS neurons significantly modulated by task cues revealed significantly blunted cue-evoked activity in preterm PFC. Permutation test of PSTH AUCs, p=0.000, Ns are indicated in main text. **o)** SI values of preterm mice were shifted towards higher values for the rewarded cue and towards lower for the non-rewarded cue (K-S test, Reward D=0.19, p=0.02; No Reward D=0.36, p=0.00097). **p-r)** SI values of FS neurons in either group of mice did not correlate with d’ (Pearson’s correlation, r values indicated in graph).

**Figure 5.**
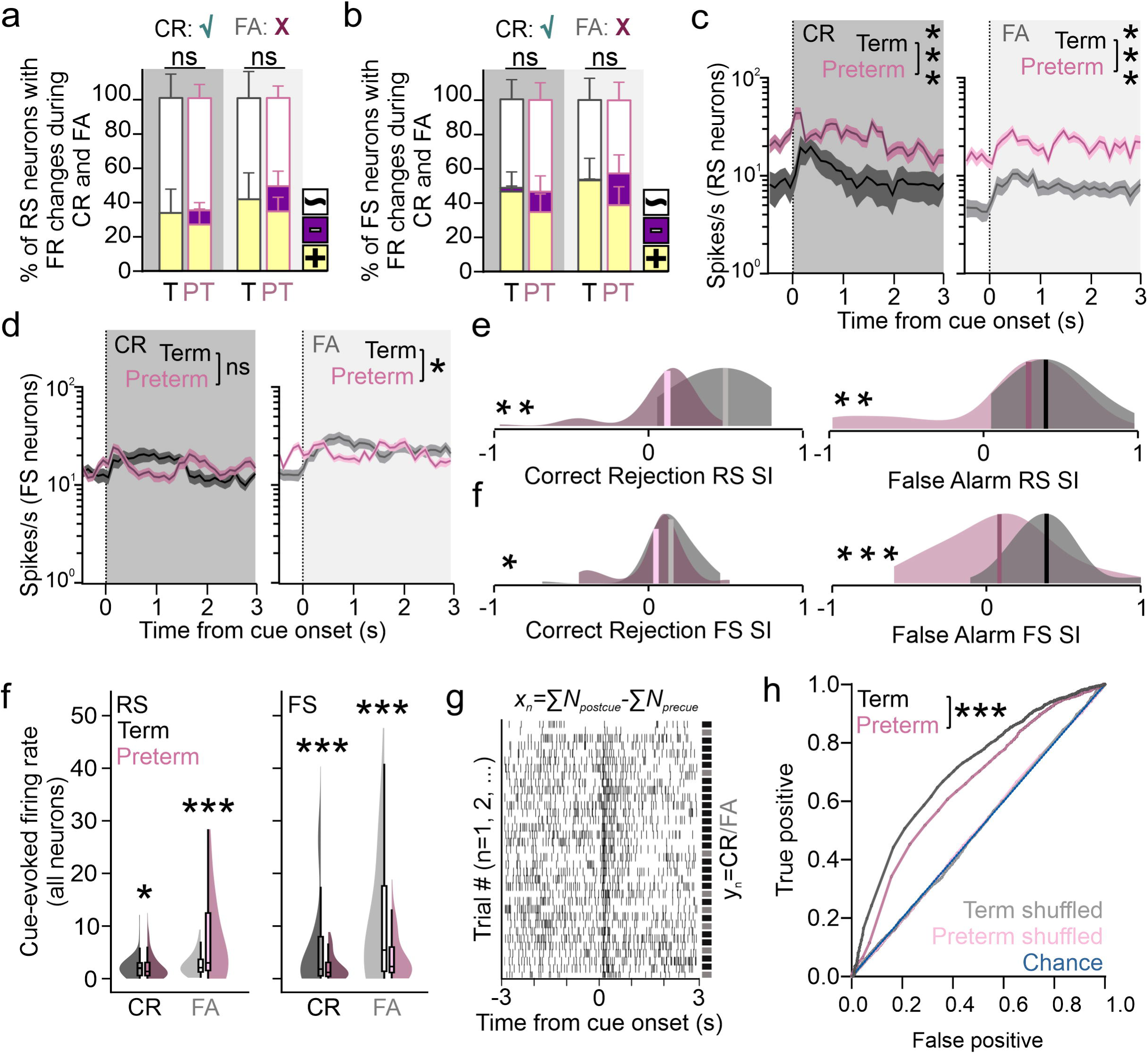
Encoding but not representation of behavioral outcomes to the non-rewarded cues is altered in the PFC of preterm mice. a-b) Representation of behavioral outcomes to the non-rewarded cue is not altered in prefrontal RS (a) or FS (b) neurons of preterm mice (ANOVA: birth x modulation RS CR: p=0.81, F_(2,34)_=0.21, FA: p=0.6, F=0.51; FS CR: p=0.65, F=0.43, FA: p=0.45, F=0.82). Data represents mean±SEM of % of positively modulated/yellow, negatively modulated/magenta and unmodulated/white neurons of 9 mice per group. **c)** PSTHs of RS neurons significantly modulated by the non-rewarded cue during CR (left) and FA (right) revealed elevated firing of RS neurons during both behavioral outcomes in preterm mice (permutation test of PSTH AUCs, p=0.000; Ns are indicated in main text). **d)** PSTHs of FS neurons significantly modulated by the non-rewarded cue during CR (left) and FA (right) revealed blunted and irregular firing of FS neurons during both behavioral outcomes in preterm mice that was significantly different to term mice only during FA trials (permutation test of PSTH AUCs, CR p=0.125, FA p=0.022; Ns are indicated in main text). **e)** SI values for both RS and FS neurons were shifted towards lower values during both behavioral outcomes (K-S test; RS CR D = 0.44, p=0.000042; FA D = 0.26, p=0.014; FS CR D=0.24, p=0.044; FA D=0.57, p=1.2x10^-10^). **f)** Non-rewarded cue-evoked firing of all neurons isolated from term and preterm mice during CR and FA revealed overall significant differences in evoked activity in preterm PFC, with a pronounced reduction of cue-evoked firing of FS neurons (K-S test; RS CR p=0.011, D=0.23; FA, p=0.0008, D=0.28; FS CR p=0.0001, D=0.26; FA, p=0.93x10^-6^, D=0.31). **g)** Logistic regression model was fit with trial-by-trial firing rates evoked by the non-rewarded cue as classifiers and CR/FA as outcomes, where N=number of spikes in stimulus/baseline window. **h)** ROC curves indicated that firing rate data of term mice performs significantly better in the model than the firing rates of preterm mice (z=6.79, p<0.0001).

**Figure 6.**
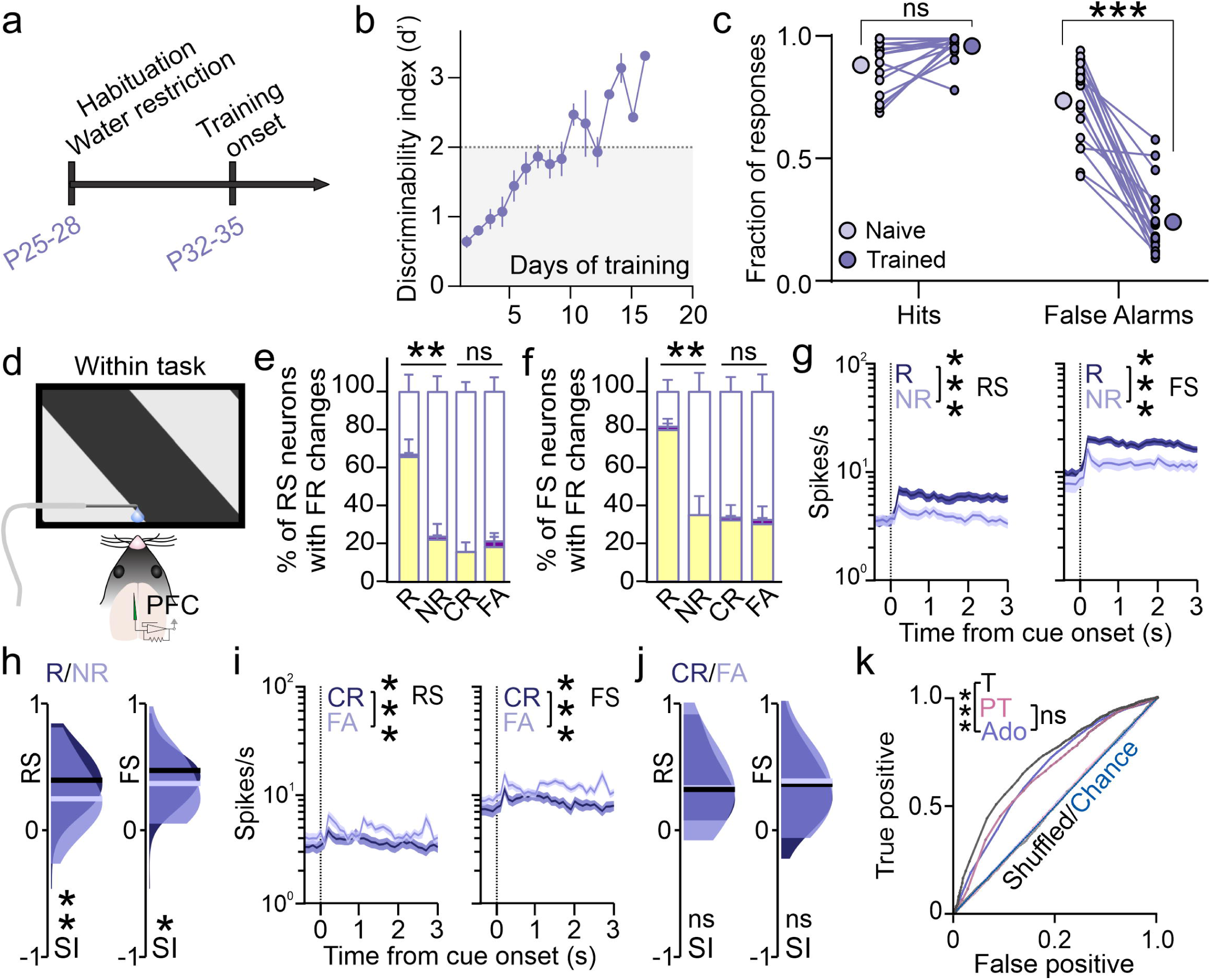
Non-rewarded cue has a significantly reduced representation in the PFC of adolescent term mice. **a)** Experimental timeline: mice were prepared for training starting at postnatal day 25, and the training began one week later. **b)** Adolescent term mice learn the task comparable to adult term mice (LMM, age p=0.42; data represents mean±SEM of N=18 male and female mice, details indicated in main text). **c)** Increase in the performance of adolescent mice is due to a reduction in False Alarm rate (RM ANOVA: response x training p<0.0001, F_(1,17)_=40.5; HR p=0.07, FA p<0.0001). **d)** Schematic of electrophysiological recordings used to collect the data. **e-f)** Representation of the non-rewarded cue is significantly weaker than that of the rewarded cue in both RS (e; RM ANOVA cue x modulation p<0.0001, F_(2.2,17.79)_=37.11) and FS (f; p<0.0001, F_(2.5,20.04)_=18.37) neurons. Data represents mean±SEM of % of positively modulated/yellow, negatively modulated/magenta and unmodulated/white neurons of 9 mice per group. **g)** PSTHs of significantly modulated neurons revealed weaker activity evoked by the non-rewarded cue in both RS (left) and FS neurons (right; p=0.000, permutation test of PSTH AUCs). Data represents mean±SEM. **h)** SI indices of neurons in (g) confirmed that the selectivity for the non-rewarded cue is shifted towards lower values in both RS and FS neurons (K-S test RS: D=0.31, p=0.005; FS: D=0.27, p=0.01). Medians are indicated. **i)** PSTHs of neurons significantly modulated by the non-rewarded cue showed a significantly higher activity during FA trials in both RS (left) and FS (right) neurons of adolescent PFC (p=0.000, permutation test of PSTH AUCs). Data represents mean±SEM. **j)** Distribution of SI values during CR and FA trials was not significantly different in RS or FS neurons (K-S test; RS: D=0.21, FS: D=0.12, p=0.89). Medians are indicated. **k)** Logistic regression model (as in Figure 5g-h) of single trial firing rates evoked by the non-rewarded cue as classifiers of CR/FA revealed that the ROCs of adolescent term and adolescent preterm mice substantially overlap (adolescent vs preterm p=0.17, z=1.38; adolescent vs term adult p<0.0001, z=-5.76). All Ns are indicated in the main text.

**Figure 7.**
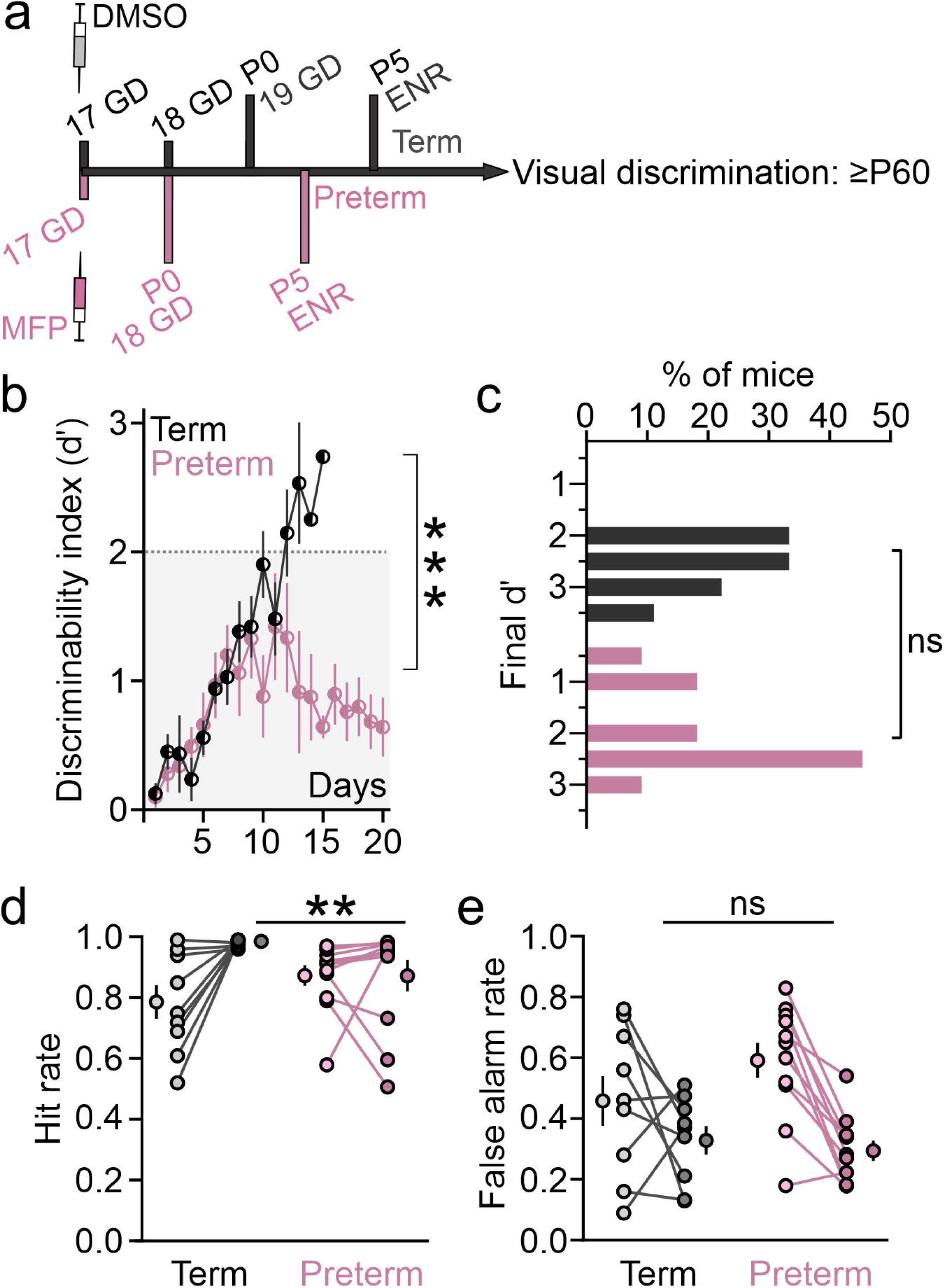
Lifelong environmental enrichment fails to improve the learning trajectory of preterm mice. **a)** Experimental timeline. Term and preterm mice were housed in an enriched environment from postnatal day 5 (P5) until the end of the training. **b)** The learning trajectory of term mice reared in an enriched environment was steep, but they performed worse than term mice from the standard environment. Preterm mice reared in an enriched environment also performed worse than preterm mice from standard environment [LMM with Enrichment (ENR) and Birth as Fixed Effects; ENR p<0.0001, Birth p<0.0001) **c)** There are no differences in the distribution of final d’ values between term and preterm mie reared in an enriched environment (K-S test, D=0.46, p=0.16**). d)** Hit rates during the final training session were significantly different between term and preterm mice reared in enriched environment (ANOVA training x birth x enrichment p=0.002, F=28.49; naïve ENR Term vs Preterm p=0.19, trained p=0.006. **e)** False Alarm rates were not significantly different between term and preterm mice reared in enriched environment (naïve ENR Term vs Preterm p=0.99, trained p=0.99; N=9 term and 11 preterm mice, statistical details are indicated in Supplementary Materials).

## Results

### Adult preterm mice commit more errors during visual discrimination

Children born preterm have an increased rate of cognitive deficits and neurodevelopmental conditions with unclear circuit origins ^9^. As we previously found that preterm birth similarly impacts the resting neural activity in the primary visual cortex (V1) of juvenile mice and infants ^39^, we hypothesized that mice born preterm would also display cognitive deficits. To test this, we chose to train the mice in a visual discrimination task, a classical learning task that facilitates electrophysiological probing of neural circuits in awake, behaving mice ^40,41^. Preterm mice were generated through subcutaneous injections of mifepristone (MFP) to timed-pregnant C57BL/6 dams on gestational day (GD) 17 (Figure 1A), resulting in birth of viable preterm pups on GD 18. Term pups in our colony are born on GD 19 with very little variability, as previously published ^42,43^. Before training, adult (2-7 months old) male and female term and preterm mice were implanted with headposts for head fixation and their baseline visual function in form of orientation selectivity was tested using *in vivo* electrophysiology ^44^. We collected neuronal responses to sinusoidal oriented gratings from all cortical layers (6 orientations 30° apart), at 100% contrast and 0.15 cycles per degree spatial frequency; Figures 1B), and determined the orientation selectivity index (OSI; Figure 1C), the number of orientation selective neurons (Figure 1D), orientation tuning width (Figure 1E), and the distribution of preferred frequencies ^45^. While neurons in the V1 of naïve preterm mice had elevated baseline firing rates (during the presentation of the blank screen; median±MAD: Term=2.41±1.01 spikes/s; Preterm=7.49±3.65 spikes/s, p=0.061, Mann-Whitney t-test U=82, 414 neurons from N=11 term mice and 306 neurons from N=10 preterm mice), we found no significant differences in any of the orientation selectivity measures between term and preterm mice (OSI median±MAD: Term=0.78±0.05, Preterm=0.81±0.08, Mann Whitney t-test p=0.86, U=52; % selective neurons mean±SEM: Term=77.59±3.81, Preterm=72.04±5.22, t-test p=0.4, t=0.86, df=16.84; tuning width (°) median±MAD: Term=14.02±1.08, Preterm=15.19±1.73, Mann Whitney t-test p=0.29, U=70.5; distribution of orientation preferences p=0.92, X^2^ test). Increase in neuronal firing at the preferred orientation (calculated as the ratio between firing at the preferred orientation and baseline firing) was not different between term and preterm mice (median±MAD Term=6.4±1.75, Preterm=6.59±2.3, Mann-Whitney t-test p=1, U=55). Our results hence demonstrated that preterm birth in mice does not negatively impact baseline orientation selectivity and that they can be trained in an orientation discrimination task.

To prepare the mice for task training, adult term and preterm mice were implanted with headposts, and then gradually water restricted and habituated to the experimental setup and handlers over 7-10 days ^36^, after which the training began. To account for the effects of prenatal mifepristone exposure on behavior, we included an additional group of mice that were born term to dams injected with mifepristone on GD 18 (MFP Term, Figure 2A). Mice were trained on the task in 2 daily sessions of 100 trials, where each session consisted of 50 randomized presentations of 120° and 60° oriented gratings each. 120° was paired with the delivery of 10 µl of water, while 60° had no consequence (Figure 2B and C). Licks of the spout were recorded and quantified as Hits (licks to 120°), Misses (no licks to 120°), False Alarms (licks to 60°) and Correct Rejections (no licks to 60°), which were then used to quantify the discriminability [d’=z(Hits)-z(False Alarms)]. Mice were considered trained after reaching a d’ of 2 or more for 3 sessions in a row, and they were trained for a maximum of 4000 trials (20 days of training) if they did not cross the threshold within that timeframe. In our task, Hits remained high throughout the training, while False Alarms gradually decreased (Figure 2D and Supplementary Figure 1), driving an increase in d’. In term mice from both groups, d’ increased in a linear fashion and most mice crossed the threshold within 2 weeks of training (Figure 2E and 2F).

While preterm mice also displayed an increase in d’, the learning trajectory was significantly different from that of term born mice (LMM, effect of Animal Group p=0.009; Term vs Preterm p=0.007, MFP Term vs Preterm p=0.02, Term vs MFP Term p=0.78; estimated marginal means (EMM)±SE Term=2.06±0.16, MFP Term=2.01±0.18, Preterm=1.6±0.13; N=22 Term, 29 Preterm and 17 MFP Term mice; Supplementary Table 1), and a fraction of them (17.24%) failed to reach the criterion within the 4000 trial limit (Figure 2E and 2F; Kolmogorov-Smirnov test of d’ values from the last training session Term vs Preterm: D=0.43, p=0.009, Term vs MFP Term: D=0.34, p=0.14, MFP Term vs Preterm: D=0.2893, p=0.23), confirming that preterm birth negatively impacts learning. Naïve preterm mice had significantly lower Hit Rates at the onset of training, which recovered as the training progressed (Figure 2G and Supplementary Figure 1; ANOVA birth x training p=0.016, F_(2, 65)_=4.37; Bonferroni post-hoc Naïve Term vs. Preterm p=0.005, Term vs MFP Term p>0.9, MFP Term vs Preterm p=0.0013; all p>0.9 among Trained groups).

However, their False Alarm rates remained high throughout the training (Supplementary Figure 1) and were significantly higher at the end of the training compared to both groups of term mice (Figure 2H; ANOVA birth x training p<0.0001, F_(2, 65)_=10.66; Bonferroni post-hoc Naïve Term vs. Preterm p>0.9, Term vs MFP Term p=0.13, MFP Term vs Preterm p=0.04; Trained Term vs Preterm p=0.036, Term vs MFP Term p=0.53, MFP Term vs Preterm p=0.0007). To test if the effect on the Hit Rates and False Alarms was driven by preterm mice that failed to reach the criterion, we repeated the analyses while excluding those animals. We found that the interaction between birth timing and training remained for both Hit Rates and False Alarms (ANOVA birth x training HR p=0.04, F_(2, 60)_=3.2; FA p=0.001, F_(2, 60)_=7.7), demonstrating that preterm birth impacts task performance regardless of whether the animal learned the task.

Our experiments included both male and female mice of sufficient numbers to segregate the data by sex and test if birth has a sex-specific effect (Supplementary Figure 2). LMM with Animal Group and Sex as fixed effects was fitted to the learning trajectories, which revealed that sex has a significant impact on the learning trajectory (p=0.021), with female mice in all groups reaching lower d’ values (Supplementary Table 1). Analyses of Hits and False Alarm rates in trained mice (Supplementary Figure 2B and C) revealed no effect of sex, while the effect of birth was still present (ANOVA Hits: birth x sex p=0.48, F_(2,62)_=0.74; sex p=0.42, birth p=0.02; False Alarms: birth x sex p=0.12, F_(2,62)_=2.2; sex p=0.38, birth p=0.03). Our results hence demonstrate that preterm birth negatively impacts the learning trajectory of both sexes, as well as that female mice from all groups reach lower d’ values during training.

To test if the poor performance of preterm mice in the task is due to a deficit in motivation, we compared the lick rates to the rewarded cue (binned in 20 trials) during the last training session (Figure 2I). Lick frequencies in response to task cues gradually dropped during the training session, which has been proposed as a measure of motivation ^46^. An LMM was fitted to the data using Animal Group and Trial Block as fixed effects, which confirmed that the frequency of licks to the rewarded cue is reduced at the end of the session (Trial Block p<0.0001; Supplementary Tables 2), and that lick frequencies do not differ significantly between animal groups (p=0.15). Lick frequencies were different in naïve mice, with preterm mice showing less licks in response to the rewarded cue (Supplementary Tables 3; Animal Group p=0.01). However, as in trained mice, all groups showed a reduction in lick frequencies towards the end of the first session (Trial Block p=0.04). Together, these results indicate that poor performance of preterm mice (at least at the end of the training) is not due to deficits in motivation.

Our task is performed in head-fixed mice (on a treadmill), which may mask locomotion deficits in preterm mice. To test this, we tested the locomotor activity of term and preterm mice in an open field using an automated approach ^38^ (Figure 2J). ANOVA (Welch’s) revealed a significant and small reduction in locomotor activity of MFP term mice when compared to term mice, while the level of locomotor activity in preterm mice was comparable to term mice (F_(2,56.98)_=3.9, p=0.03; Dunnett’s T3 multiple comparisons test Term vs. MFP Term p=0.03, Term vs. Preterm p=0.83, MFP Term vs. Preterm p=0.41). While ANOVA found a significant difference between groups of term mice in their preference for moving close to the walls of the open field arena (Figure 2K, thigmotaxis; F_(2,51.62)_=4.08, p=0.03), post hoc tests revealed only marginal differences (Dunnett’s T3 multiple comparisons test Term vs. MFP Term p=0.07, Term vs. Preterm p=0.06, MFP Term vs. Preterm p=0.87), indicating largely preserved exploratory activity in preterm mice.

Altogether, our results demonstrate that preterm mice display impaired learning in a visual discrimination task, despite visual function, locomotor activity and motivation comparable to term born mice. As learning impairments in mouse models of neurodevelopmental conditions are often associated with impaired visual processing during task performance ^47,48^, we next characterized the processing of task cues in the V1 of trained term and preterm mice while the mice were engaged in the task.

### Selectivity for task cues is weaker in putative pyramidal neurons in preterm V1

Visual discrimination training increases the representation of and responsiveness to trained cues in the V1, both thought to support task acquisition ^40,41,49–51^. As preterm mice display impairments in task acquisition (Figure 2), we asked if the processing and representation of task cues in the V1 of preterm mice are altered compared to term mice. To address this, we used *in vivo* electrophysiology to isolate neuronal responses to task cues in trained mice while they were engaged in the task (Figure 3A and B). Firing of neurons in the V1 of trained term and preterm mice displayed the typical transient (early, 100 ms after the onset of lick) and sustained (late) components, where the peak of the sustained component typically preceded the onset of licks (Figure 3C). To test the relationship between V1 spiking and licks, we computed event correlations in 1 ms time bins for all isolated units while using licks as the reference (Figure 3D). As trained mice lick to the rewarded cue in bouts (Figure 2D), with 100-200 ms on average between licks ^52^, we restricted the correlation window to 200 ms. We found that the bulk of V1 unit firing occurs prior to lick events, with a peak at 55 ms and a significantly higher number of spiking events preceding the licks (Figure 2D; Wilcoxon matched-pairs signed rank test, p=0.018, W=-1354).

To compare the representation of task cues between term and preterm V1, we calculated the percentiles of neurons (per mouse) whose activity was significantly modulated by task cues (as determined by Wilcoxon signed-rank test of pre- and postcue firing rates). We calculated the direction of the modulation as the selectivity index (SI): (R_cue_-R_baseline_)/(R_cue_+R_baseline_), with positive values indicating cue-evoked increase in firing and negative values indicating cue-evoked suppression. We further split the units into broad spiking (putative pyramidal neurons) and narrow spiking (putative fast-spiking inhibitory interneurons) based on the width of their action potential waveform (Supplementary Figure 3; 406 neurons from 12 term mice and 253 neurons from 9 preterm mice) ^53^.

Both regular-spiking (RS) and fast-spiking (FS) neurons in term and preterm mice had comparable representation of task cues in the V1 with a slight overrepresentation of the rewarded cue (RM ANOVA with birth and the direction of modulation as factors; Figure 3E: RS Reward: birth x modulation p=0.93, F_(2,38)_=0.07; RS No Reward: birth x modulation p=0.7, F=0.36; Figure 3F: 2-tailed t-test, p=0.9, t=0.0046, df=19; Figure 3G: FS Reward: birth x modulation p=0.95, F=0.09; FS No Reward: birth x modulation p=0.24, F=1.46; Figure 3H: p=0.66, t=0.45). However, when we compared the peristimulus time histograms (PSTHs) of neurons whose firing was significantly modulated by task cues (% of all RS neurons: Term_R_=54.2, Preterm_R_=59.59, Term_NR_=42.05, Preterm_NR_=44.89; % of all FS neurons: Term_R_=50.67, Preterm_R_=78.14, Term_NR_=36.7, Preterm_NR_=54.96), we found starkly elevated firing in preterm mice [Figures 3I-L; R and NR: p=0.000, permutation test of PSTH area under the curve (AUC)].

To test the magnitude and the direction of neuronal responses to task cues, we measured selectivity index (SI) as (R_cue_-R_baseline_)/(R_cue_+R_baseline_), where R_cue_ is the firing rate during cure presentation. SI ranges from -1 to 1, where the indices of suppressed neurons fall below 0, and those of excited neurons above it. As the distribution of SI values in a population of neurons can provide more information about cue evoked activity than population averages, we compared the cumulative distributions of SI values from neurons significantly modulated by cues in term and preterm mice. Despite increased firing, RS neurons in preterm V1 had reduced selectivity for task cues, evident in the distribution of selectivity indices (SI) shifted towards lower values (Figure 3M; SI median±MAD Term_R_=0.34±0.12, Preterm_R_=0.19±0.09, K-S test: D=0.35, p=0.0007; Term_NR_=0.21±0.1, Preterm_NR_=0.08±0.04, K-S test, D=0.46, p=0.00005). On the other hand, a large fraction of FS neurons in preterm mice had high SI values, with no changes in the selectivity for the non-rewarded cue (Figure 3N; Term_R_=0.36±0.13, Preterm_R_=0.43±0.23, K-S test: D=0.29, p=0.000006; Term_NR_=0.17±0.1, Preterm_NR_=0.13±0.09, D=0.11, p=0.36).

As locomotion can increase V1 firing rates ^54^, we measured the amount of movement on the treadmill during the task (Supplementary Figure 4A, N=8 term and 5 preterm mice). Locomotion during the task was very variable in both groups of mice and overall not statistically different (p=0.37, permutation test of AUC). Elevated firing in the V1 was task-specific, as PTSHs revealed similar levels of activity in both groups of mice outside of task context (cues presented in the absence of the reward delivery spout, Supplementary Figure 4B, compare with Figure 3G). Further, licking activity in preterm mice was slightly elevated (Supplementary Figure 4B; # of licks per bout Term_R_=23.65±2.23, Preterm_R_=28.48±1.49, Term_NR_=8.37±1.28, Preterm_NR_=13.31±3.056; birth x cue p=0.97, F_(1, 22)_=0.00065; birth p=0.024, F=5.85; cue p<0.0001, F=46.26), but the level of cue-evoked sustained activity in the V1 [mean sustained response (0.1-1.5 s)/mean transient response (0-0.1s) ^55^] was not significantly correlated with lick frequency in either group of mice (Supplementary Figure 4E; Pearson’s correlation, Term_RS_ p=0.97, Term_FS_ p=0.22, Preterm_RS_ p=0.28, Preterm_FS_ p=0.08).

Neurons in the V1 display activity related to behavioral outcomes ^56^. While the behavioral relevance of this activity is unclear, we checked if this activity can be detected in the V1 of preterm mice. Trained mice typically had a 100% hit rate during the last session (Supplementary Figure 1), so we compared the responses of RS and FS neurons during the Correct Rejections (CR) and False Alarms (FA) between term and preterm mice. RS neurons represented outcomes to a similar degree in term and preterm mice (Supplementary Figure 5A; ANOVA cue x modulation: Correct Rejection p=0.24, F_(2,38)_=1.47; False Alarms p=0.7, F=0.36), despite elevated neuronal activity in preterm mice and weakened SIs for non-rewarded cue during both outcomes (Supplementary Figure 5B and C; p=0.000, permutation test of PSTH AUC; SI Term_CR_=0.2±0.09, Preterm_CR_=0.09±0.06; K-S test, D=0.48, p=0.0002; Term_FA_=0.29±0.09, Preterm_FA_=0.09±0.25; D=0.37, p=0.02). While FS neurons of term and preterm mice represented the behavioral outcomes to a comparable degree (Supplementary Figure 5D; ANOVA cue x modulation: Correct Rejection p=0.36, F=1.03; False Alarms p=0.75, F=0.28), neuronal activity was elevated in preterm mice and the cue-evoked peak in activity was barely detectable during the False Alarms (Supplementary Figures 5E and F; p=0.000, permutation test of PSTH AUC). Interestingly, selectivity for the non-rewarded cue in FS neurons displayed a significant shift towards lower values once it was segregated into behavioral outcomes; SI Term_CR_=0.26±0.12, Preterm_CR_=0.17±0.11; K-S test, D=0.23, p=0.009; Term_FA_=0.36±0.18, Preterm_FA_=0.23±0.32; D=0.31, p=0.001).

Altogether, our data demonstrates that preterm mice have intact representation of task cues and behavioral outcomes in the V1, despite significantly elevated neuronal activity and overall weakened selectivity for the task cues. Increase in neuronal activity in preterm mice appeared to be specific to task context, suggesting the involvement of top-down modulation. Therefore, we next examined the neuronal responses to task cues in term and preterm prefrontal cortex (PFC), a top-down area that sends major glutamatergic monosynaptic input to the V1 and modulates its activity during discrimination tasks ^57,58^.

### Representation of task cues is altered in the PFC of preterm mice

Within the PFC, anterior cingulate cortex (ACC) sends and receives dense glutamatergic projections to and from the V1 ^57,59,60^. Neurons in the ACC are not responsive to sensory cues in naïve mice, but develop robust context-specific responses to cues during training in sensory discrimination tasks ^61–63^. Further, optogenetic activation of ACC neurons that project to the V1 (ACC→V1) evokes a robust increase in V1 firing rates and promotes behavioral performance in visual tasks ^57,58^, while inactivating ACC impairs visual discrimination (Supplementary Figure 6) ^64^, suggesting that ACC exerts top-down modulation of cue processing in the V1 of trained mice. We therefore hypothesized that elevated activity and altered cue selectivity in preterm V1 during task performance reflects the activity and selectivity in their PFC. To test this, we recorded the activity of the PFC while the mice were engaged in the task (Figure 4A and B) and sorted the isolated neurons by their waveform width as in the V1. While most of the units recorded were from deep cortical layers (5 and 6), our recordings collected units from layer 2/3 as well (Figure 4B). As expected, putative FS interneurons had higher precue firing rates overall in both groups of mice (median±MAD FS: Term=8.61±6.97, Preterm=7.57±5.75 spikes/s 143 and 183 units respectively from 10 and 9 preterm mice; RS: Term=2.7±1.08, Preterm=2.46±1.2 spikes/s, 70 and 170 units respectively), but pre-cue firing rates between term and preterm mice were comparable (Mann-Whitney t-test, FS: p=0.6, U=43; RS: p=0.96, U=25). Similar to V1, the firing of PFC neurons preceded the licks, with event correlogram revealing peaks at 14 and 65 ms prior to lick events, as well as a significantly higher number of spiking events prior to lick events (Figure 4C; Wilcoxon matched-pairs signed rank test, p<0.0001, W=-2770).

We next evaluated cue representation and found that the non-rewarded cue had a significantly weaker representation in RS neurons of preterm mice, shifting the overall representation in favor of the rewarded cue (Figure 4D; ANOVA Reward: birth x modulation, p=0.6, F_(2,34)_=0.52; No Reward: p=0.014, F=4.88; Figure 4E: Mann-Whitney test p=0.045, U=20.5). Significantly modulated neurons in both groups of mice had an overall higher firing rate than the entire recorded population (median±MAD; precue firing Term=4.3±2.84, Preterm=3.27±1.73 spikes/s; % of all neurons Term_R_=78.7, Term_NR_=52.9, Preterm_R_=88.1, Preterm_NR_=23.01), and their PSTHs revealed a significantly increased pre- and postcue firing rate in preterm mice (Figure 4F and G; p=0.000, permutation test of PSTH AUC). Additionally, the PSTH to the non-rewarded cue in preterm mice had a very irregular shape with no clear peak, suggesting weakened selectivity of prefrontal RS neurons for the non-rewarded cue in preterm mice (Figure 4G). Indeed, while the SI values for the rewarded cue were not different between term and preterm mice (Figure 4H: median±MAD, Term_R_=0.72±0.13, Preterm_R_=0.71±0.1; K-S test, D=0.13, p=0.36), the SI values for the non-rewarded cue were significantly lower in preterm mice (Figure 4H: Term_NR_=0.33±0.09, Preterm_NR_=0.16±0.07; K-S test, D=0.4, p=0.000014).

Previous work demonstrated a strong correlation between prefrontal responsivity to task cues during training and the progressive increase in performance ^61^. As the SI reflects the responsiveness of neurons to task cues, we tested the correlation between the SI values (averaged per animal) and d’ values during the recording session, as well as the strength of the SI values in predicting behavioral performance using linear regression when a significant correlation was detected. In term mice, SI values for the rewarded and non-rewarded cues were not significantly correlated with d’ values (Pearson’s correlation, Figure 4I). In contrast, SI values for the rewarded cue were significantly negatively correlated with d’ in preterm mice (r=-0.75, p=0.02) and were a predictor of performance (Figure 4J and Supplementary Tables 4). The correlation between SI values for the non-rewarded cue and d’ in preterm mice was not significant (Figure 4J).

Our bulk recordings did not isolate the activity of PFC neurons that project to the V1, making it difficult to draw connections between the activity patterns in the V1 and PFC. To do that, we used an intersectional viral approach to optogenetically label the V1-projecting PFC neurons, where we injected the retrograde AAV2 variant encoding the Cre-recombinase into the V1 ^65^ and DIO-ChR2-mCherry into the PFC (Supplementary Figure 7A and B). We used 10 ms pulses at 473 nm after the behavioral sessions ended to elicit the ChR2-mediated responses and isolate the activity of PFC→V1 neurons (short latency responses with >50% reliability, Supplementary Figure 7C) ^66^. Interestingly, we found a significantly increased representation of the rewarded cue in PFC→V1 neurons of preterm mice (Supplementary Figure 7D; 28 neurons from N=9 term mice, 35 neurons from N=6 preterm mice; ANOVA; Reward: birth x modulation, p=0.0017, F_(2,26)_=8.27, No Reward: birth x modulation, p=0.38, F=0.99). PSTHs of neurons significantly modulated by the cue (% of all neurons: Term_R_=66.67, Preterm_R_=98.33, Term_NR_=55.21, Preterm_NR_=58.45) revealed elevated activity of PFC→V1 neurons in preterm mice after the presentation of the rewarded cue, especially in the late stages of their response (Supplementary Figure 7F; p=0.000, permutation test of AUCs). SI values for the rewarded cue were also substantially higher in preterm than in term mice (median±MAD, Term_R_=0.31±0.31, Preterm_R_=0.57±0.09; K-S test, D=0.79, p=3.15x10^-6^). PSTHs of PFC→V1 neuron responses to the non-rewarded cue revealed irregular firing in preterm mice, in line with weakened activity evoked by the non-rewarded cue in V1 and PFC (Supplementary Figure 7G; p=0.000, permutation test of AUCs). The SI values for the non-rewarded cue were also elevated in preterm mice (Term_NR_=0.11±0.19, Preterm_NR_=0.2±0.07), but the differences in their distribution between term and preterm mice were marginally significant (K-S test, D=0.41, p=0.06), likely due to low number of significantly modulated neurons (Term=16 and Preterm=23).

The representation of cues in FS neurons was similar to RS neurons, although the difference in the representation of the non-rewarded cue was not statistically significant (Figure 4K; ANOVA Reward: birth x modulation, p=0.71, F_(2,34)_=0.34; No Reward: p=0.2, F=1.67). As in RS neurons, the representation of cues was slightly shifted in favor of the rewarded cue (Figure 4L), but it was not statistically different between term and preterm mice (Mann-Whitney test p=0.17, U=28). However, when we compared the PSTHs of neurons significantly modulated by task cues between term and preterm mice (% of all neurons Term_R_=86.42, Term_NR_=56.37, Preterm_R_=90.8, Preterm_NR_=37.6), we found that the responses to both cues in preterm mice were significantly blunted (Figure 4M and N; p=0.000, permutation test of PSTH AUC). Similar to V1 FS neurons, the median SI of prefrontal FS neurons for the rewarded cue in preterm mice was marginally larger than in term mice, while the SI for the non-rewarded cue was lower (Figure 4O; median±MAD SI: Term_R_=0.52±0.17, Preterm_R_=0.58±0.11, K-S test: D=0.19, p=0.02; Term_NR_=0.21±0.06, Preterm_NR_=0.092±0.07, K-S test, D=0.36, p=0.00097). None of the SI values in either group of mice correlated with d’ (Figures 4P and Q; Pearson’s correlation).

FS interneurons can remain undetected during spike sorting of bulk electrophysiological recordings due to their low spontaneous firing rates ^67^. To ensure that our spike sorting approach reliably captured the weaker responsivity to task cues in preterm mice, we used a transcriptional enhancer S5E2 ^68^ to transduce FS interneurons with ChR2 for optogenetic identification (“optotagging”, Supplementary Figure 8). S5E2-mCherry only partially colocalized with Parvalbumin (PV) signal in the ACC, which was significantly weaker compared to other cortical areas (Supplementary Figure 8B), as previously reported ^69^. As expected, 10 ms pulses of blue light evoked strong rhythmic firing in transduced neurons (Supplementary Figure 8C) ^66^, with all isolated neurons having a narrow waveform (average spike half-width: 0.33±3.2x10^-6^ms, N=50 units from 5 term mice and 52 units from 4 preterm mice). Indeed, the pre-cue firing rate of optotagged FS units was significantly lower than that of FS units isolated through spike sorting of bulk electrophysiological data (Figure 4B and 4C; median±MAD, N=units; Term/Preterm narrow-spiking units=5.9±4.79 spikes/s vs Term/Preterm optotagged S5E2^+^ units=1.45±1.45 spikes/s; Mann-Whitney t-test, p<0.0001, U=11297), with no significant differences between term and preterm optotagged FS units (Term=0.63±0.63 spikes/s, Preterm=1.69±1.52; Mann-Whitney t-test, p=0.18, U=1104). While the fractions of neurons significantly modulated by task cues was overall lower in the optotagged population (Supplementary Figure 8D; compare with 4K), they were not significantly different between term and preterm mice (ANOVA Reward: birth x modulation: p=0.91, F_(2 14)_=0.094; No Reward: p=0.99, F=0.014). PSTHs of significantly modulated optotagged neurons (% of all neurons: Term_R_=61.56, Preterm_R_=71.18, Term_NR_=52.66, Preterm_NR_=53.26) revealed their weak responsivity to task cues in preterm mice (p=0.000, permutation test of PSTH AUC), similar to what we found in FS neurons isolated through bulk recordings (Supplementary Figure 8E and F; compare with Figure 4M and N). However, the selectivity for the rewarded cue was significantly weakened in preterm mice, while the selectivity for the non-rewarded cue appeared comparable between term and preterm mice (median±MAD SI: Term_R_=0.78±0.22, Preterm_R_=0.38±0.15, K-S test: D=0.79, p=0.000003; Term_NR_=0.31±0.06, Preterm_NR_=0.35±0.13, K-S test: D=0.41, p=0.057).

Our results hence demonstrate a significant shift in the representation of task cues towards the rewarded one in putative principal neurons of preterm PFC, along with pronounced disinhibition of their responses to task cues. This shift comes at the expense of the non-rewarded cue, which elicits blunted responses in both putative pyramidal neurons and interneurons.

Surprisingly, the selectivity of the population responses to the rewarded cue was predictive of performance in preterm, but not in term mice. As the preterm mice that underwent electrophysiological recordings had comparable d’ values to term mice (Figure 4I and J; mean±SEM Term=2.56±0.16, Preterm=2.83±0.13, N=9 mice/group), our results indicate divergent processing of task cues in preterm mice and further suggest that preterm mice use different strategies to encode behavioral outcomes. To test this further, we next compared the representation of behavioral outcomes to the non-rewarded cue in prefrontal RS and FS neurons of term and preterm mice and modeled their ability to classify trial-by-trial behavioral outcomes.

### Cue-evoked activity of prefrontal neurons is a weaker predictor of behavioral outcomes in preterm mice

Prefrontal cortex is critical for many forms of goal-directed behavior and its activity is predictive of behavioral outcomes across different tasks ^70–72^. Surprisingly, the population selectivity for the rewarded cue was predictive of performance in preterm but not in term mice (Figure 4), suggesting that preterm mice use a strategy different from term mice to encode task-relevant information. Indeed, both the increase in licks to the rewarded cue and suppression of licks to the non-rewarded cue drive the increase in performance in preterm mice, while the suppression of behavioral responses to the non-rewarded cue drives the performance increase in term mice (Figures 2 and Supplementary Figure 1). We therefore hypothesized that the PFC of term and preterm mice uses divergent strategies to encode behavioral responses to task cues, and to the non-rewarded cue in particular as preterm mice show increased False Alarm rates (Figure 2H). As almost 100% of behavioral responses to the rewarded cue in trained mice from both groups are Hits (Figure 2 and Supplementary Figure 1), we focused our analysis on neuronal responses to the non-rewarded cue, i.e. Correct Rejections (CRs) and False Alarms (FAs; Figure 2C).

We first tested the representation of behavioral outcomes in prefrontal neurons and surprisingly found no differences between term and preterm mice in either regular spiking (Figure 5A; ANOVA: birth x modulation CR: p=0.81, F_(2,34)=_0.21, FA: p=0.6, F=0.51) or fast-spiking neurons (Figure 5B; CR: p=0.65, F=0.43, FA: p=0.45, F=0.82). PSTHs of significantly modulated RS neurons (% of all neurons Term_CR_=33.5, Term_FA_=41.42, Preterm_CR_=35, Preterm_FA_=49.03) revealed elevated firing and irregular cue-evoked activity in preterm mice during both outcomes (Figure 5C; p=0.000, permutation test of AUCs). While cue-evoked activity in preterm FS neurons also appeared irregular, analysis revealed that term and preterm FS responses to the non-rewarded cue are comparable during CR but not during FA trials (Figure 5D, CR p=0.125, FA p=0.022, permutation test of AUCs; % of all neurons Term_CR_=48.53, Term_FA_=53.34, Preterm_CR_=46.34, Preterm_FA_=56.94). Selectivity indices were shifted towards lower values in both cell populations of preterm mice during both behavioral outcomes (Figure 5E; median±MAD, K-S test for all; RS CR Term=0.49±0.21, Preterm=0.13±0.09, D = 0.44, p=0.000042; RS FA Term=0.4±0.16, Preterm=0.29±0.18, D = 0.26, p=0.014; FS CR Term=0.14±0.07, Preterm=0.09±0.06, D=0.24, p=0.044; FA Term=0.37±0.11, Preterm=0.08±0.19, D=0.57, p=1.2x10^-10^). Further, FS neurons in term mice showed higher SI values during FA trials, while the same was true for RS neurons in preterm mice (Mann-Whitney test, Term FA vs CR: RS p=0.82, U=150, FS p<0.0001, U=526.5; Preterm FA vs CR: RS p<0.0001, U=1593, FS p=0.7, U=2517).

To test if neuronal responses can predict behavioral outcomes on a single trial basis, we used logistic regression with CR and FA as binary outcomes and cue-evoked (baseline subtracted) spike rates (Figure 5F and G) as classifiers. We included all isolated neurons in the analysis as the number of neurons significantly modulated by the non-rewarded cue was low for some mice. Cue evoked spike rate distributions of all isolated neurons mirrored the trends seen in SI values of neurons significantly modulated by task cues, with FS and RS neurons in term and preterm mice, respectively, preferentially responding to the non-rewarded cue during the FA trials and weaker cue-evoked firing in preterm FS neurons during both trial outcomes (Figure 5K; median±MAD spikes/s, RS: Term_CR_=1.5±1.43, Preterm_CR_=0.65±1, K-S test p=0.011, D=0.23; Term_FA_=2±1.15, Preterm_FA_=2.3±3.31, p=0.0008, D=0.28; FS: Term_CR_=1.28±1.27, Preterm_CR_=0.34±0.89, p=0.0001, D=0.26; Term_FA_=4.41±4.29, Preterm_FA_=1.25±1.86, p=0.93x10^-6^, D=0.31). We constructed Receiver Operating Characteristic (ROC) curves for each group of mice and compared their performance first to shuffled outcomes from each group, and then to each other (Figure 5H) ^73^. We found that neuronal responses to the non-rewarded cue in both groups of mice classified trial outcomes significantly better than shuffled data (au(ROC)±SE: Term=0.703±0.0058, 7928 trials; Preterm=0.65±0.005, 11164 trials; Shuffled term=0.5±0.006, Shuffled preterm=0.5±0.005; Term vs Shuffled term, z=23.26, p<0.0001, Preterm vs shuffled preterm, z=19.93, p<0.0001). However, firing rates of term mice performed significantly better as classifiers than those of preterm mice (z=6.79, p<0.0001). ROC values for each mouse did not correlate with the fraction of significantly modulated neurons nor with their SI values (Pearson’s r=-0.24, p=0.42 and r=0.25, p=0.4, respectively), indicating that including all recorded neurons in the model did not impair the classification.

Our results hence provide further evidence for weak selectivity of prefrontal neurons for the non-rewarded cue in preterm mice, indicating divergent processing of task cues and encoding of behavioral outcomes in preterm PFC. Preterm birth is a significant risk for delays in cognitive development, especially in the development of response inhibition ^74^, suggesting that the impairments in the processing and representation of task cues in preterm PFC reflect a maturational deficit. As prior research found weak representation of non-rewarded cue in the brains of adolescent mice trained on an auditory task ^75^, we next tested if the representation and processing of cues in our task is similar between adult preterm mice and adolescent term mice.

### Similar encoding of behavioral outcomes to the non-rewarded cue in adolescent term and adult preterm mice

To test for impaired maturation of preterm PFC, we examined the representation and processing of task cues in adolescent, term born male and female mice and compared it to adult preterm mice (Supplementary Figure 9). To prepare the adolescent mice for the task, they were implanted with a headpost between postnatal days 21 and 24 and begun habituation and water restriction 4 days after the implantation (Figure 6A). The training started between postnatal days 32 and 35 and lasted until mice crossed the criterion (all mice learned the task, N=18).

Adolescent mice performed comparable to term mice (6B, LMM with Age as Fixed Factor and Animal and Training Day as Random Factors, p=0.42, Supplementary Tables 5), and showed a significant increase reduction in FA rate between naïve and trained stages (Figure 6C; RM ANOVA: response x training p<0.0001, F_(1,17)_=40.5; HR p=0.07, FA p<0.0001), indicating that a decrease in response inhibition drives behavioral performance in our task regardless of age tested.

We next recorded the activity of prefrontal neurons in adolescent mice during task performance (N=324 neurons from 9 mice), which we classified into regular and fast-spiking neurons as in Figures 3 and 4 (Figure 6D and E). As previously reported ^75^, the representation of the non-rewarded cue was significantly weaker than that of the rewarded cue (RM ANOVA cue x modulation Figure 6E: p<0.0001, F_(2.2,17.79)_=37.11; Figure 6F: p<0.0001, F_(2.5,20.04)_=18.37).

Similar to preterm mice, the representation of the non-rewarded cue was significantly weaker only in RS neurons when compared to adult term mice (Supplementary Figure 9). While the rewarded cue elicited substantially higher activity in both RS and FS neurons (Figure 6G; p=0.000, permutation test of AUCs), the PSTHs of neurons whose activity was significantly modulated by task cues revealed that cue-evoked activity was lower in adolescents than in adult term mice (% of all neurons: RS R=72.98, NR=29.42, FS R=93.11, NR=35; Supplementary Figure 9). Preference for the rewarded cue was also evident in SI values, with higher selectivity for the rewarded cue in both RS and FS of adolescent mice (Figure 6H; median±MAD RS R=0.4±0.16, NR=0.25±0.16; K-S test, D=0.31, p=0.005; FS R=0.41±0.12, NR=0.26±0.18; D=0.27, p=0.01).

In terms of behavioral outcomes to the non-rewarded cue, both CR and FA were equally represented in the adolescent PFC (RM ANOVA outcome x modulation Figure 6E: p=0.17, F_(2.16)_=1.96; Figure 6F: p=0.95, F=0.04). PSTHs of neurons significantly modulated by the non-rewarded cue revealed slightly higher neuronal responses during FA trials in both neuronal populations (Figure 6I; p=0.000, permutation test of AUCs; % of all neurons: RS CR=15.71, FA=34.03, FS CR=41.73 FA=44.75), but their SI values were comparable (Figure 6J; median±MAD RS CR=0.28±0.16, FA=0.3±0.22; K-S test, D=0.21, p=0.47; FS CR=0.25±0.16, FA=0.29±0.21; D=0.12, p=0.89). We next isolated single trial firing for adolescent neurons and tested if their evoked firing can accurately classify the two behavioral outcomes to the non-rewarded cue (Figure 6K; term and preterm data from Figure 5M added for comparison). While the logistic model built on adolescent evoked firing rates performed significantly better than the shuffled data (p<0.0001, z= 22.92), there was no significant difference between the auROC of preterm mice and that of adolescent mice (adolescent vs preterm p=0.17, z=1.38; adolescent vs term adult p<0.0001, z=-5.76).

### Life-long environmental enrichment fails to improve the learning trajectory of preterm mice

Adult preterm mice and adolescent term mice display similarly weak representation of the non-rewarded cue in the PFC (Supplementary Figure 10), as well as similar encoding of its behavioral outcomes (Figure 6K), suggesting a maturational delay in the PFC of preterm mice. We therefore tested if environmental enrichment (ENR), a classical paradigm that promotes sensory maturation and rescues cognitive deficits in animal models of prematurity-related brain injury, improves learning in preterm mice ^21,28,31,76,77^. To do this, we housed timed-pregnant dams with their term and preterm litters from postnatal day 5 (P5) to weaning (P28) in standard rat cages supplemented with a running wheel, multiple play and chew objects (such as gnawing sticks and tunnels), as well as a non-related dam with an age-matched litter for social enrichment. The object location was shuffled in the cage every 2-3 days. Prior to P5, pups were housed in a standard cage with enrichment for the dams in the form of igloos, extra nesting material and sunflower seeds. After weaning, term and preterm mice were housed in groups of 2-3 in standard cages supplemented with an igloo, small running platform and gnawing sticks.

When mice reached adulthood (>2 months of age), they were prepared for the visual discrimination task, water-restricted and trained to criterion or to 4000 trials if they did not pass the criterion (Figure 7A). While term mice had a steep learning curve, the enrichment had a negative impact of their performance, lowering the overall d’ values when compared to term mice that received no enrichment [LMM with Enrichment (ENR) and Birth as Fixed Effects, and Animal and Training Day as Random Effects; ENR p<0.0001, Estimated Marginal Means (EMM) Term ENR=1.6±0.2, Term=2.22±0.19; Supplementary Tables 6]. However, just like term mice that received no enrichment, all ENR mice learned the task within 15 days (Figure 7B). In contrast, preterm mice that received ENR had a substantially impaired learning curve (LMM: Birth p<0.0001) and performed worse than preterm mice that received no enrichment (EMM Preterm ENR=1.003±0.68, Preterm=1.6±0.14). While a significant fraction of ENR preterm mice failed to learn the task within 4000 trials (27.27%), the distribution of final d’ values was not significantly different between ENR term and preterm mice (Figure 7C; K-S test, D=0.46, p=0.16).

When we analyzed Hit Rates and False Alarms in ENR mice and compared them to standardly reared term and preterm mice (STD; Figure 7D and E, compare with Figure 2G and H), we found a significant interaction between training, enrichment and birth timing (RM ANOVA, p=0.002, F=28.49, Supplementary Tables 7). Post-hoc tests revealed no significant differences between naïve ENR term and preterm mice in Hit Rates and a significant reduction in trained preterm mice (naïve ENR Term vs Preterm p=0.19, trained p=0.006; Figure 7D). Both groups of mice had significantly reduced HR compared to STD mice at the onset of training, and this trend persisted in preterm mice only (Supplementary Tables 7). There were no significant differences in FA rates between term and preterm mice raised in ENR (Figure 7E; naïve ENR Term vs Preterm p=0.99, trained p=0.99; Supplementary Tables 7), but trained ENR preterm mice had comparable FA rates to STD trained term mice (p=0.99).

## Discussion

Through extensive electrophysiological characterization of primary visual (V1) and prefrontal cortices in adult preterm mice trained on a visual associative task, our study identified impaired representation and processing of sensory cues in the prefrontal cortex (PFC) as a potential driver of cognitive impairments common in preterm born children and adolescents.

Diminished representation of the non-rewarded cue was present in putative pyramidal neurons of the preterm PFC, while both putative pyramidal neurons and interneurons displayed reduced selectivity for the non-rewarded cue. Activity evoked by the non-rewarded cue more accurately predicted behavioral outcomes in term mice, pointing to their divergent encoding in term and preterm PFC. Indeed, our study revealed substantial similarities in the representation of the non-rewarded cue and encoding of its behavioral outcomes between adult preterm mice and term adolescent mice, indicating that preterm birth disrupts the maturational trajectory of the PFC. Environmental enrichment, however, did not improve the learning deficit of preterm mice. Unexpectedly, enrichment impaired learning in both term and preterm mice, highlighting previous findings on the variability of its outcomes in laboratory animals ^78,79^.

Our study identified substantial hyperactivity of cue-modulated neurons in the PFC and V1 of preterm mice. In the PFC, the hyperactivity was specific to regular-spiking (RS), putative pyramidal neurons, while the response of prefrontal fast-spiking (FS) interneurons to both cues was blunted in preterm mice. Our findings hence suggest that elevated activity of prefrontal pyramidal neurons is a consequence of dysfunctional perisomatic inhibition, but do not exclude a dysfunction of other interneuronal subtypes: task-related neurons in preterm PFC show elevated firing even before the onset of cue presentation, when FS interneurons do not show reduced firing rates. Indeed, abnormalities in multiple interneuron subtypes have been reported in postmortem studies of prematurely born fetuses and in animal models of prematurity ^17,19,20,80^. Future electrophysiological studies can now test the function of other interneuronal subtypes in the preterm PFC.

While cue-evoked activity of FS interneurons in preterm mice may contribute to the elevated firing of prefrontal RS neurons, it is unclear which inputs fail to sufficiently activate the FS interneurons upon cue presentation. These inputs are unlikely to come from local pyramidal neurons-their responses to the rewarded cue in preterm mice are robust. Another major source of excitation to prefrontal FS interneurons is the mediodorsal thalamus (MDT), whose role in learning is well established ^81–83^. Optogenetic activation of MDT results in a reduction of putative pyramidal neuron firing rates due to strong feedforward inhibition mediated by MDT inputs onto FS interneurons ^84,85^. Our results point to these inputs being dysfunctional in preterm mice, resulting in elevated activity of prefrontal pyramidal neurons during cue presentation. This notion is in line with impairments in thalamic structure observed in preterm infants and adolescents ^13,14^ and the hypothesized role of thalamic dysfunction as a driver of impaired cortical function after preterm birth ^86^. In sensory areas of mice, preterm birth accelerates the refinement of thalamocortical inputs through serotonin signaling ^87^, but it is unclear if prematurity has the same effect on higher order thalamic nuclei, such as MDT, whose cortical target is the PFC. We plan to address this in our future studies.

We previously found that juvenile preterm mice show sparser neuronal firing rates and increased size of putative inhibitory synapses in the V1 compared to term-born mice ^39^. Our current results indicate that these changes in the activity of V1 in preterm mice are transient, given the comparable neuronal firing rates of V1 neurons outside of task context in term and preterm mice. Furthermore, the baseline orientation selectivity is intact in preterm mice and the firing of V1 neurons is elevated only during task performance, indicating that visual processing deficits common after preterm birth are due to top-down modulation and not local V1 computations ^88,89^. A previous study demonstrated that optogenetic activation of V1-projecting prefrontal neurons results in robust elevation of V1 firing ^58^. Indeed, our results found increased selectivity for the rewarded cue in V1-projecting PFC neurons, as well as their heightened activity, supporting that the increase in V1 neuron firing during task performance originates from top-down projections. Our finding that both RS and FS neurons show similarly elevated task-related activity further supports the involvement of top-down inputs in altered processing of task cues in the V1 of preterm mice and warrant further investigation, especially given the previous reports of top-down sensory processing impairments in preterm infants ^90–92^. The firing rates of V1 neurons in our study are somewhat higher than those previously reported in literature ^53^ and it is possible our analysis failed to consistently separate single units from multiunits. On the other hand, extracellular electrophysiology is biased towards neurons that fire action potentials at higher frequencies (such as layer 5 pyramidal neurons) as they are easier to isolate from the background ^93,94^. Indeed, this is demonstrated in our study by comparison of optogenetically identified FS neurons in the PFC and those identified through spike sorting of bulk signals where we found that optotagged FS neurons show lower median firing rates before cue onset. As our unit isolation criteria was kept constant for all groups of mice in this study, the presence of multiunit data is unlikely to bias the results towards elevated firing in preterm mice, especially as their prefrontal FS neurons display weak cue-evoked firing.

While we have not addressed the precise circuit mechanisms that drive behavioral impairments and altered processing of task cues in preterm mice in this study, we attempted to test for the presence of developmental delay in the preterm PFC as they are common in the preterm population ^95,96^. Interestingly, adolescent term-born mice show similarly reduced representation of the non-rewarded cue in putative prefrontal pyramidal neurons, and their neurons classify its behavioral outcomes comparable to preterm mice. As the preterm mice used in our study were adults that ranged in age from 2 to 7 months, our findings indicate that preterm-birth associated impairments in prefrontal function persist through adulthood, arguing for a disrupted (instead of arrested) development. This argument is strengthened by our findings that environmental enrichment, a commonly used developmental intervention that rescues arrested sensory maturation ^77^, is ineffective in mitigating the deficits seen in preterm mice. In fact, enrichment impaired performance in both term and preterm mice, highlighting the variability in its effects on behavior, particularly in mouse models of neurological conditions ^79,97^. Previous work demonstrated that enrichment negatively impacts motivation in certain tasks ^98^. Indeed, ENR mice in our study had significantly reduced Hit Rates and False Alarms, suggesting that their motor output was reduced, impairing their performance. Future studies can test if the type of enrichment used in our study is optimal for promoting the maturation of prefrontal areas, especially given recent studies that demonstrated the positive effects of tactile enrichment on sensory information coding in the somatosensory and motor areas of mice ^30^.

An important finding of our study is that adolescent PFC encodes task-relevant information differently to adult PFC. Our results confirmed a previous report that the non-rewarded cue has weak representation in adolescent PFC, tilting the overall cue representation towards the rewarded cue ^75^. Our study, however, identified significantly weaker cue-evoked activity in adolescent PFC when compared to adults, especially in FS interneurons, which showed remarkable similarities in firing to adult preterm mice. Our results hence support that adolescence is a sensitive period for the development of prefrontal function in the context of goal-directed behaviors ^12^. Adolescents in our study ranged 41-48 days of age at the time of the recording, while adult term mice were 80-90 days old, indicating that task-related activity in the PFC rapidly and substantially changes between late adolescence and adulthood. Similar to the previous report ^75^, we also found that adolescents performed in our task as well as adult mice, and that they learned to inhibit the licking after the presentation of the non-rewarded cue as successfully as adult mice. Altogether, these findings imply that behavioral inhibition in the adolescent brain is either mediated by other brain areas or by the PFC in a manner distinct from that in adult brain. It will be important to address this in future studies, especially in the context of prematurity, given the similarities in the encoding of the non-rewarded cue in the PFC of adult preterm and adolescent term mice.

While our study identified potential circuit loci of impaired neurodevelopmental outcomes after preterm birth, it did not test if optogenetic or chemogenetic manipulations of circuit- or cell type-specific activity improved behavioral performance of preterm mice in our task. Chemogenetic activation of prefrontal FS interneuron promotes behavioral response inhibition ^99^, but it is unlikely that chemogenetic or pharmacological approaches would have the same effect in adult preterm mice given the cue- and outcome-specific deficits in their activity. Such specificity could be achieved via optogenetic manipulation, but prolonged optogenetic stimulation (such as during task training) can result in neurotoxicity and tissue damage ^100^.

Previous research highlighted windows of increased plasticity and vulnerability during prefrontal development beginning shortly after birth ^101^. It is possible that targeted activity and/or circuit manipulations during these windows could improve neurocognitive outcomes after preterm birth, but the mechanistic impact of prematurity on the developmental trajectory of the PFC is unknown. Mice are considered premature at term birth already, and the first 7-10 postnatal days is typically thought to be the equivalent of third trimester in fetal development ^102^. While we cannot reliably estimate what would be the equivalent of human fetal age of prematurely born mice, our study nevertheless highlights the long-term impact of advanced birth on prefrontal function. As most models of prematurity use term born mice subjected to neonate hypoxia/ischemia ^103^, future studies need to address the differences and similarities between different models of prematurity, especially in the context of prefrontal function in prematurely born humans.

In conclusion, our results identify impairments in the representation and processing of sensory cues in the visual and prefrontal cortices of preterm mice, and provide further evidence of interneuron dysfunction in the preterm brain ^17,18,103^. Our study highlights the profound and persistent impact of preterm birth on the function of top-down circuits, further stressing the vulnerability of the PFC to perinatal and postnatal stressors. ^12^

## Acknowledgements

The authors would like to thank the Cang and Liu labs at the Departments of Biology and Psychology, as well as the Program in Fundamental Neuroscience, for generous access to their confocal microscopes. The authors would like to acknowledge Dr. Francesca Sciaccotta and Alia Minaya for their help with data analysis.

## Inclusion & ethics

All authors of this study have fulfilled the criteria for authorship required by Nature Portfolio journals, as their participation was essential for the design and implementation of the study.

## Data Availability

All data from this study is available upon reasonable request. Source data is provided with this paper.

## Code Availability

All analysis codes used in this study are available online (CED website).

## Author Contributions

E.M. and A.R. performed all electrophysiological experiments and confocal imaging, and E.M., V.P. and A.R. performed all behavioral experiments, with assistance from M.F., L.Y.D., O.B., S.F., J.S., L.E., T.D.H. and P. A.-A. All authors contributed to the analysis of the behavioral data. Electrophysiological data was analyzed by X.T. and A.R. A.R. conceived the study and wrote the manuscript with input from all authors.

## References

1. Heuvelman, H. et al. Gestational age at birth and risk of intellectual disability without a common genetic cause. Eur. J. Epidemiol. 33, 667–678 (2018).

2. Perapoch, J. et al. Prematurity and ADHD in Childhood: An Observational Register-Based Study in Catalonia. J. Atten. Disord. 25, 933–941 (2021).

3. Crump, C., Sundquist, J. & Sundquist, K. Preterm or early term birth and risk of attention-deficit/hyperactivity disorder: a national cohort and co-sibling study. Ann. Epidemiol. 86, 119–125.e4 (2023).

4. Allen, L., Leon-Attia, O., Shaham, M., Shefer, S. & Gabis, L. V. Autism risk linked to prematurity is more accentuated in girls. PLoS ONE 15, e0236994 (2020).

5. Crump, C., Sundquist, J. & Sundquist, K. Preterm or early term birth and risk of autism. Pediatrics 148, (2021).

6. Husby, A., Wohlfahrt, J. & Melbye, M. Gestational age at birth and cognitive outcomes in adolescence: population based full sibling cohort study. BMJ 380, e072779 (2023).

7. van Houdt, C. A., Oosterlaan, J., van Wassenaer-Leemhuis, A. G., van Kaam, A. H. & Aarnoudse-Moens, C. S. H. Executive function deficits in children born preterm or at low birthweight: a meta-analysis. Dev. Med. Child Neurol. 61, 1015–1024 (2019).

8. Schnider, B. et al. Executive function deficits mediate the association between very preterm birth and behavioral problems at school-age. Early Hum. Dev. 146, 105076 (2020).

9. Johnson, S. & Marlow, N. Preterm birth and childhood psychiatric disorders. Pediatr. Res. 69, 11R–8R (2011).

10. Johnson, S. & Marlow, N. Early and long-term outcome of infants born extremely preterm. Arch. Dis. Child. 102, 97–102 (2017).

11. Tokariev, A. et al. Preterm birth changes networks of newborn cortical activity. Cereb. Cortex 29, 814–826 (2019).

12. Chini, M. & Hanganu-Opatz, I. L. Prefrontal Cortex Development in Health and Disease: Lessons from Rodents and Humans. Trends Neurosci. 44, 227–240 (2021).

13. Ball, G. et al. The effect of preterm birth on thalamic and cortical development. Cereb. Cortex 22, 1016–1024 (2012).

14. Nosarti, C. et al. Preterm birth and structural brain alterations in early adulthood. Neuroimage Clin. 6, 180–191 (2014).

15. Wehrle, F. M. et al. Multimodal assessment shows misalignment of structural and functional thalamocortical connectivity in children and adolescents born very preterm. Neuroimage 215, 116779 (2020).

16. Yrjölä, P., Stjerna, S., Palva, J. M., Vanhatalo, S. & Tokariev, A. Phase-Based Cortical Synchrony Is Affected by Prematurity. Cereb. Cortex 32, 2265–2276 (2022).

17. Lacaille, H., Vacher, C.-M. & Penn, A. A. Preterm birth alters the maturation of the gabaergic system in the human prefrontal cortex. Front. Mol. Neurosci. 14, 827370 (2021).

18. Lacaille, H. et al. Impaired interneuron development in a novel model of neonatal brain injury. eNeuro 6, (2019).

19. Panda, S. et al. Estrogen Treatment Reverses Prematurity-Induced Disruption in Cortical Interneuron Population. J. Neurosci. 38, 7378–7391 (2018).

20. Tibrewal, M. et al. Disruption of Interneuron Neurogenesis in Premature Newborns and Reversal with Estrogen Treatment. J. Neurosci. 38, 1100–1113 (2018).

21. Komitova, M. et al. Hypoxia-induced developmental delays of inhibitory interneurons are reversed by environmental enrichment in the postnatal mouse forebrain. J. Neurosci. 33, 13375–13387 (2013).

22. Moloney, R. A. et al. Ongoing effects of preterm birth on the dopaminergic and noradrenergic pathways in the frontal cortex and hippocampus of guinea pigs. Dev. Neurobiol. 84, 93–110 (2024).

23. Moloney, R. A., Palliser, H. K., Pavy, C. L., Shaw, J. C. & Hirst, J. J. Zuranolone therapy protects frontal cortex neurodevelopment and improves behavioral outcomes after preterm birth. Brain Behav. 14, e70009 (2024).

24. Gramatté, T. & Schmidt, J. The effect of early postnatal hypoxia on the development of locomotor activity in rats. Biomed. Biochim. Acta 45, 523–529 (1986).

25. Speiser, Z., Korczyn, A. D., Teplitzky, I. & Gitter, S. Hyperactivity in rats following postnatal anoxia. Behav. Brain Res. 7, 379–382 (1983).

26. van der Kooij, M. A. et al. Mild neonatal hypoxia-ischemia induces long-term motor- and cognitive impairments in mice. Brain Behav. Immun. 24, 850–856 (2010).

27. Domnick, N.-K. et al. Neonatal hypoxia-ischemia impairs juvenile recognition memory by disrupting the maturation of prefrontal-hippocampal networks. Exp. Neurol. 273, 202– 214 (2015).

28. Forbes, T. A. et al. Environmental enrichment ameliorates perinatal brain injury and promotes functional white matter recovery. Nat. Commun. 11, 964 (2020).

29. Korkhin, A., Zubedat, S., Aga-Mizrachi, S. & Avital, A. Developmental effects of environmental enrichment on selective and auditory sustained attention. Psychoneuroendocrinology 111, 104479 (2020).

30. Zheng, H. J. V. et al. Environmental enrichment sharpens sensory acuity by enhancing information coding in barrel cortex and premotor cortex. eNeuro 8, (2021).

31. Cancedda, L. et al. Acceleration of visual system development by environmental enrichment. J. Neurosci. 24, 4840–4848 (2004).

32. Schuch, C. P. et al. Early environmental enrichment affects neurobehavioral development and prevents brain damage in rats submitted to neonatal hypoxia-ischemia. Neurosci. Lett. 617, 101–107 (2016).

33. Cheong, J. L. Y., Burnett, A. C., Treyvaud, K. & Spittle, A. J. Early environment and long-term outcomes of preterm infants. J. Neural Transm. 127, 1–8 (2020).

34. Allen, T. A. et al. Imaging the spread of reversible brain inactivations using fluorescent muscimol. J. Neurosci. Methods 171, 30–38 (2008).

35. Ingrao, J. C. et al. Aqueous stability and oral pharmacokinetics of meloxicam and carprofen in male C57BL/6 mice. J. Am. Assoc. Lab. Anim. Sci. 52, 553–559 (2013).

36. Guo, Z. V. et al. Procedures for behavioral experiments in head-fixed mice. PLoS ONE 9, e88678 (2014).

37. Hayar, A., Bryant, J. L., Boughter, J. D. & Heck, D. H. A low-cost solution to measure mouse licking in an electrophysiological setup with a standard analog-to-digital converter. J. Neurosci. Methods 153, 203–207 (2006).

38. Patel, T. P. et al. An open-source toolbox for automated phenotyping of mice in behavioral tasks. Front. Behav. Neurosci. 8, 349 (2014).

39. Witteveen, I. F. et al. Preterm birth accelerates the maturation of spontaneous and resting activity in the visual cortex. Front. Integr. Neurosci. 17, 1149159 (2023).

40. Jurjut, O., Georgieva, P., Busse, L. & Katzner, S. Learning Enhances Sensory Processing in Mouse V1 before Improving Behavior. J. Neurosci. 37, 6460–6474 (2017).

41. Liu, D. et al. Orbitofrontal control of visual cortex gain promotes visual associative learning. Nat. Commun. 11, 2784 (2020).

42. Murray, S. A. et al. Mouse gestation length is genetically determined. PLoS ONE 5, e12418 (2010).

43. Castillo-Ruiz, A., Hite, T. A., Yakout, D. W., Rosen, T. J. & Forger, N. G. Does birth trigger cell death in the developing brain? eNeuro 7, (2020).

44. Ribic, A., Crair, M. C. & Biederer, T. Synapse-Selective Control of Cortical Maturation and Plasticity by Parvalbumin-Autonomous Action of SynCAM 1. Cell Rep. 26, 381–393.e6 (2019).

45. Niell, C. M. & Stryker, M. P. Highly selective receptive fields in mouse visual cortex. J. Neurosci. 28, 7520–7536 (2008).

46. Berditchevskaia, A., Cazé, R. D. & Schultz, S. R. Performance in a GO/NOGO perceptual task reflects a balance between impulsive and instrumental components of behaviour. Sci. Rep. 6, 27389 (2016).

47. Del Rosario, J. et al. Diminished cortical excitation and elevated inhibition during perceptual impairments in a mouse model of autism. Cereb. Cortex 31, 3462–3474 (2021).

48. Goel, A. et al. Impaired perceptual learning in a mouse model of Fragile X syndrome is mediated by parvalbumin neuron dysfunction and is reversible. Nat. Neurosci. 21, 1404– 1411 (2018).

49. Poort, J. et al. Learning Enhances Sensory and Multiple Non-sensory Representations in Primary Visual Cortex. Neuron 86, 1478–1490 (2015).

50. Henschke, J. U. et al. Reward Association Enhances Stimulus-Specific Representations in Primary Visual Cortex. Curr. Biol. 30, 1866–1880.e5 (2020).

51. Corbo, J., McClure, J. P., Erkat, O. B. & Polack, P.-O. Dynamic Distortion of Orientation Representation after Learning in the Mouse Primary Visual Cortex. J. Neurosci. 42, 4311–4325 (2022).

52. Boughter, J. D., St John, S. J., Noel, D. T., Ndubuizu, O. & Smith, D. V. A brief-access test for bitter taste in mice. Chem. Senses 27, 133–142 (2002).

53. Durand, S. et al. A comparison of visual response properties in the lateral geniculate nucleus and primary visual cortex of awake and anesthetized mice. J. Neurosci. 36, 12144–12156 (2016).

54. Niell, C. M. & Stryker, M. P. Modulation of visual responses by behavioral state in mouse visual cortex. Neuron 65, 472–479 (2010).

55. Xing, D., Yeh, C.-I., Burns, S. & Shapley, R. M. Laminar analysis of visually evoked activity in the primary visual cortex. Proc Natl Acad Sci USA 109, 13871–13876 (2012).

56. Stringer, C., Michaelos, M., Tsyboulski, D., Lindo, S. E. & Pachitariu, M. High-precision coding in visual cortex. Cell 184, 2767–2778.e15 (2021).

57. Zhang, S. et al. Selective attention. Long-range and local circuits for top-down modulation of visual cortex processing. Science 345, 660–665 (2014).

58. Norman, K. J. et al. Post-error recruitment of frontal sensory cortical projections promotes attention in mice. Neuron 109, 1202–1213.e5 (2021).

59. Leinweber, M., Ward, D. R., Sobczak, J. M., Attinger, A. & Keller, G. B. A sensorimotor circuit in mouse cortex for visual flow predictions. Neuron 95, 1420–1432.e5 (2017).

60. Zhang, S. et al. Organization of long-range inputs and outputs of frontal cortex for top-down control. Nat. Neurosci. 19, 1733–1742 (2016).

61. Le Merre, P. et al. Reward-Based Learning Drives Rapid Sensory Signals in Medial Prefrontal Cortex and Dorsal Hippocampus Necessary for Goal-Directed Behavior. Neuron 97, 83–91.e5 (2018).

62. Peters, A. J., Marica, A.-M., Fabre, J. M. J., Harris, K. D. & Carandini, M. Visuomotor learning promotes visually evoked activity in the medial prefrontal cortex. Cell Rep. 41, 111487 (2022).

63. Wal, A., Klein, F. J., Born, G., Busse, L. & Katzner, S. Evaluating visual cues modulates their representation in mouse visual and cingulate cortex. J. Neurosci. 41, 3531–3544 (2021).

64. Kim, J.-H., Ma, D.-H., Jung, E., Choi, I. & Lee, S.-H. Gated feedforward inhibition in the frontal cortex releases goal-directed action. Nat. Neurosci. 24, 1452–1464 (2021).

65. Tervo, D. G. R. et al. A designer AAV variant permits efficient retrograde access to projection neurons. Neuron 92, 372–382 (2016).

66. Lima, S. Q., Hromádka, T., Znamenskiy, P. & Zador, A. M. PINP: a new method of tagging neuronal populations for identification during in vivo electrophysiological recording. PLoS ONE 4, e6099 (2009).

67. Cardin, J. A. Dissecting local circuits in vivo: integrated optogenetic and electrophysiology approaches for exploring inhibitory regulation of cortical activity. J. Physiol. Paris 106, 104–111 (2012).

68. Vormstein-Schneider, D. et al. Viral manipulation of functionally distinct interneurons in mice, non-human primates and humans. Nat. Neurosci. 23, 1629–1636 (2020).

69. Courcelles, E. J. et al. Association cortical areas in the mouse contain a large population of fast-spiking GABAergic neurons that do not express parvalbumin. Eur. J. Neurosci. 59, 3236–3255 (2024).

70. Le Merre, P., Ährlund-Richter, S. & Carlén, M. The mouse prefrontal cortex: Unity in diversity. Neuron 109, 1925–1944 (2021).

71. Jeong, J. H. & Choi, J.-S. Population analyses reveal heterogenous encoding in the medial prefrontal cortex during naturalistic foraging. (2025) doi:10.7554/eLife.93994.2.

72. Lui, J. H. et al. Differential encoding in prefrontal cortex projection neuron classes across cognitive tasks. Cell 184, 489–506.e26 (2021).

73. Hanley, J. A. & McNeil, B. J. The meaning and use of the area under a receiver operating characteristic (ROC) curve. Radiology 143, 29–36 (1982).

74. Réveillon, M., Hüppi, P. S. & Barisnikov, K. Inhibition difficulties in preterm children: Developmental delay or persistent deficit? Child Neuropsychol. 24, 734–762 (2018).

75. Nieves, G. M., & Liston, C., Divergent reward cue representations in the prefrontal cortex drive reward motivation in adolescence and adulthood. BioRxiv (2023) doi:10.1101/2023.11.07.565069.

76. Orso, R. et al. Early environmental enrichment rescues memory impairments provoked by mild neonatal hypoxia-ischemia in adolescent mice. Behav. Brain Res. 407, 113237 (2021).

77. Bartoletti, A., Medini, P., Berardi, N. & Maffei, L. Environmental enrichment prevents effects of dark-rearing in the rat visual cortex. Nat. Neurosci. 7, 215–216 (2004).

78. Akhund-Zade, J., Ho, S., O’Leary, C. & de Bivort, B. The effect of environmental enrichment on behavioral variability depends on genotype, behavior, and type of enrichment. J. Exp. Biol. 222, (2019).

79. Martínez-Cué, C. et al. Differential effects of environmental enrichment on behavior and learning of male and female Ts65Dn mice, a model for Down syndrome. Behav. Brain Res. 134, 185–200 (2002).

80. Stolp, H. B. et al. Interneuron development is disrupted in preterm brains with diffuse white matter injury: observations in mouse and human. Front. Physiol. 10, 955 (2019).

81. Parnaudeau, S. et al. Mediodorsal thalamus hypofunction impairs flexible goal-directed behavior. Biol. Psychiatry 77, 445–453 (2015).

82. Marton, T. F., Seifikar, H., Luongo, F. J., Lee, A. T. & Sohal, V. S. Roles of prefrontal cortex and mediodorsal thalamus in task engagement and behavioral flexibility. J. Neurosci. 38, 2569–2578 (2018).

83. Wimmer, R. D. et al. Thalamic control of sensory selection in divided attention. Nature 526, 705–709 (2015).

84. Mukherjee, A. et al. Variation of connectivity across exemplar sensory and associative thalamocortical loops in the mouse. eLife 9, (2020).

85. Delevich, K., Tucciarone, J., Huang, Z. J. & Li, B. The mediodorsal thalamus drives feedforward inhibition in the anterior cingulate cortex via parvalbumin interneurons. J. Neurosci. 35, 5743–5753 (2015).

86. Molnár, Z. & Rutherford, M. Brain maturation after preterm birth. Sci. Transl. Med. 5, 168ps2 (2013).

87. Toda, T. et al. Birth regulates the initiation of sensory map formation through serotonin signaling. Dev. Cell 27, 32–46 (2013).

88. Leung, M. P., Thompson, B., Black, J., Dai, S. & Alsweiler, J. M. The effects of preterm birth on visual development. Clin. Exp. Optom. 101, 4–12 (2018).

89. Hunt, B. A. E. et al. Disrupted visual cortex neurophysiology following very preterm birth. Biol. Psychiatry Cogn. Neurosci. Neuroimaging (2019) doi:10.1016/j.bpsc.2019.08.012.

90. Emberson, L. L., Boldin, A. M., Riccio, J. E., Guillet, R. & Aslin, R. N. Deficits in Top-Down Sensory Prediction in Infants At Risk due to Premature Birth. Curr. Biol. 27, 431– 436 (2017).

91. Boldin, A. M., Geiger, R. & Emberson, L. L. The emergence of top-down, sensory prediction during learning in infancy: A comparison of full-term and preterm infants. Dev. Psychobiol. 60, 544–556 (2018).

92. Jaffe-Dax, S., Boldin, A. M., Daw, N. D. & Emberson, L. L. A Computational Role for Top-Down Modulation from Frontal Cortex in Infancy. J. Cogn. Neurosci. 32, 508–514 (2020).

93. Siegle, J. H. et al. Reconciling functional differences in populations of neurons recorded with two-photon imaging and electrophysiology. eLife 10, (2021).

94. Wei, Z. et al. A comparison of neuronal population dynamics measured with calcium imaging and electrophysiology. PLoS Comput. Biol. 16, e1008198 (2020).

95. Kerstjens, J. M., de Winter, A. F., Bocca-Tjeertes, I. F., Bos, A. F. & Reijneveld, S. A. Risk of developmental delay increases exponentially as gestational age of preterm infants decreases: a cohort study at age 4 years. Dev. Med. Child Neurol. 54, 1096–1101 (2012).

96. Pierrat, V. et al. Neurodevelopmental outcome at 2 years for preterm children born at 22 to 34 weeks’ gestation in France in 2011: EPIPAGE-2 cohort study. BMJ 358, j3448 (2017).

97. Tang, Y. P., Wang, H., Feng, R., Kyin, M. & Tsien, J. Z. Differential effects of enrichment on learning and memory function in NR2B transgenic mice. Neuropharmacology 41, 779–790 (2001).

98. Imperio, C. G. et al. Exposure to environmental enrichment attenuates addiction-like behavior and alters molecular effects of heroin self-administration in rats. Neuropharmacology 139, 26–40 (2018).

99. Jendryka, M. M. et al. Control of sustained attention and impulsivity by Gq-protein signalling in parvalbumin interneurons of the anterior cingulate cortex. Transl. Psychiatry 13, 243 (2023).

100. Cardin, J. A. et al. Targeted optogenetic stimulation and recording of neurons in vivo using cell-type-specific expression of Channelrhodopsin-2. Nat. Protoc. 5, 247–254 (2010).

101. Bitzenhofer, S. H., Pöpplau, J. A., Chini, M., Marquardt, A. & Hanganu-Opatz, I. L. A transient developmental increase in prefrontal activity alters network maturation and causes cognitive dysfunction in adult mice. Neuron 109, 1350–1364.e6 (2021).

102. Colonnese, M. T. et al. A conserved switch in sensory processing prepares developing neocortex for vision. Neuron 67, 480–498 (2010).

103. Salmaso, N., Jablonska, B., Scafidi, J., Vaccarino, F. M. & Gallo, V. Neurobiology of premature brain injury. Nat. Neurosci. 17, 341–346 (2014).

