## Supplementary figures and images for "Divergent Representation and Processing of Task Cues in Sensory and Prefrontal Cortices of Preterm-Born Mice"

### Supplemental Figure 1

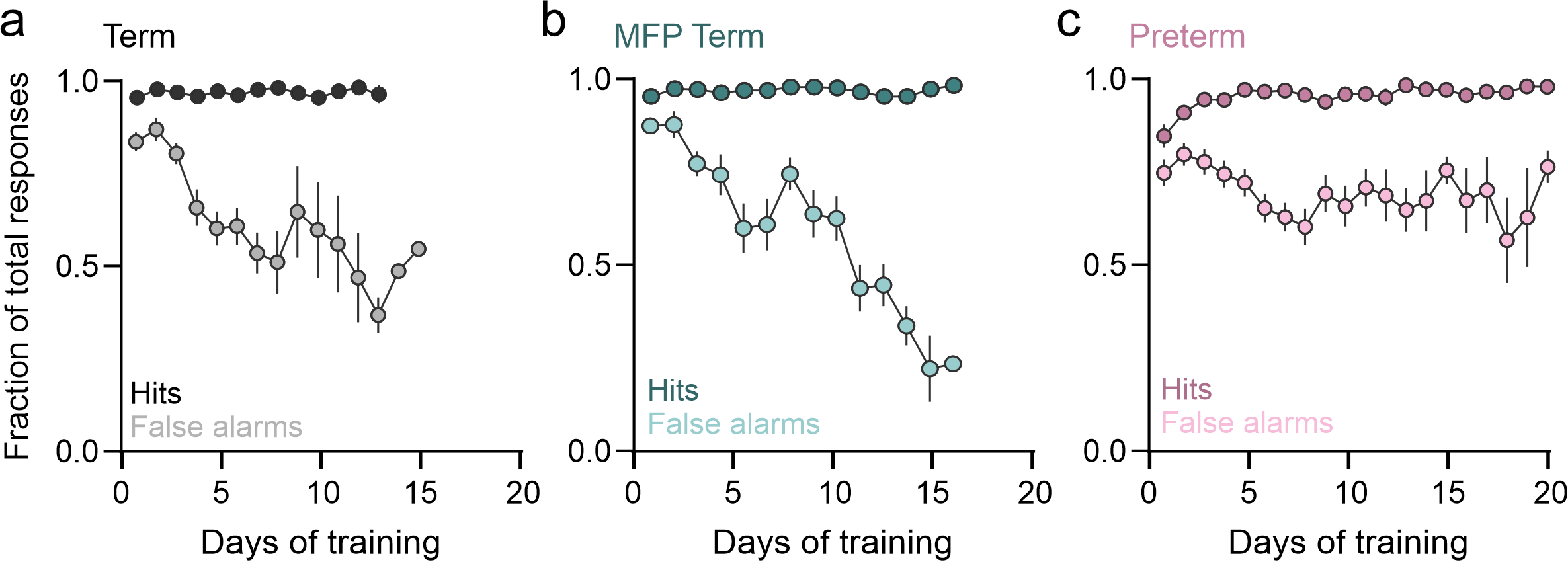

### Supplemental Figure 2

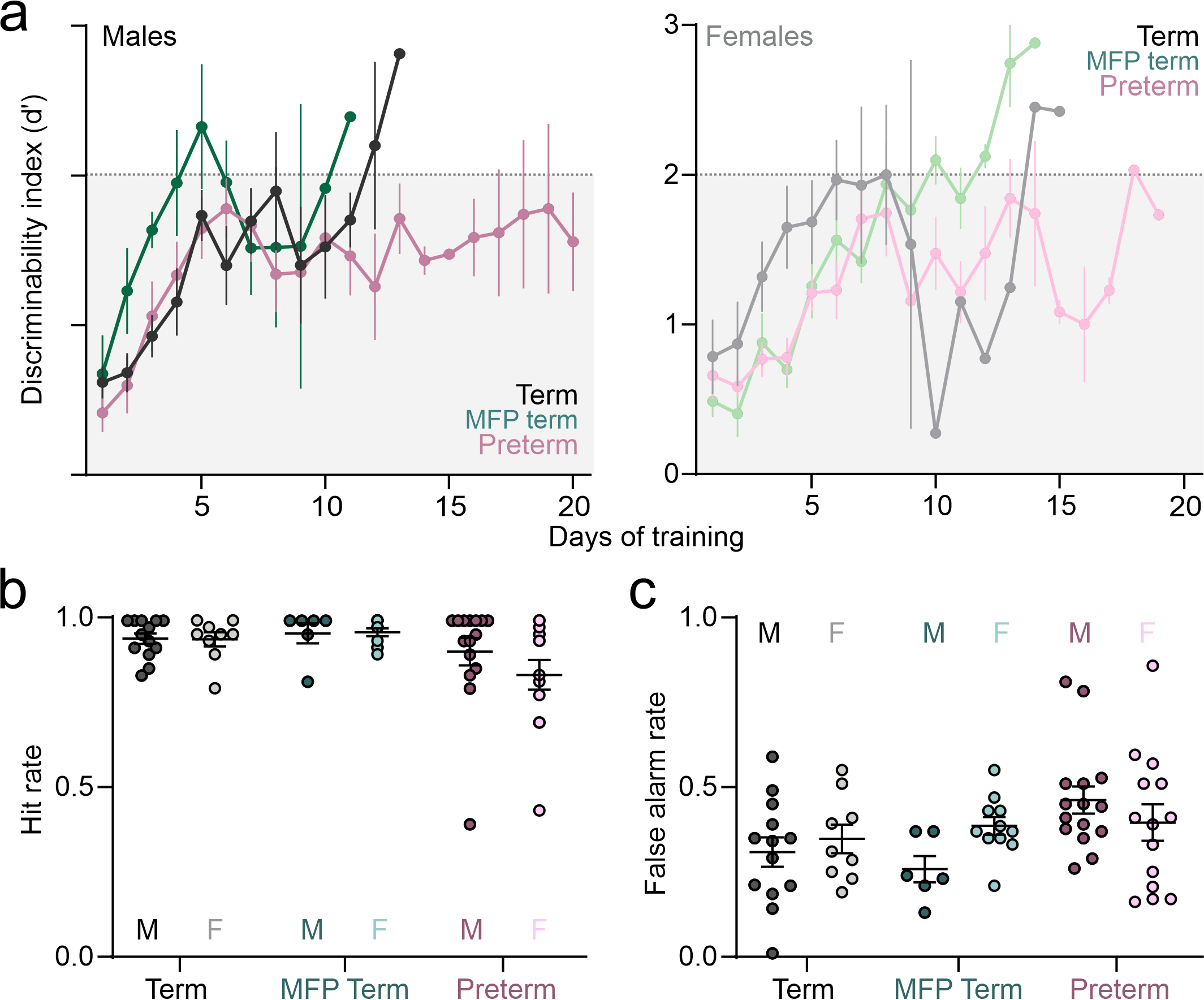

### Supplemental Figure 3

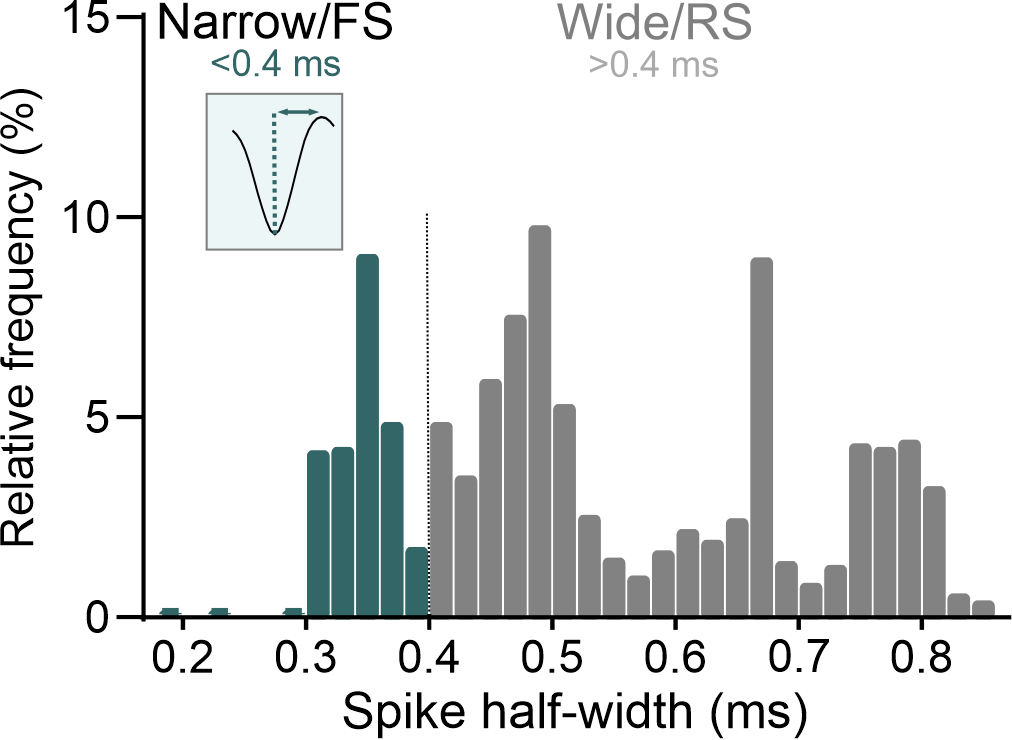

### Supplemental Figure 4

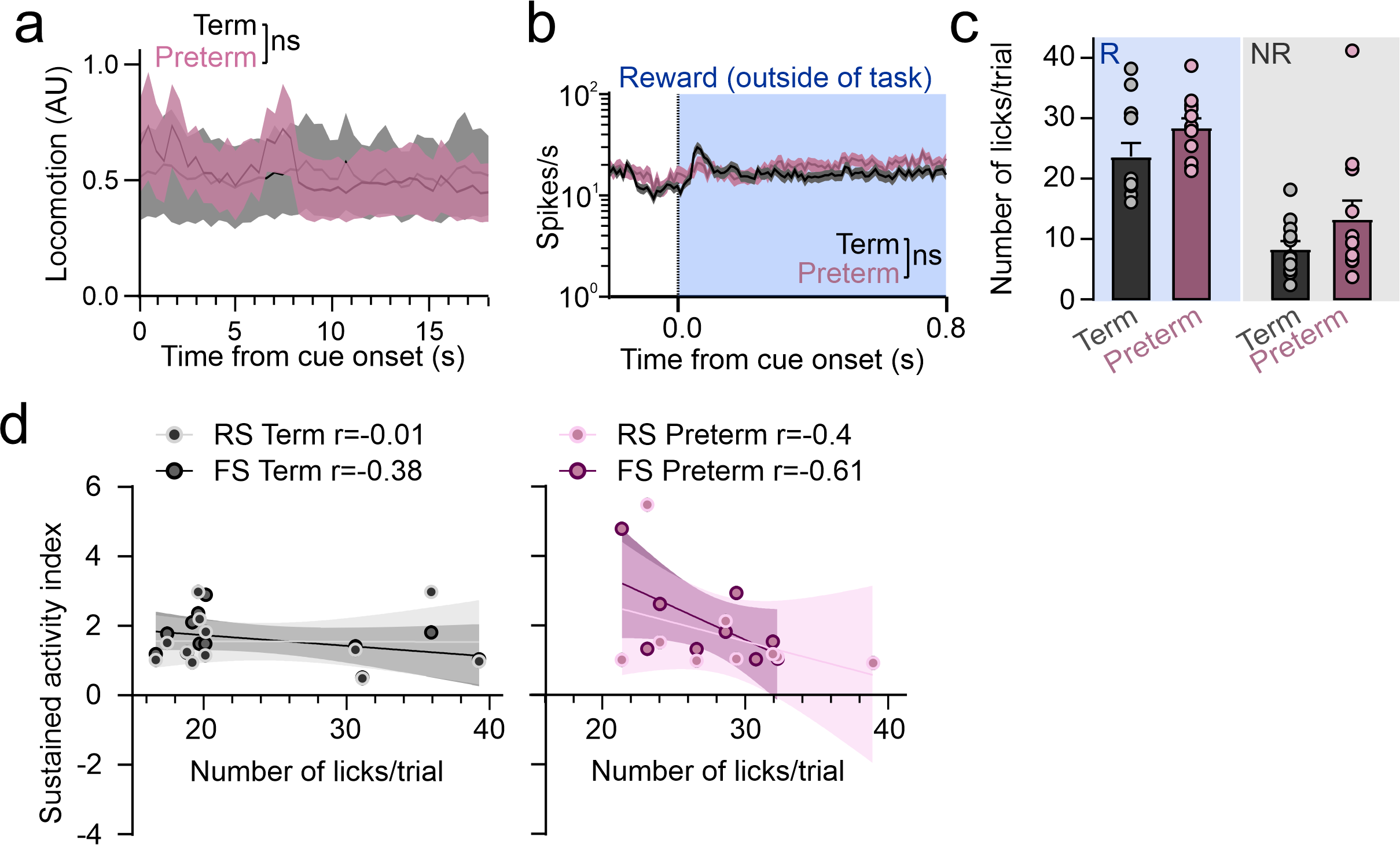

### Supplemental Figure 5

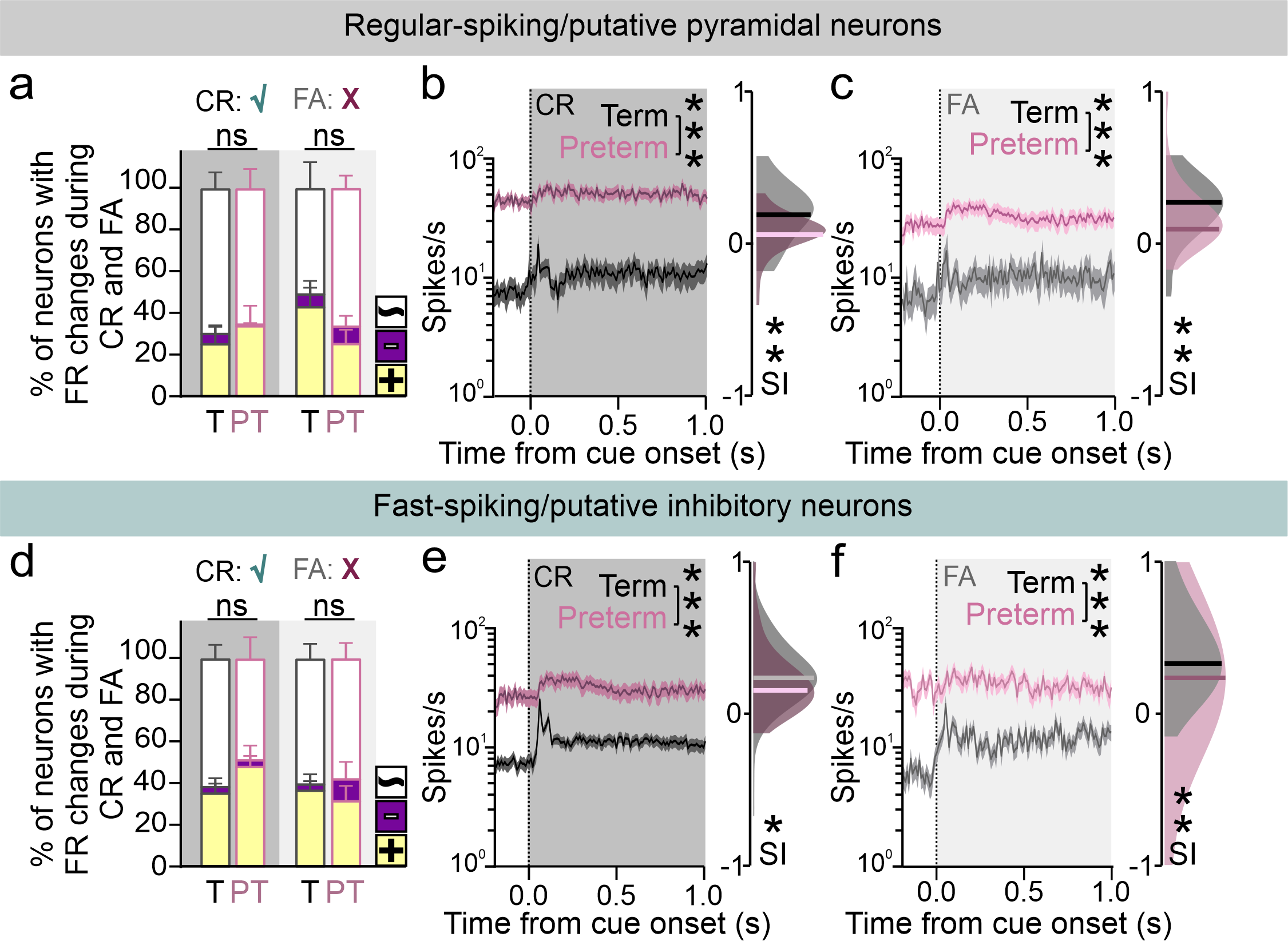

### Supplemental Figure 6

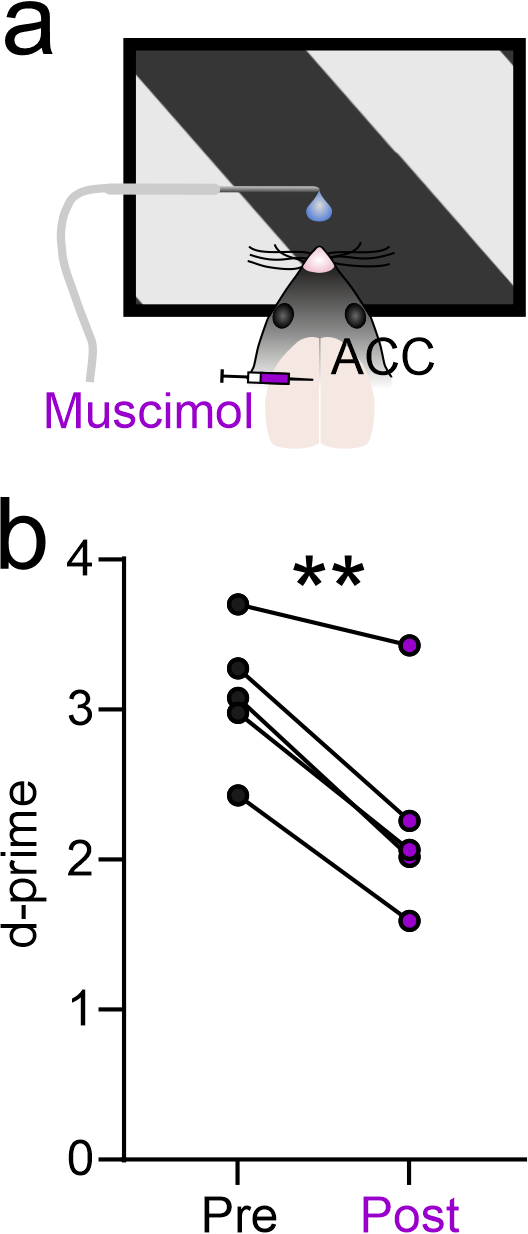

### Supplemental Figure 7

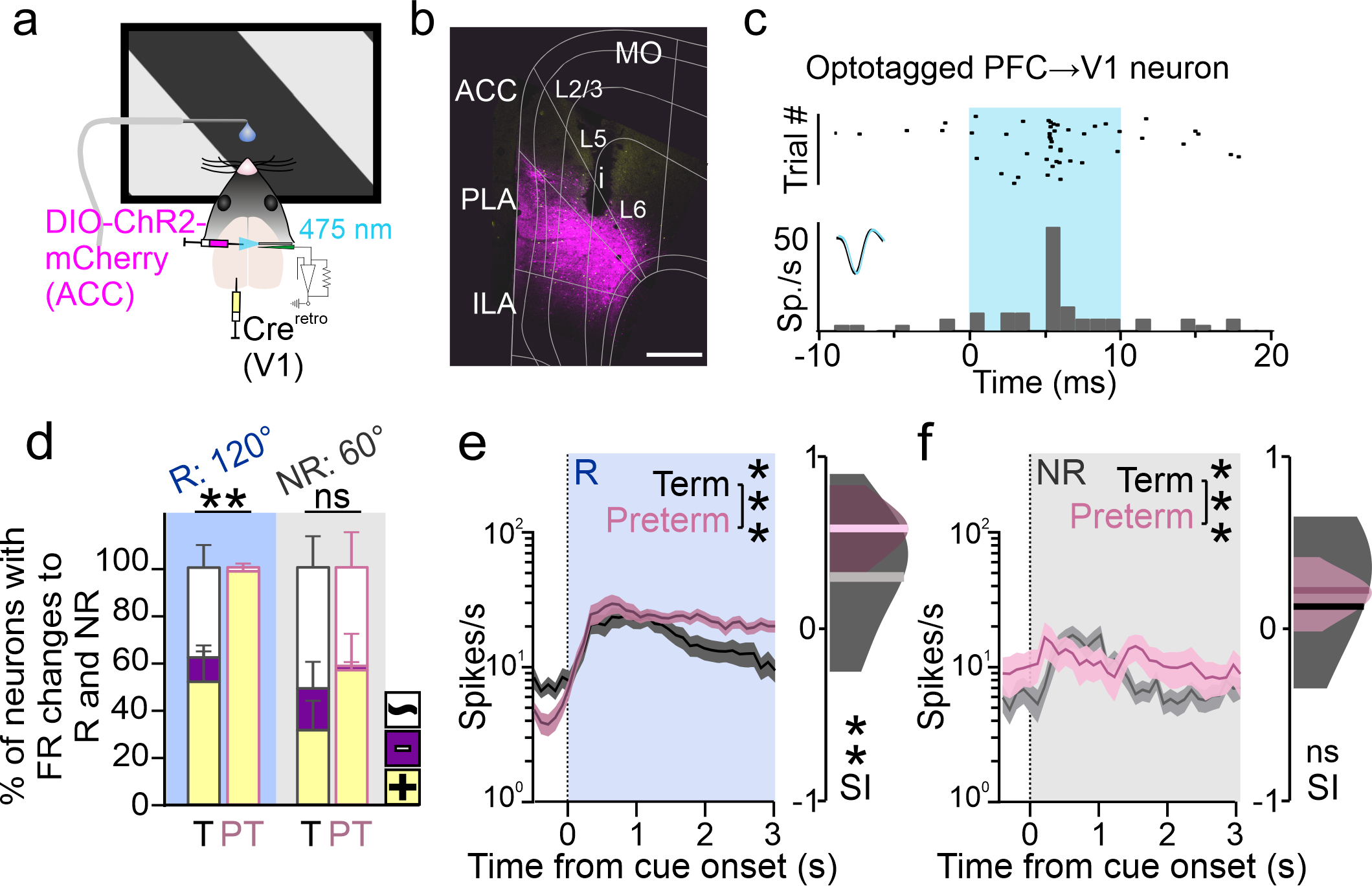

### Supplemental Figure 8

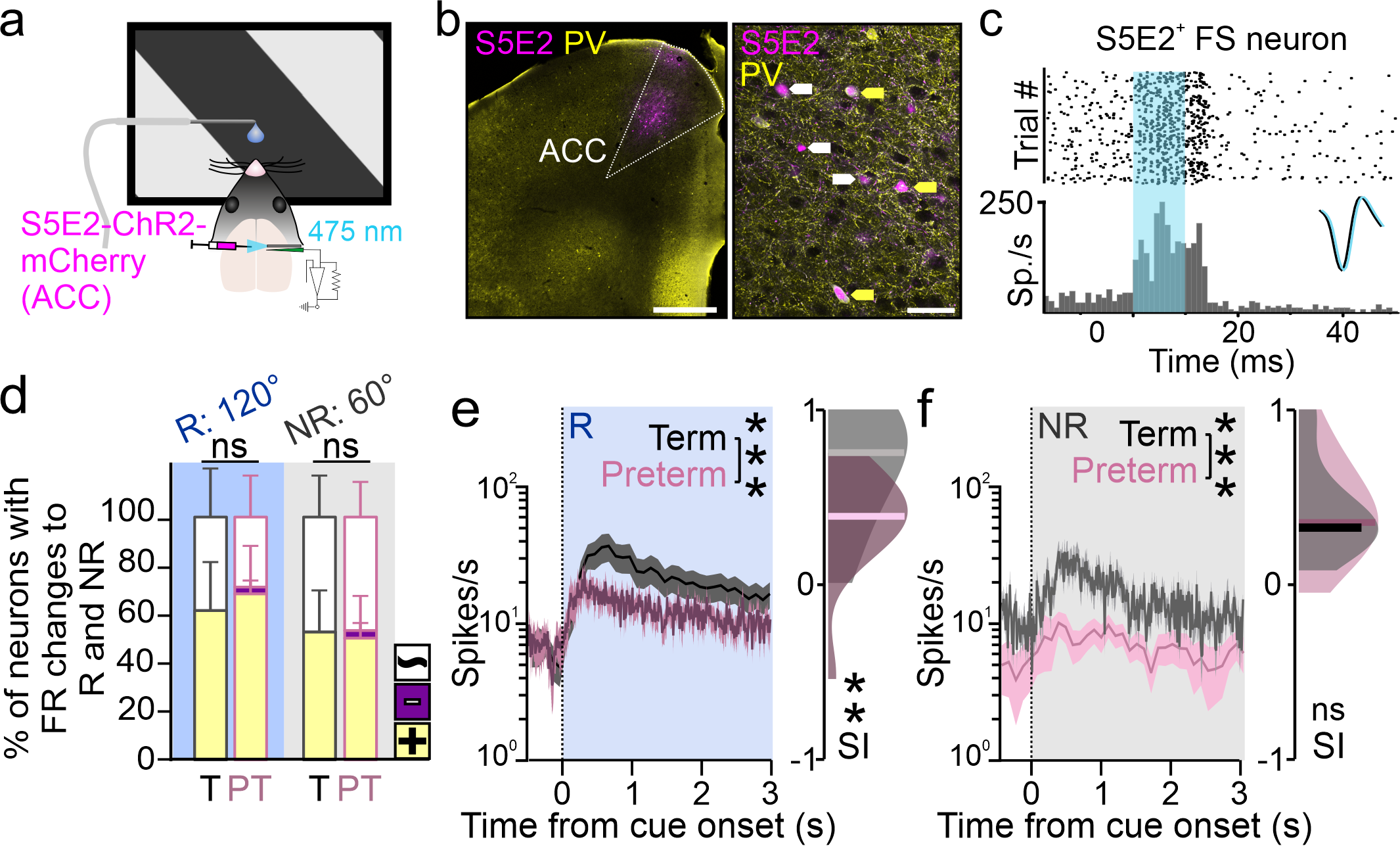

### Supplemental Figure 9

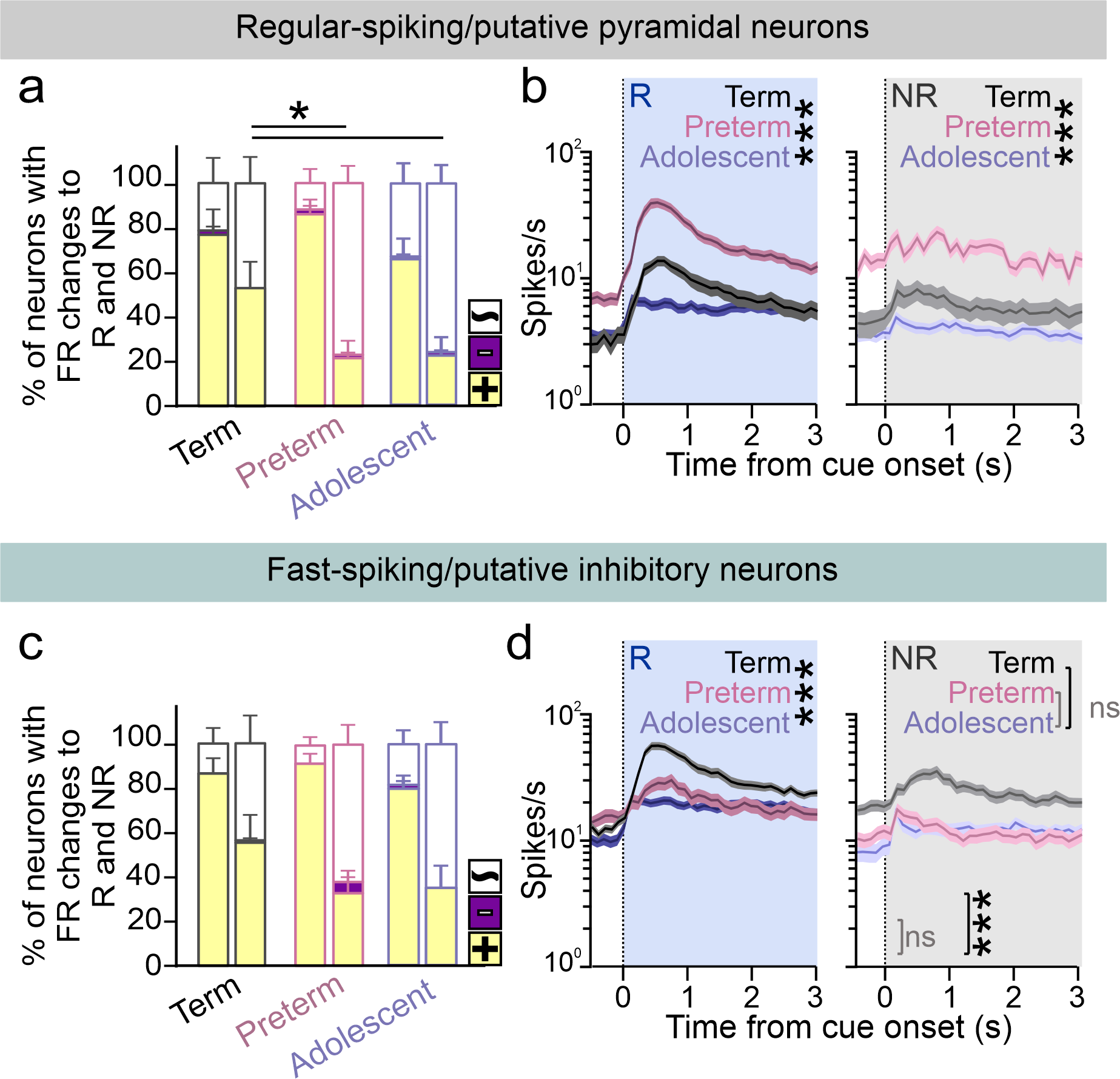
