## Supplemental Figure Legend for "Divergent Representation and Processing of Task Cues in Sensory and Prefrontal Cortices of Preterm-Born Mice"

**Supplementary Figure 1. The increase in task performance is driven by a reduction in false alarms.** Hit rates and False Alarm rates for term and preterm mice during training. Data are represented as mean±SEM of mice from Figure 2. Note the steady Hit Rates in both groups of term mice compared to preterm mice, as well as a precipitous decline in False Alarms in both groups of term mice that is absent from preterm mice.

**Supplementary Figure 2. Preterm birth has no-sex-specific effects on performance. a)** Learning trajectories of male (left) and female (right) mice. b) Hit Rates and c) False Alarm rates in trained male and female mice from all groups. Hits: birth x sex p=0.48, F_(2,62)_=0.74; sex p=0.42, birth p=0.02; False Alarms: birth x sex p=0.12, F_(2,62)_=2.2; sex p=0.38, birth p=0.03. Ns are indicated, data in (a) represents mean±SEM.

**Supplementary Figure 3.** Histogram of all recorded neurons in the study depicting multimodal distribution of spike widths (Durand et al., 2016). Cutoff for separation of FS and RS is indicated.

**Supplementary Figure 4. Elevated firing of V1 neurons is not related to increased locomotion or licking during task. a)** Treadmill motion in term and preterm mice measured during the task revealed no significant differences between term and preterm mice. Data represents averaged waveforms of motion recorded through piezo detectors with cues as triggers (permutation test of AUCs). **b)** PSTHs of V1 neurons significantly modulated by the rewarded cue outside of task context revealed no differences between term and preterm mice (N=111 units from 12 term mice and N=53 units from 10 preterm mice). Data are represented as mean±SEM. **c)** Number of licks to the rewarded and non-rewarded cues was significantly elevated in preterm mice (# of licks per bout Term_R_=23.65±2.23, Preterm_R_=28.48±1.49, Term_NR_=8.37±1.28, Preterm_NR_=13.31±3.056; ANOVA: birth x cue p=0.97, F_(1, 22)_=0.00065; birth p=0.024, F=5.85; cue p<0.0001, F=46.26); N=12 term and 10 preterm mice). **d)** Sustained activity index of V1 neurons and lick numbers for term (left) and preterm mice (right) are not significantly correlated (Pearson’s correlation, TermRS p=0.97, TermFS p=0.22, PretermRS p=0.28, PretermFS p=0.08).

**Supplementary Figure 5.** **Representation of behavioral outcomes is not impaired in V1 of preterm mice. a)** Fractions of RS neurons significantly modulated by the non-rewarded cue during Correct Rejections and False Alarms are not different between term and preterm mice (ANOVA cue x modulation: Correct Rejection p=0.24, F_(2,38)_=1.47; False Alarms p=0.7, F=0.36). Data represents mean±SEM of % of positively modulated/yellow, negatively modulated/magenta and unmodulated/white neurons. **b-c)** PSTHs of neurons significantly modulated by the non-rewarded cue during CR (b) and FA (c) trials show hyperactivity of RS neurons during both outcomes. Data represents mean±SEM. Permutation test of AUC, p=0.000. SI distributions demonstrate significant differences during both outcomes (K-S test, CR: D=0.48, p=0.0002; FA: D=0.37, p=0.02; medians are indicated with lines). **d)** Fractions of FS neurons significantly modulated by the non-rewarded cue during Correct Rejections and False Alarms are not different between term and preterm mice (ANOVA cue x modulation: Correct Rejection p=0.36, F=1.03; False Alarms p=0.75, F=0.28). Data represents mean±SEM of % of positively modulated/yellow, negatively modulated/magenta and unmodulated/white neurons. **e-f)** PSTHs of neurons significantly modulated by the non-rewarded cue during CR (e) and FA (f) trials show hyperactivity of FS neurons during both outcomes. Data represents mean±SEM. Permutation test of AUC, p=0.000. SI distributions demonstrate significant differences during both outcomes (K-S test, CR: D=0.23, p=0.009; FA: D=0.31, p=0.001; medians are indicated with lines). Ns are indicated in main text.

**Supplementary Figure 6. Pharmacological inactivation of the ACC impairs performance in trained mice. A)** GABA-agonist muscimol was injected into the ACC 1.5-2 hours before the training session began. **B)** Behavioral performance of trained mice before (black-filled circles) and after (violet-filled circles) muscimol injection. Pre-d’=3.05±0.28, post-d’=2.25±0.78; paired t-test p=0.02, t=4.42, df=3. N=mice.

**Supplementary Figure 7.** **V1-projecting PFC neurons of preterm mice are strongly driven by the rewarded cue. a)** PFC→V1 neurons were labeled using AAV-Cre^retro^ and DIO-ChR2-mCherry and their activity recorded during task performance (left). **b)** Representative image of labeled neurons (magenta) with injection site (i) and PFC subregions indicated. Scale bar: 250 µm. **c)** Representative raster and PSTH of an optotagged PFC→V1 neuron, with robust, regular and short latency firing in response to a 10 ms pulse of blue (473 nm) laser (indicated with blue rectangle). **d)** Rewarded cue activates almost all PFC→V1 neurons in preterm mice (ANOVA; Reward: birth x modulation, p=0.0017, F(2,26)=8.27, No Reward: birth x modulation, p=0.38, F=0.99; data represents mean±SEM of % of positively modulated/yellow, negatively modulated/magenta and unmodulated/white neurons; Ns are indicated in main text). **e-f)** PSTHs of neurons significantly modulated by task cues revealed elevated firing of PFC→V1 neurons in response to the rewarded cue in preterm mice (e) and irregular firing in response to the non-rewarded cue (f). Permutation test of PSTH AUCs, p=0.000. SI values of PFC→V1 neurons in preterm mice were significantly shifted towards higher values, with no significant differences for the non-rewarded cue (K-S test; Reward D=0.79, p=3.15x10^-6^; No Reward D=0.41, p=0.06). Medians are indicated with lines. PSTHs are presented as mean±SEM with Ns are indicated in main text.

**Supplementary Figure 8. The activity of optogenetically identified prefrontal FS overlaps with the activity of FS neurons isolated through spike sorting of bulk recordings.** **a)** FS neurons were labeled using S5E2-ChR2-mCherry and their activity recorded during task performance. **b)** Representative image of labeled neurons (magenta) with ACC indicated. Sections were counterstained with anti-Parvalbumin (PV) antibody (yellow). Yellow arrows indicate neurons with overlapping signals, white arrows indicate neurons with only mCherry fluorescence. Scale bars: 400 and 100 µm. **c)** Representative raster and PSTH of an optotagged S5E2^+^ FS neuron, with robust, rhythmic firing in response to a 10 ms pulse of blue (473 nm) laser (indicated with blue rectangle). **d)** Representation of task cues is not impaired in prefrontal neurons of preterm mice (ANOVA; Reward: birth x modulation: p=0.91, F_(2 14)_=0.094; No Reward: p=0.99, F=0.014; data represents mean±SEM of % of positively modulated/yellow, negatively modulated/magenta and unmodulated/white neurons; Ns are indicated in main text). **e-f)** PSTHs of neurons significantly modulated by task cues revealed substantially reduced cue-evoked activity in preterm mice (p=0.000, permutation test of PSTH AUCs). SI for the rewarded cue was shifted towards lower values (K-S test: D=0.79, p=0.000003). Shift in SI vales for the non-rewarded cue was marginally significant (D=0.41, p=0.057). Medians are indicated with lines. PSTHs are presented as mean±SEM with Ns are indicated in main text.

**Supplementary Figure 9.** Representation of cues in adult term, adult preterm and adolescent term mice depicted side by side for comparison. a-b) Cue representation and PSTHs of RS neurons (ANOVA; modulation x group Reward p=0.28, F_(4, 50)_=1.3; No Reward p=0.01, F=3.4; CR p=0.37, F=1.08; FA p=0.19, F=1.57). c-d) Cue representation and PSTHs of FS neurons. (ANOVA; modulation x group Reward p=0.29, F_(4, 50)_=1.27; No Reward p=0.56, F=0.75; CR p=0.6, F=0.68; FA p=0.23, F=1.4). PSTHs: permutation tests of AUCs, p values are indicated (***, p<0.0001). Data in (a) and (c) represents mean±SEM of % of positively modulated/yellow, negatively modulated/magenta and unmodulated/white neurons; PSTHs represent mean±SEM of neurons significantly modulated by task cues; Ns are indicated in main text.
