## Supplemental Tables for "Divergent Representation and Processing of Task Cues in Sensory and Prefrontal Cortices of Preterm-Born Mice"

**Supplementary Tables 1. LMM for Figure 2e**

| ***ANOVA Summary*** | | | | | | | | |
| --- | --- | --- | --- | --- | --- | --- | --- | --- |
| Effect | | df | | ChiSq | | p | | VS-MPR* |
| Intercept |  | 1 |  | 89.080 |  | 3.791×10^-21^ |  | 2.064×10^+18^ |
| Group |  | 2 |  | 9.485 |  | 0.009 |  | 8.899 |
| *Note.*  The following variables are used as random effects grouping factors: 'Subject', 'Training Day'. | | | | | | | | |
| *Note.*  Type III Sum of Squares | | | | | | | | |
| * Vovk-Sellke Maximum *p* -Ratio: Based on a two-sided *p* -value, the maximum possible odds in favor of H₁ over H₀ equals 1/(-e *p* log(*p* )) for *p* ≤ .37 (Sellke, Bayarri, & Berger, 2001). | | | | | | | | |

| ***Fit statistics*** | | | | | | | | |
| --- | --- | --- | --- | --- | --- | --- | --- | --- |
| Deviance | | log Lik. | | df | | AIC | | BIC |
| 2660.184 |  | -1330.092 |  | 12 |  | 2684.184 |  | 2745.154 |

| ***Fixed Effects Estimates*** | | | | | | | | | | | | |
| --- | --- | --- | --- | --- | --- | --- | --- | --- | --- | --- | --- | --- |
| Term | | Estimate | | SE | | df | | t | | p | | VS-MPR* |
| Intercept |  | 1.887 |  | 0.123 |  | 56.863 |  | 15.364 |  | 6.712×10^-22^ |  | 1.124×10^+19^ |
| Group (MFP Term) |  | 0.122 |  | 0.100 |  | 60.652 |  | 1.223 |  | 0.226 |  | 1.095 |
| Group (Preterm) |  | -0.292 |  | 0.089 |  | 57.401 |  | -3.267 |  | 0.002 |  | 31.819 |
| ***Estimated Marginal Means*** \| \| \| \| \| \| \| \| \| \| \|; \| \| \| \| \| \| \| \| 95% CI \| \| \|; Row \| \| Group \| \| Estimate \| \| SE \| \| Lower \| \| Upper \|; 1 \| \| MFP Term \| \| 2.009 \| \| 0.176 \| \| 1.664 \| \| 2.355 \|; 2 \| \| Preterm \| \| 1.595 \| \| 0.126 \| \| 1.347 \| \| 1.843 \|; 3 \| \| Term \| \| 2.057 \| \| 0.157 \| \| 1.749 \| \| 2.364 \| | | | | | | | | | | | | |

**Supplementary Tables 2. LMM for Figure 2i**

| ***ANOVA Summary*** | | | | | | | | |
| --- | --- | --- | --- | --- | --- | --- | --- | --- |
| Effect | | df | | ChiSq | | p | | VS-MPR* |
| Group |  | 2 |  | 4.548 |  | 0.103 |  | 1.572 |
| Trial block |  | 4 |  | 32.278 |  | 1.678×10^-6^ |  | 16483.591 |
| *Note.*  The following variable is used as a random effects grouping factor: 'Animal'. | | | | | | | | |
| *Note.*  Type III Sum of Squares | | | | | | | | |
| * Vovk-Sellke Maximum *p* -Ratio: Based on a two-sided *p* -value, the maximum possible odds in favor of H₁ over H₀ equals 1/(-e *p* log(*p* )) for *p* ≤ .37 (Sellke, Bayarri, & Berger, 2001). | | | | | | | | |

| ***Fit statistics*** | | | | | | | | |
| --- | --- | --- | --- | --- | --- | --- | --- | --- |
| Deviance | | log Lik. | | df | | AIC | | BIC |
| 2662.023 |  | -1331.012 |  | 9 |  | 2680.023 |  | 2714.105 |

| ***Fixed Effects Estimates*** | | | | | | | | | | | | |
| --- | --- | --- | --- | --- | --- | --- | --- | --- | --- | --- | --- | --- |
| Term | | Estimate | | SE | | df | | t | | p | | VS-MPR* |
| Intercept |  | 47.059 |  | 2.463 |  | 67.581 |  | 19.103 |  | 6.027×10^-29^ |  | 9.394×10^+25^ |
| Group (MFP Term) |  | -6.631 |  | 3.669 |  | 67.520 |  | -1.807 |  | 0.075 |  | 1.891 |
| Group (Preterm) |  | 6.414 |  | 3.291 |  | 67.688 |  | 1.949 |  | 0.055 |  | 2.293 |
| Trial block (1) |  | 3.638 |  | 1.171 |  | 257.827 |  | 3.106 |  | 0.002 |  | 28.321 |
| Trial block (2) |  | 2.938 |  | 1.171 |  | 257.827 |  | 2.508 |  | 0.013 |  | 6.617 |
| Trial block (3) |  | 1.922 |  | 1.171 |  | 257.827 |  | 1.641 |  | 0.102 |  | 1.580 |
| Trial block (4) |  | -2.860 |  | 1.201 |  | 258.038 |  | -2.382 |  | 0.018 |  | 5.104 |

| ***Estimated Marginal Means*** | | | | | | | | | | | | |
| --- | --- | --- | --- | --- | --- | --- | --- | --- | --- | --- | --- | --- |
|  | | | | | | | | | | 95% CI | | |
| Row | | Trial block | | Group | | Estimate | | SE | | Lower | | Upper |
| 1 |  | Block 1 |  | MFP Term |  | 44.066 |  | 4.849 |  | 34.562 |  | 53.570 |
| 2 |  | Block 2 |  | MFP Term |  | 43.366 |  | 4.849 |  | 33.862 |  | 52.870 |
| 3 |  | Block 3 |  | MFP Term |  | 42.350 |  | 4.849 |  | 32.846 |  | 51.855 |
| 4 |  | Block 4 |  | MFP Term |  | 37.567 |  | 4.863 |  | 28.036 |  | 47.099 |
| 5 |  | Block 5 |  | MFP Term |  | 34.790 |  | 4.883 |  | 25.219 |  | 44.361 |
| 6 |  | Block 1 |  | Preterm |  | 57.111 |  | 3.950 |  | 49.369 |  | 64.852 |
| 7 |  | Block 2 |  | Preterm |  | 56.411 |  | 3.950 |  | 48.669 |  | 64.152 |
| 8 |  | Block 3 |  | Preterm |  | 55.395 |  | 3.950 |  | 47.653 |  | 63.137 |
| 9 |  | Block 4 |  | Preterm |  | 50.612 |  | 3.974 |  | 42.823 |  | 58.402 |
| 10 |  | Block 5 |  | Preterm |  | 47.835 |  | 4.006 |  | 39.984 |  | 55.686 |
| 11 |  | Block 1 |  | Term |  | 50.914 |  | 4.413 |  | 42.265 |  | 59.563 |
| 12 |  | Block 2 |  | Term |  | 50.214 |  | 4.413 |  | 41.565 |  | 58.863 |
| 13 |  | Block 3 |  | Term |  | 49.198 |  | 4.413 |  | 40.549 |  | 57.848 |
| 14 |  | Block 4 |  | Term |  | 44.416 |  | 4.421 |  | 35.751 |  | 53.080 |
| 15 |  | Block 5 |  | Term |  | 41.638 |  | 4.446 |  | 32.924 |  | 50.353 |

**Supplementary Tables 3. LMM for Lick Frequencies During the First Session**

| ***ANOVA Summary*** | | | | | | | | |
| --- | --- | --- | --- | --- | --- | --- | --- | --- |
| Effect | | df | | ChiSq | | p | | VS-MPR* |
| Trial block |  | 4 |  | 10.413 |  | 0.034 |  | 3.198 |
| Group |  | 2 |  | 8.664 |  | 0.013 |  | 6.462 |
| *Note.*  The following variable is used as a random effects grouping factor: 'Animal'. | | | | | | | | |
| *Note.*  Type III Sum of Squares | | | | | | | | |
| * Vovk-Sellke Maximum *p* -Ratio: Based on a two-sided *p* -value, the maximum possible odds in favor of H₁ over H₀ equals 1/(-e *p* log(*p* )) for *p* ≤ .37 (Sellke, Bayarri, & Berger, 2001). | | | | | | | | |

| ***Fit statistics*** | | | | | | | | |
| --- | --- | --- | --- | --- | --- | --- | --- | --- |
| Deviance | | log Lik. | | df | | AIC | | BIC |
| 3111.352 |  | -1555.676 |  | 9 |  | 3129.352 |  | 3163.489 |

| ***Fixed Effects Estimates*** | | | | | | | | | | | | |
| --- | --- | --- | --- | --- | --- | --- | --- | --- | --- | --- | --- | --- |
| Term | | Estimate | | SE | | df | | t | | p | | VS-MPR* |
| Intercept |  | 39.747 |  | 3.109 |  | 68.135 |  | 12.783 |  | 9.093×10^-20^ |  | 9.227×10^+16^ |
| Trial block (1) |  | 1.284 |  | 2.533 |  | 260.764 |  | 0.507 |  | 0.613 |  | 1.000 |
| Trial block (2) |  | 7.248 |  | 2.533 |  | 260.764 |  | 2.862 |  | 0.005 |  | 14.980 |
| Trial block (3) |  | -0.755 |  | 2.533 |  | 260.764 |  | -0.298 |  | 0.766 |  | 1.000 |
| Trial block (4) |  | -2.399 |  | 2.579 |  | 261.189 |  | -0.930 |  | 0.353 |  | 1.001 |
| Group (MFP Term) |  | 3.578 |  | 4.622 |  | 67.613 |  | 0.774 |  | 0.442 |  | 1.000 |
| Group (Preterm) |  | -12.174 |  | 4.154 |  | 68.217 |  | -2.931 |  | 0.005 |  | 14.870 |

| ***Estimated Marginal Means*** | | | | | | | | | | | | |
| --- | --- | --- | --- | --- | --- | --- | --- | --- | --- | --- | --- | --- |
|  | | | | | | | | | | 95% CI | | |
| Row | | Trial block | | Group | | Estimate | | SE | | Lower | | Upper |
| 1 |  | Block 1 |  | MFP Term |  | 44.608 |  | 6.443 |  | 31.981 |  | 57.235 |
| 2 |  | Block 2 |  | MFP Term |  | 50.573 |  | 6.443 |  | 37.946 |  | 63.200 |
| 3 |  | Block 3 |  | MFP Term |  | 42.570 |  | 6.443 |  | 29.943 |  | 55.197 |
| 4 |  | Block 4 |  | MFP Term |  | 40.926 |  | 6.461 |  | 28.262 |  | 53.589 |
| 5 |  | Block 5 |  | MFP Term |  | 37.946 |  | 6.508 |  | 25.191 |  | 50.701 |
| 6 |  | Block 1 |  | Preterm |  | 28.856 |  | 5.386 |  | 18.301 |  | 39.412 |
| 7 |  | Block 2 |  | Preterm |  | 34.821 |  | 5.386 |  | 24.265 |  | 45.376 |
| 8 |  | Block 3 |  | Preterm |  | 26.818 |  | 5.386 |  | 16.262 |  | 37.373 |
| 9 |  | Block 4 |  | Preterm |  | 25.174 |  | 5.424 |  | 14.543 |  | 35.804 |
| 10 |  | Block 5 |  | Preterm |  | 22.194 |  | 5.538 |  | 11.340 |  | 33.048 |
| 11 |  | Block 1 |  | Term |  | 49.627 |  | 5.937 |  | 37.989 |  | 61.264 |
| 12 |  | Block 2 |  | Term |  | 55.591 |  | 5.937 |  | 43.954 |  | 67.228 |
| 13 |  | Block 3 |  | Term |  | 47.588 |  | 5.937 |  | 35.951 |  | 59.226 |
| 14 |  | Block 4 |  | Term |  | 45.944 |  | 5.995 |  | 34.193 |  | 57.695 |
| 15 |  | Block 5 |  | Term |  | 42.965 |  | 6.082 |  | 31.045 |  | 54.885 |

**Supplementary Tables 3. LMM for Supplementary Figure 2**

| ***ANOVA Summary*** | | | | | | | | |
| --- | --- | --- | --- | --- | --- | --- | --- | --- |
| Effect | | df | | ChiSq | | p | | VS-MPR* |
| Sex |  | 1 |  | 5.367 |  | 0.021 |  | 4.612 |
| Group |  | 2 |  | 9.371 |  | 0.009 |  | 8.507 |
| *Note.*  The following variables are used as random effects grouping factors: 'Subject', 'Training Day'. | | | | | | | | |
| *Note.*  Type III Sum of Squares | | | | | | | | |
| * Vovk-Sellke Maximum *p* -Ratio: Based on a two-sided *p* -value, the maximum possible odds in favor of H₁ over H₀ equals 1/(-e *p* log(*p* )) for *p* ≤ .37 (Sellke, Bayarri, & Berger, 2001). ***Fit statistics*** \| \| \| \| \| \| \| \| \|; Deviance \| \| log Lik. \| \| df \| \| AIC \| \| BIC \|; 2651.980 \| \| -1325.990 \| \| 16 \| \| 2683.980 \| \| 2765.274 \| | | | | | | | | |
| ***Fixed Effects Estimates*** \| \| \| \| \| \| \| \| \| \| \| \| \|; Term \| \| Estimate \| \| SE \| \| df \| \| t \| \| p \| \| VS-MPR* \|; Intercept \| \| 1.895 \| \| 0.122 \| \| 58.435 \| \| 15.519 \| \| 2.167×10^-22^ \| \| 3.403×10^+19^ \|; Sex (Female) \| \| -0.161 \| \| 0.065 \| \| 60.721 \| \| -2.477 \| \| 0.016 \| \| 5.547 \|; Group (MFP Term) \| \| 0.189 \| \| 0.102 \| \| 61.673 \| \| 1.848 \| \| 0.069 \| \| 1.987 \|; Group (Preterm) \| \| -0.294 \| \| 0.089 \| \| 57.570 \| \| -3.313 \| \| 0.002 \| \| 35.724 \| | | | | | | |  |  |

| ***Estimated Marginal Means*** | | | | | | | | | | | | |
| --- | --- | --- | --- | --- | --- | --- | --- | --- | --- | --- | --- | --- |
|  | | | | | | | | | | 95% CI | | |
| Row | | Sex | | Group | | Estimate | | SE | | Lower | | Upper |
| 1 |  | Female |  | MFP Term |  | 1.923 |  | 0.177 |  | 1.575 |  | 2.270 |
| 2 |  | Male |  | MFP Term |  | 2.245 |  | 0.204 |  | 1.845 |  | 2.646 |
| 3 |  | Female |  | Preterm |  | 1.440 |  | 0.136 |  | 1.173 |  | 1.708 |
| 4 |  | Male |  | Preterm |  | 1.762 |  | 0.147 |  | 1.475 |  | 2.050 |
| 5 |  | Female |  | Term |  | 1.840 |  | 0.165 |  | 1.516 |  | 2.163 |
| 6 |  | Male |  | Term |  | 2.162 |  | 0.167 |  | 1.834 |  | 2.490 |

**Supplementary Tables 4. Linear Regression Model for Figure 4J**

| ***^Model Summary-d’^*** | | | | | | | | | | | | | | | | | | |
| --- | --- | --- | --- | --- | --- | --- | --- | --- | --- | --- | --- | --- | --- | --- | --- | --- | --- | --- |
|  | | | | | | | | | | | | | | Durbin-Watson | | | | |
| Model | | R | | R² | | Adjusted R² | | RMSE | | AIC | | BIC | | Autocorrelation | | Statistic | | p |
| M₀ |  | 0.000 |  | 0.000 |  | 0.000 |  | 0.485 |  | 15.460 |  | 15.854 |  | -0.165 |  | 2.128 |  | 0.841 |
| M₁ |  | 0.359 |  | 0.129 |  | -0.161 |  | 0.523 |  | 18.217 |  | 19.006 |  | -0.132 |  | 2.013 |  | 0.823 |
| *Note.*  M₁ includes SI R Term, SI NR Term | | | | | | | | | | | | | | | | | | |

| ***ANOVA*** | | | | | | | | | | | | |
| --- | --- | --- | --- | --- | --- | --- | --- | --- | --- | --- | --- | --- |
| Model | |  | | Sum of Squares | | df | | Mean Square | | F | | p |
| M₁ |  | Regression |  | 0.243 |  | 2 |  | 0.121 |  | 0.444 |  | 0.661 |
|  |  | Residual |  | 1.640 |  | 6 |  | 0.273 |  |  |  |  |
|  |  | Total |  | 1.883 |  | 8 |  |  |  |  |  |  |
| *Note.*  The intercept model is omitted, as no meaningful information can be shown. | | | | | | | | | | | | |

| *Coefficients* | | | | | | | | | | | | | | | | |
| --- | --- | --- | --- | --- | --- | --- | --- | --- | --- | --- | --- | --- | --- | --- | --- | --- |
|  | | | | | | | | | | | | | | Collinearity Statistics | | |
| Model | |  | | Unstandardized | | Standard Error | | Standardized | | t | | p | | Tolerance | | VIF |
| M₀ |  | (Intercept) |  | 2.560 |  | 0.162 |  |  |  | 15.832 |  | 2.534×10^-7^ |  |  |  |  |
| M₁ |  | (Intercept) |  | 2.708 |  | 0.639 |  |  |  | 4.234 |  | 0.005 |  |  |  |  |
|  |  | RS R ACC |  | -0.825 |  | 0.932 |  | -0.352 |  | -0.885 |  | 0.410 |  | 0.916 |  | 1.092 |
|  |  | RS NR SCC |  | 1.244 |  | 2.193 |  | 0.226 |  | 0.567 |  | 0.591 |  | 0.916 |  | 1.092 |

**Supplementary Tables 5. LMM for Figure 6b**

| ***ANOVA Summary*** | | | | | | | | |
| --- | --- | --- | --- | --- | --- | --- | --- | --- |
| Effect | | df | | ChiSq | | p | | VS-MPR* |
| Age 2 |  | 1 |  | 0.573 |  | 0.449 |  | 1.000 |
| *Note.*  Type III Sum of Squares | | | | | | | | |
| * Vovk-Sellke Maximum *p* -Ratio: Based on a two-sided *p* -value, the maximum possible odds in favor of H₁ over H₀ equals 1/(-e *p* log(*p* )) for *p* ≤ .37 (Sellke, Bayarri, & Berger, 2001). | | | | | | | | |

| ***Fit statistics*** | | | | | | | | |
| --- | --- | --- | --- | --- | --- | --- | --- | --- |
| Deviance | | log Lik. | | df | | AIC | | BIC |
| 479.632 |  | -239.816 |  | 7 |  | 493.632 |  | 517.997 |

| ***Fixed Effects Estimates*** | | | | | | | | | | | | |
| --- | --- | --- | --- | --- | --- | --- | --- | --- | --- | --- | --- | --- |
| Term | | Estimate | | SE | | df | | t | | p | | VS-MPR* |
| Intercept |  | 1.939 |  | 0.219 |  | 18.340 |  | 8.847 |  | 4.874×10^-8^ |  | 448279.161 |
| Age |  | 0.070 |  | 0.084 |  | 26.097 |  | 0.837 |  | 0.410 |  | 1.000 |

| ***Estimated Marginal Means*** | | | | | | | | |
| --- | --- | --- | --- | --- | --- | --- | --- | --- |
|  | | | | | | 95% CI | | |
| Age | | Estimate | | SE | | Lower | | Upper |
| Adolescent |  | 2.009 |  | 0.239 |  | 1.541 |  | 2.477 |
| Adult |  | 1.869 |  | 0.231 |  | 1.417 |  | 2.321 |

**Supplementary Tables 6. LMM for Figure 7b**

| ***ANOVA Summary*** | | | | | | |
| --- | --- | --- | --- | --- | --- | --- |
| Effect | | df | | ChiSq | | p |
| Enrichment |  | 1 |  | 17.514 |  | 2.852×10^-5^ |
| Birth |  | 1 |  | 17.035 |  | 3.670×10^-5^ |
| *Note.*  The following variables are used as random effects grouping factors: 'Subject', 'Training day'. | | | | | | |
| *Note.*  Type III Sum of Squares | | | | | | |

| ***Fit statistics*** | | | | | | | | |
| --- | --- | --- | --- | --- | --- | --- | --- | --- |
| Deviance | | log Lik. | | df | | AIC | | BIC |
| 1551.408 |  | -775.704 |  | 11 |  | 1573.408 |  | 1624.772 |

| ***Fixed Effects Estimates*** | | | | | | | | | | |
| --- | --- | --- | --- | --- | --- | --- | --- | --- | --- | --- |
| Term | | Estimate | | SE | | df | | t | | p |
| Intercept |  | 1.612 |  | 0.150 |  | 27.411 |  | 10.749 |  | 2.463×10^-11^ |
| Enrichment |  | 0.316 |  | 0.070 |  | 71.572 |  | 4.536 |  | 2.256×10^-5^ |
| Birth |  | -0.294 |  | 0.068 |  | 49.998 |  | -4.340 |  | 6.929×10^-5^ |

| ***Estimated Marginal Means*** | | | | | | | | | | | | |
| --- | --- | --- | --- | --- | --- | --- | --- | --- | --- | --- | --- | --- |
|  | | | | | | | | | | 95% CI | | |
| Row | | Enrichment | | Birth | | Estimate | | SE | | Lower | | Upper |
| 1 |  | No |  | Preterm |  | 1.634 |  | 0.138 |  | 1.364 |  | 1.904 |
| 2 |  | Yes |  | Preterm |  | 1.003 |  | 0.161 |  | 0.687 |  | 1.318 |
| 3 |  | No |  | Term |  | 2.222 |  | 0.192 |  | 1.846 |  | 2.597 |
| 4 |  | Yes |  | Term |  | 1.590 |  | 0.215 |  | 1.169 |  | 2.011 |

**Supplementary Tables 7. ANOVA for Figure 7d and e**

| ***Within Subjects Effects*** | | | | | | | | | | | | |
| --- | --- | --- | --- | --- | --- | --- | --- | --- | --- | --- | --- | --- |
| Cases | | Sphericity Correction | | Sum of Squares | | df | | Mean Square | | F | | p |
| Training |  | None |  | 0.080 |  | 1.000 |  | 0.080 |  | 2.940 |  | 0.091 |
| Training ✻ Birth |  | None |  | 0.036 |  | 1.000 |  | 0.036 |  | 1.326 |  | 0.254 |
| Training ✻ Enrichment |  | None |  | 0.777 |  | 1.000 |  | 0.777 |  | 28.491 |  | 1.205×10^-6^ |
| Training ✻ Birth ✻ Enrichment |  | None |  | 0.293 |  | 1.000 |  | 0.293 |  | 10.748 |  | 0.002 |
| Residuals |  | None |  | 1.827 |  | 67.000 |  | 0.027 |  |  |  |  |
| Trial type |  | None |  | 6.503 |  | 1.000 |  | 6.503 |  | 443.179 |  | 3.032×10^-31^ |
| Trial type ✻ Birth |  | None |  | 0.083 |  | 1.000 |  | 0.083 |  | 5.644 |  | 0.020 |
| Trial type ✻ Enrichment |  | None |  | 0.053 |  | 1.000 |  | 0.053 |  | 3.641 |  | 0.061 |
| Trial type ✻ Birth ✻ Enrichment |  | None |  | 5.773×10^-4^ |  | 1.000 |  | 5.773×10^-4^ |  | 0.039 |  | 0.843 |
| Residuals |  | None |  | 0.983 |  | 67.000 |  | 0.015 |  |  |  |  |
| Training ✻ Trial type |  | None |  | 3.557 |  | 1.000 |  | 3.557 |  | 262.116 |  | 7.541×10^-25^ |
| Training ✻ Trial type ✻ Birth |  | None |  | 0.051 |  | 1.000 |  | 0.051 |  | 3.727 |  | 0.058 |
| Training ✻ Trial type ✻ Enrichment |  | None |  | 0.007 |  | 1.000 |  | 0.007 |  | 0.543 |  | 0.464 |
| Training ✻ Trial type ✻ Birth ✻ Enrichment |  | None |  | 0.012 |  | 1.000 |  | 0.012 |  | 0.885 |  | 0.350 |
| Residuals |  | None |  | 0.909 |  | 67.000 |  | 0.014 |  |  |  |  |
| *Note.*  Sphericity corrections not available for factors with 2 levels. | | | | | | | | | | | | |
| *Note.*  Type III Sum of Squares | | | | | | | | | | | | |

**Post Hoc Tests**

| *Post Hoc Comparisons - Birth ✻ Enrichment ✻ Training ✻ Trial type* | | | | | | | | | | | | | |
| --- | --- | --- | --- | --- | --- | --- | --- | --- | --- | --- | --- | --- | --- |
|  | |  | | Mean Difference | | SE | | df | | t | | p_holm_ | |
| Preterm, ENR, First session, Hits |  | Term, ENR, First session, Hits |  | 0.152 |  | 0.064 |  | 67 |  | 2.387 |  | 0.674 |  |
|  |  | Preterm, STD, First session, Hits |  | -0.216 |  | 0.050 |  | 67 |  | -4.296 |  | 0.003 | ** |
|  |  | Term, STD, First session, Hits |  | -0.286 |  | 0.052 |  | 67 |  | -5.464 |  | 5.028×10^-5^ | *** |
|  |  | Preterm, ENR, Last session, Hits |  | -0.200 |  | 0.045 |  | 67 |  | -4.398 |  | 0.002 | ** |
|  |  | Term, ENR, Last session, Hits |  | -0.331 |  | 0.050 |  | 67 |  | -6.559 |  | 7.492×10^-7^ | *** |
|  |  | Preterm, STD, Last session, Hits |  | -0.332 |  | 0.045 |  | 67 |  | -7.322 |  | 3.449×10^-8^ | *** |
|  |  | Term, STD, Last session, Hits |  | -0.339 |  | 0.046 |  | 67 |  | -7.369 |  | 2.898×10^-8^ | *** |
|  |  | Preterm, ENR, First session, FAs |  | 0.055 |  | 0.047 |  | 67 |  | 1.184 |  | 1.000 |  |
|  |  | Term, ENR, First session, FAs |  | 0.189 |  | 0.080 |  | 67 |  | 2.365 |  | 0.690 |  |
|  |  | Preterm, STD, First session, FAs |  | -0.112 |  | 0.057 |  | 67 |  | -1.962 |  | 1.000 |  |
|  |  | Term, STD, First session, FAs |  | -0.130 |  | 0.061 |  | 67 |  | -2.140 |  | 1.000 |  |
|  |  | Preterm, ENR, Last session, FAs |  | 0.291 |  | 0.059 |  | 67 |  | 4.931 |  | 3.688×10^-4^ | *** |
|  |  | Term, ENR, Last session, FAs |  | 0.319 |  | 0.070 |  | 67 |  | 4.531 |  | 0.001 | ** |
|  |  | Preterm, STD, Last session, FAs |  | 0.220 |  | 0.053 |  | 67 |  | 4.159 |  | 0.005 | ** |
|  |  | Term, STD, Last session, FAs |  | 0.326 |  | 0.056 |  | 67 |  | 5.843 |  | 1.209×10^-5^ | *** |
| Term, ENR, First session, Hits |  | Preterm, STD, First session, Hits |  | -0.368 |  | 0.054 |  | 67 |  | -6.798 |  | 2.923×10^-7^ | *** |
|  |  | Term, STD, First session, Hits |  | -0.439 |  | 0.056 |  | 67 |  | -7.810 |  | 4.824×10^-9^ | *** |
|  |  | Preterm, ENR, Last session, Hits |  | -0.352 |  | 0.053 |  | 67 |  | -6.630 |  | 5.746×10^-7^ | *** |
|  |  | Term, ENR, Last session, Hits |  | -0.483 |  | 0.050 |  | 67 |  | -9.617 |  | 3.014×10^-12^ | *** |
|  |  | Preterm, STD, Last session, Hits |  | -0.484 |  | 0.050 |  | 67 |  | -9.759 |  | 1.706×10^-12^ | *** |
|  |  | Term, STD, Last session, Hits |  | -0.492 |  | 0.050 |  | 67 |  | -9.778 |  | 1.594×10^-12^ | *** |
|  |  | Preterm, ENR, First session, FAs |  | -0.097 |  | 0.077 |  | 67 |  | -1.254 |  | 1.000 |  |
|  |  | Term, ENR, First session, FAs |  | 0.037 |  | 0.052 |  | 67 |  | 0.708 |  | 1.000 |  |
|  |  | Preterm, STD, First session, FAs |  | -0.264 |  | 0.060 |  | 67 |  | -4.370 |  | 0.002 | ** |
|  |  | Term, STD, First session, FAs |  | -0.282 |  | 0.064 |  | 67 |  | -4.409 |  | 0.002 | ** |
|  |  | Preterm, ENR, Last session, FAs |  | 0.138 |  | 0.069 |  | 67 |  | 2.000 |  | 1.000 |  |
|  |  | Term, ENR, Last session, FAs |  | 0.166 |  | 0.065 |  | 67 |  | 2.553 |  | 0.479 |  |
|  |  | Preterm, STD, Last session, FAs |  | 0.068 |  | 0.057 |  | 67 |  | 1.196 |  | 1.000 |  |
|  |  | Term, STD, Last session, FAs |  | 0.173 |  | 0.059 |  | 67 |  | 2.925 |  | 0.178 |  |
| Preterm, STD, First session, Hits |  | Term, STD, First session, Hits |  | -0.070 |  | 0.040 |  | 67 |  | -1.756 |  | 1.000 |  |
|  |  | Preterm, ENR, Last session, Hits |  | 0.016 |  | 0.036 |  | 67 |  | 0.453 |  | 1.000 |  |
|  |  | Term, ENR, Last session, Hits |  | -0.115 |  | 0.037 |  | 67 |  | -3.061 |  | 0.124 |  |
|  |  | Preterm, STD, Last session, Hits |  | -0.116 |  | 0.028 |  | 67 |  | -4.137 |  | 0.005 | ** |
|  |  | Term, STD, Last session, Hits |  | -0.123 |  | 0.031 |  | 67 |  | -3.936 |  | 0.009 | ** |
|  |  | Preterm, ENR, First session, FAs |  | 0.271 |  | 0.066 |  | 67 |  | 4.084 |  | 0.006 | ** |
|  |  | Term, ENR, First session, FAs |  | 0.405 |  | 0.072 |  | 67 |  | 5.591 |  | 3.188×10^-5^ | *** |
|  |  | Preterm, STD, First session, FAs |  | 0.104 |  | 0.029 |  | 67 |  | 3.612 |  | 0.024 | * |
|  |  | Term, STD, First session, FAs |  | 0.086 |  | 0.051 |  | 67 |  | 1.699 |  | 1.000 |  |
|  |  | Preterm, ENR, Last session, FAs |  | 0.506 |  | 0.057 |  | 67 |  | 8.896 |  | 5.639×10^-11^ | *** |
|  |  | Term, ENR, Last session, FAs |  | 0.534 |  | 0.062 |  | 67 |  | 8.662 |  | 1.451×10^-10^ | *** |
|  |  | Preterm, STD, Last session, FAs |  | 0.436 |  | 0.036 |  | 67 |  | 12.010 |  | 2.542×10^-16^ | *** |
|  |  | Term, STD, Last session, FAs |  | 0.541 |  | 0.044 |  | 67 |  | 12.206 |  | 1.224×10^-16^ | *** |
| Term, STD, First session, Hits |  | Preterm, ENR, Last session, Hits |  | 0.087 |  | 0.039 |  | 67 |  | 2.240 |  | 0.880 |  |
|  |  | Term, ENR, Last session, Hits |  | -0.044 |  | 0.040 |  | 67 |  | -1.097 |  | 1.000 |  |
|  |  | Preterm, STD, Last session, Hits |  | -0.045 |  | 0.034 |  | 67 |  | -1.343 |  | 1.000 |  |
|  |  | Term, STD, Last session, Hits |  | -0.053 |  | 0.032 |  | 67 |  | -1.653 |  | 1.000 |  |
|  |  | Preterm, ENR, First session, FAs |  | 0.342 |  | 0.068 |  | 67 |  | 5.020 |  | 2.680×10^-4^ | *** |
|  |  | Term, ENR, First session, FAs |  | 0.475 |  | 0.074 |  | 67 |  | 6.430 |  | 1.251×10^-6^ | *** |
|  |  | Preterm, STD, First session, FAs |  | 0.175 |  | 0.048 |  | 67 |  | 3.620 |  | 0.024 | * |
|  |  | Term, STD, First session, FAs |  | 0.156 |  | 0.033 |  | 67 |  | 4.722 |  | 7.672×10^-4^ | *** |
|  |  | Preterm, ENR, Last session, FAs |  | 0.577 |  | 0.059 |  | 67 |  | 9.805 |  | 1.444×10^-12^ | *** |
|  |  | Term, ENR, Last session, FAs |  | 0.605 |  | 0.063 |  | 67 |  | 9.531 |  | 4.231×10^-12^ | *** |
|  |  | Preterm, STD, Last session, FAs |  | 0.506 |  | 0.043 |  | 67 |  | 11.673 |  | 9.098×10^-16^ | *** |
|  |  | Term, STD, Last session, FAs |  | 0.612 |  | 0.042 |  | 67 |  | 14.687 |  | 1.441×10^-20^ | *** |
| Preterm, ENR, Last session, Hits |  | Term, ENR, Last session, Hits |  | -0.131 |  | 0.036 |  | 67 |  | -3.645 |  | 0.023 | * |
|  |  | Preterm, STD, Last session, Hits |  | -0.132 |  | 0.028 |  | 67 |  | -4.664 |  | 9.337×10^-4^ | *** |
|  |  | Term, STD, Last session, Hits |  | -0.140 |  | 0.029 |  | 67 |  | -4.737 |  | 7.371×10^-4^ | *** |
|  |  | Preterm, ENR, First session, FAs |  | 0.255 |  | 0.064 |  | 67 |  | 3.958 |  | 0.009 | ** |
|  |  | Term, ENR, First session, FAs |  | 0.389 |  | 0.072 |  | 67 |  | 5.427 |  | 5.705×10^-5^ | *** |
|  |  | Preterm, STD, First session, FAs |  | 0.088 |  | 0.045 |  | 67 |  | 1.971 |  | 1.000 |  |
|  |  | Term, STD, First session, FAs |  | 0.070 |  | 0.049 |  | 67 |  | 1.411 |  | 1.000 |  |
|  |  | Preterm, ENR, Last session, FAs |  | 0.490 |  | 0.054 |  | 67 |  | 9.035 |  | 3.216×10^-11^ | *** |
|  |  | Term, ENR, Last session, FAs |  | 0.518 |  | 0.061 |  | 67 |  | 8.529 |  | 2.489×10^-10^ | *** |
|  |  | Preterm, STD, Last session, FAs |  | 0.420 |  | 0.039 |  | 67 |  | 10.674 |  | 4.394×10^-14^ | *** |
|  |  | Term, STD, Last session, FAs |  | 0.525 |  | 0.043 |  | 67 |  | 12.203 |  | 1.231×10^-16^ | *** |
| Term, ENR, Last session, Hits |  | Preterm, STD, Last session, Hits |  | -0.001 |  | 0.030 |  | 67 |  | -0.034 |  | 1.000 |  |
|  |  | Term, STD, Last session, Hits |  | -0.009 |  | 0.032 |  | 67 |  | -0.280 |  | 1.000 |  |
|  |  | Preterm, ENR, First session, FAs |  | 0.386 |  | 0.067 |  | 67 |  | 5.800 |  | 1.417×10^-5^ | *** |
|  |  | Term, ENR, First session, FAs |  | 0.519 |  | 0.071 |  | 67 |  | 7.288 |  | 3.924×10^-8^ | *** |
|  |  | Preterm, STD, First session, FAs |  | 0.219 |  | 0.046 |  | 67 |  | 4.752 |  | 7.066×10^-4^ | *** |
|  |  | Term, STD, First session, FAs |  | 0.201 |  | 0.051 |  | 67 |  | 3.957 |  | 0.009 | ** |
|  |  | Preterm, ENR, Last session, FAs |  | 0.621 |  | 0.057 |  | 67 |  | 10.887 |  | 1.927×10^-14^ | *** |
|  |  | Term, ENR, Last session, FAs |  | 0.649 |  | 0.060 |  | 67 |  | 10.820 |  | 2.481×10^-14^ | *** |
|  |  | Preterm, STD, Last session, FAs |  | 0.550 |  | 0.041 |  | 67 |  | 13.453 |  | 1.176×10^-18^ | *** |
|  |  | Term, STD, Last session, FAs |  | 0.656 |  | 0.045 |  | 67 |  | 14.739 |  | 1.214×10^-20^ | *** |
| Preterm, STD, Last session, Hits |  | Term, STD, Last session, Hits |  | -0.008 |  | 0.023 |  | 67 |  | -0.346 |  | 1.000 |  |
|  |  | Preterm, ENR, First session, FAs |  | 0.387 |  | 0.063 |  | 67 |  | 6.166 |  | 3.370×10^-6^ | *** |
|  |  | Term, ENR, First session, FAs |  | 0.520 |  | 0.069 |  | 67 |  | 7.539 |  | 1.453×10^-8^ | *** |
|  |  | Preterm, STD, First session, FAs |  | 0.220 |  | 0.040 |  | 67 |  | 5.537 |  | 3.888×10^-5^ | *** |
|  |  | Term, STD, First session, FAs |  | 0.202 |  | 0.046 |  | 67 |  | 4.420 |  | 0.002 | ** |
|  |  | Preterm, ENR, Last session, FAs |  | 0.622 |  | 0.053 |  | 67 |  | 11.829 |  | 5.060×10^-16^ | *** |
|  |  | Term, ENR, Last session, FAs |  | 0.650 |  | 0.058 |  | 67 |  | 11.263 |  | 4.405×10^-15^ | *** |
|  |  | Preterm, STD, Last session, FAs |  | 0.551 |  | 0.033 |  | 67 |  | 16.504 |  | 3.167×10^-23^ | *** |
|  |  | Term, STD, Last session, FAs |  | 0.657 |  | 0.039 |  | 67 |  | 17.007 |  | 6.255×10^-24^ | *** |
| Term, STD, Last session, Hits |  | Preterm, ENR, First session, FAs |  | 0.395 |  | 0.063 |  | 67 |  | 6.235 |  | 2.584×10^-6^ | *** |
|  |  | Term, ENR, First session, FAs |  | 0.528 |  | 0.070 |  | 67 |  | 7.596 |  | 1.158×10^-8^ | *** |
|  |  | Preterm, STD, First session, FAs |  | 0.228 |  | 0.041 |  | 67 |  | 5.520 |  | 4.097×10^-5^ | *** |
|  |  | Term, STD, First session, FAs |  | 0.209 |  | 0.046 |  | 67 |  | 4.593 |  | 0.001 | ** |
|  |  | Preterm, ENR, Last session, FAs |  | 0.630 |  | 0.053 |  | 67 |  | 11.829 |  | 5.060×10^-16^ | *** |
|  |  | Term, ENR, Last session, FAs |  | 0.658 |  | 0.058 |  | 67 |  | 11.281 |  | 4.155×10^-15^ | *** |
|  |  | Preterm, STD, Last session, FAs |  | 0.559 |  | 0.035 |  | 67 |  | 15.785 |  | 3.404×10^-22^ | *** |
|  |  | Term, STD, Last session, FAs |  | 0.665 |  | 0.038 |  | 67 |  | 17.331 |  | 2.248×10^-24^ | *** |
| Preterm, ENR, First session, FAs |  | Term, ENR, First session, FAs |  | 0.133 |  | 0.091 |  | 67 |  | 1.468 |  | 1.000 |  |
|  |  | Preterm, STD, First session, FAs |  | -0.167 |  | 0.072 |  | 67 |  | -2.334 |  | 0.723 |  |
|  |  | Term, STD, First session, FAs |  | -0.185 |  | 0.075 |  | 67 |  | -2.483 |  | 0.560 |  |
|  |  | Preterm, ENR, Last session, FAs |  | 0.235 |  | 0.073 |  | 67 |  | 3.209 |  | 0.084 |  |
|  |  | Term, ENR, Last session, FAs |  | 0.263 |  | 0.083 |  | 67 |  | 3.183 |  | 0.088 |  |
|  |  | Preterm, STD, Last session, FAs |  | 0.164 |  | 0.068 |  | 67 |  | 2.403 |  | 0.666 |  |
|  |  | Term, STD, Last session, FAs |  | 0.270 |  | 0.071 |  | 67 |  | 3.822 |  | 0.013 | * |
| Term, ENR, First session, FAs |  | Preterm, STD, First session, FAs |  | -0.301 |  | 0.077 |  | 67 |  | -3.895 |  | 0.011 | * |
|  |  | Term, STD, First session, FAs |  | -0.319 |  | 0.080 |  | 67 |  | -3.984 |  | 0.008 | ** |
|  |  | Preterm, ENR, Last session, FAs |  | 0.102 |  | 0.084 |  | 67 |  | 1.207 |  | 1.000 |  |
|  |  | Term, ENR, Last session, FAs |  | 0.130 |  | 0.081 |  | 67 |  | 1.601 |  | 1.000 |  |
|  |  | Preterm, STD, Last session, FAs |  | 0.031 |  | 0.074 |  | 67 |  | 0.418 |  | 1.000 |  |
|  |  | Term, STD, Last session, FAs |  | 0.137 |  | 0.076 |  | 67 |  | 1.791 |  | 1.000 |  |
| Preterm, STD, First session, FAs |  | Term, STD, First session, FAs |  | -0.018 |  | 0.057 |  | 67 |  | -0.320 |  | 1.000 |  |
|  |  | Preterm, ENR, Last session, FAs |  | 0.402 |  | 0.063 |  | 67 |  | 6.395 |  | 1.407×10^-6^ | *** |
|  |  | Term, ENR, Last session, FAs |  | 0.430 |  | 0.067 |  | 67 |  | 6.398 |  | 1.407×10^-6^ | *** |
|  |  | Preterm, STD, Last session, FAs |  | 0.332 |  | 0.045 |  | 67 |  | 7.352 |  | 3.079×10^-8^ | *** |
|  |  | Term, STD, Last session, FAs |  | 0.437 |  | 0.052 |  | 67 |  | 8.441 |  | 3.558×10^-10^ | *** |
| Term, STD, First session, FAs |  | Preterm, ENR, Last session, FAs |  | 0.421 |  | 0.066 |  | 67 |  | 6.336 |  | 1.744×10^-6^ | *** |
|  |  | Term, ENR, Last session, FAs |  | 0.449 |  | 0.071 |  | 67 |  | 6.361 |  | 1.591×10^-6^ | *** |
|  |  | Preterm, STD, Last session, FAs |  | 0.350 |  | 0.053 |  | 67 |  | 6.583 |  | 6.865×10^-7^ | *** |
|  |  | Term, STD, Last session, FAs |  | 0.456 |  | 0.052 |  | 67 |  | 8.795 |  | 8.465×10^-11^ | *** |
| Preterm, ENR, Last session, FAs |  | Term, ENR, Last session, FAs |  | 0.028 |  | 0.075 |  | 67 |  | 0.372 |  | 1.000 |  |
|  |  | Preterm, STD, Last session, FAs |  | -0.071 |  | 0.059 |  | 67 |  | -1.191 |  | 1.000 |  |
|  |  | Term, STD, Last session, FAs |  | 0.035 |  | 0.062 |  | 67 |  | 0.567 |  | 1.000 |  |
| Term, ENR, Last session, FAs |  | Preterm, STD, Last session, FAs |  | -0.099 |  | 0.064 |  | 67 |  | -1.544 |  | 1.000 |  |
|  |  | Term, STD, Last session, FAs |  | 0.007 |  | 0.066 |  | 67 |  | 0.106 |  | 1.000 |  |
| Preterm, STD, Last session, FAs |  | Term, STD, Last session, FAs |  | 0.106 |  | 0.047 |  | 67 |  | 2.232 |  | 0.880 |  |
| * p < .05, ** p < .01, *** p < .001 | | | | | | | | | | | | | |
| *Note.*  P-value adjusted for comparing a family of 120 estimates. | | | | | | | | | | | | | |
